# Histone binding of ASF1 is required for fruiting body development, but not for genome stability in the filamentous fungus *Sordaria macrospora*

**DOI:** 10.1101/2023.06.16.545311

**Authors:** Jan Breuer, Tobias Busche, Jörn Kalinowski, Minou Nowrousian

**Affiliations:** Department of Molecular and Cellular Botany, Ruhr University Bochum, 44801 Bochum, Germany; Center for Biotechnology, Bielefeld University, 33594 Bielefeld, Germany; Medical School OWL, Bielefeld University, 33615 Bielefeld, Germany

## Abstract

The highly conserved eukaryotic histone chaperone ASF1 is involved in the assembly and disassembly of nucleosomes during transcription, DNA replication and repair. It was the first chaperone discovered to be involved in all three of these processes. The filamentous fungus *Sordaria macrospora* is one of only two multicellular organisms where *asf1* deletions are viable, which makes it useful for *in vivo* analysis of this central regulator of eukaryotic chromatin structure. Deletion of *asf1* in *S. macrospora* leads to sterility, a reduction of DNA methylation, and upregulation of genes that are usually weakly expressed in the wild type. Here, we focused on the functions of the highly conserved core and the divergent C-terminal tail of ASF1, studied the effects of ASF1 on histone modifications and tested its relevance for genomic stability. By Co-IP and complementation analysis we showed that substitutions of amino acid V94 or truncations of the C-terminal tail abolish histone binding and do not complement the sterile mutant phenotype. Δasf1 is sensitive to the DNA damaging agent MMS, while complementation strains, even those with non-histone-binding variants, regain wild type-like resistance. To aid in subsequent ChIP-seq and Hi-C analyses, we generated a chromosome-resolved genome assembly of *S. macrospora*. ChIP-Seq analysis revealed a global increase of H3K27me3 in Δasf1, accompanied by a global decrease of H3K56ac. By using Hi-C we detected a tandem duplication of around 600 kb on chromosome 2 in the mutant. Crossing experiments indicated linkage between the viability of Δasf1 strains and the presence of the duplication.

**Importance:** Histone chaperones are proteins that are involved in nucleosome assembly and disassembly and can therefore influence all DNA-dependent processes including transcription, DNA replication and repair. ASF1 is a histone chaperone that is conserved throughout eukaryotes. In contrast to most other multicellular organisms, a deletion mutant of *asf1* in the fungus *Sordaria macrospora* is viable; however, the mutant is sterile. In this study, we could show that the histone binding ability of ASF1 is required for fertility in *S. macrospora*, whereas the function of ASF1 in maintenance of genome stability does not require histone binding. We also showed that the histone modifications H3K27me3 and H3K56ac are misregulated in the Δasf1 mutant. Furthermore, we identified a large duplication on chromosome 2 of the mutant strain that is genetically linked to the Δasf1 allele present on chromosome 6, suggesting that viability of the mutant might depend on the presence of the duplicated region.

## Introduction

Multicellular development is a complex process that evolved independently in multiple branches of eukaryotes (1). In fungi, multicellular development evolved at least twice and possibly more times, the most prominent examples being the fruiting bodies of filamentous ascomycetes and basidiomycetes (2). To study the genetic background of multicellular development, simple, easy to handle model organisms are of particular interest. Several species of fungi have proven to be usable models for the analysis of developmental processes, and the generation of complex, multicellular fruiting bodies of filamentous ascomycetes is an excellent example (3). *Sordaria macrospora*, a homothallic ascomycete that can form its perithecia in 7 days under laboratory conditions, has been used as a model to study fungal sexual development and multicellular development in general. During the formation of fruiting bodies in *S. macrospora*, at least 13 different cell types are generated, many of which cannot be found in vegetative mycelium (4). Sexual development and perithecia formation are accompanied by transcriptional changes, and transcriptomics can be a helpful tool to identify important developmental genes by reverse genetic approaches. A number of transcription factors and chromatin modifiers have been identified this way and were proven to be fundamental for multicellular development (5). Among them is the conserved histone chaperone ASF1 that was shown to be upregulated during sexual development and is essential for this process in *S. macrospora* (6). ASF1 is involved in the assembly and disassembly of nucleosomes during transcription, replication and DNA-repair and was the first chaperone found to be involved in all three of these processes. It mediates nucleosome assembly and disassembly in promoter regions during transcriptional activation and elongation in *Saccharomyces cerevisiae* (7). ASF1 is also important for histone modification by interacting with enzymes that mediate the corresponding modifications. For example, it has been shown to be involved in the acetylation of histone H3 at lysines 9 and 56 and also to play a role in the genome-wide deacetylation of histones (8, 9). The methylation of histones also appears to proceed in conjunction with ASF1. In *Mus musculus*, an altered distribution of H3K9me3 and H3K27me3 was shown under ASF1 deficiency. ASF1 was also proven to be important for histone recycling (10). Additionally, ASF1-depleted cells were shown to exhibit higher tendencies for double strand breaks, showcasing a role of ASF1 during the maintenance of genomic stability (11). The function and structure of ASF1 is highly conserved in all eukaryotes. The 155 amino acids of the N-terminus of ASF1 form a globular core with conserved acidic patches that mediate interaction with the C-terminus of histone H3. ASF1 has been shown to interact with the histone H3-H4 heterodimer *in vitro* and *in vivo*. In addition to binding of H3, binding of the H4 C-terminus to ASF1 also occurs. ASF1 and the H3-H4 dimer can form a heterotrimeric complex, although both histones can be bound individually by ASF1 (7). The central role of ASF1 for eukaryotic chromatin structure is underscored by findings in human cell lines that showed all non-DNA bound histones H3 and H4 are bound to ASF1 (12, 13). Therefore, researching ASF1 functions during multicellular development might be fundamental for understanding the contribution of the chromatin landscape to development in fungi and multicellular eukaryotes in general. Gaining insight into the functions of ASF1 is hindered by the fact that deletions of the respective gene are lethal in almost all higher eukaryotes. Thus, research on ASF1 was mostly conducted with ASF1-depleted cell lines, *in vitro* experiments and studies of the unicellular *S. cerevisiae* where an *asf1* deletion is viable (14–16)*. S. macrospora* and *Arabidopsis thaliana* were shown to be exceptions in this regard and deletions are viable in these organisms (6, 17) (for *S. macrospora*, this might depend on the presence of a secondary mutation as described in the results). Therefore, the analysis of *S. macrospora* Δasf1 strains offers the unique opportunity to perform ASF1 research *in vivo* in a multicellular fungus. These strains were shown to be sterile, highlighting a central role of ASF1 during sexual development. Unexpectedly, the positioning of nucleosomes was not affected in *asf1* deletion strains. A significant reduction of DNA methylation was observed in *S. macrospora* Δasf1, and a large number of genes that are usually weakly expressed in wild type strains are upregulated in Δasf1, indicating a role of ASF1 during gene regulation and DNA methylation (18). The *S. macrospora* ASF1 has been shown to bind histones H3 and H4 (6), however, it has not yet been analyzed if the histone-binding activity is actually required for fruiting body formation. Therefore, in this study, we analyzed the histone binding function of ASF1 and its connection to sexual development. We performed interaction studies and complementation analysis to verify the importance of amino acid V94 (valine at position 94), a highly conserved histone interaction site known from *in vitro* studies (7, 19). The C-terminal region of ASF1 is much less conserved than the first 155 amino acids and parts of it do not exist in animal or plant models (Figure 1). We generated truncated versions of this region of *asf1* and used these to perform interaction studies and complementation analysis to study the relevance of the divergent and functionally less well studied C-terminus of ASF1. Additionally, we performed genomic stability assays based on growth under the influence of the genotoxic substance methyl methansulfonate (MMS). We further focused on the role of ASF1 in histone modification and chromatin structure by performing ChIP (chromatin immunoprecipitation)-seq and Hi-C (chromosome conformation capture with high-throughput sequencing) experiments with the wild type and the Δasf1 mutant. For improved analysis of the ChIP-seq and Hi-C data, we first generated a chromosome-level genome sequence of *S. macrospora*. Participation of ASF1 in histone acetylation and deacetylation has been documented in *S. cerevisiae*, and a function during histone methylation is regarded as highly probable and connections to the effect on DNA-methylation and gene regulation might be possible (10). Therefore, ChIP-seq was performed for H3K27me3 and H3K56ac. Furthermore, we performed Hi-C experiments to compare overall chromatin structure of young, vegetative mycelium to older, sexual mycelium and *asf1* deletion mutants.

**Figure 1.**
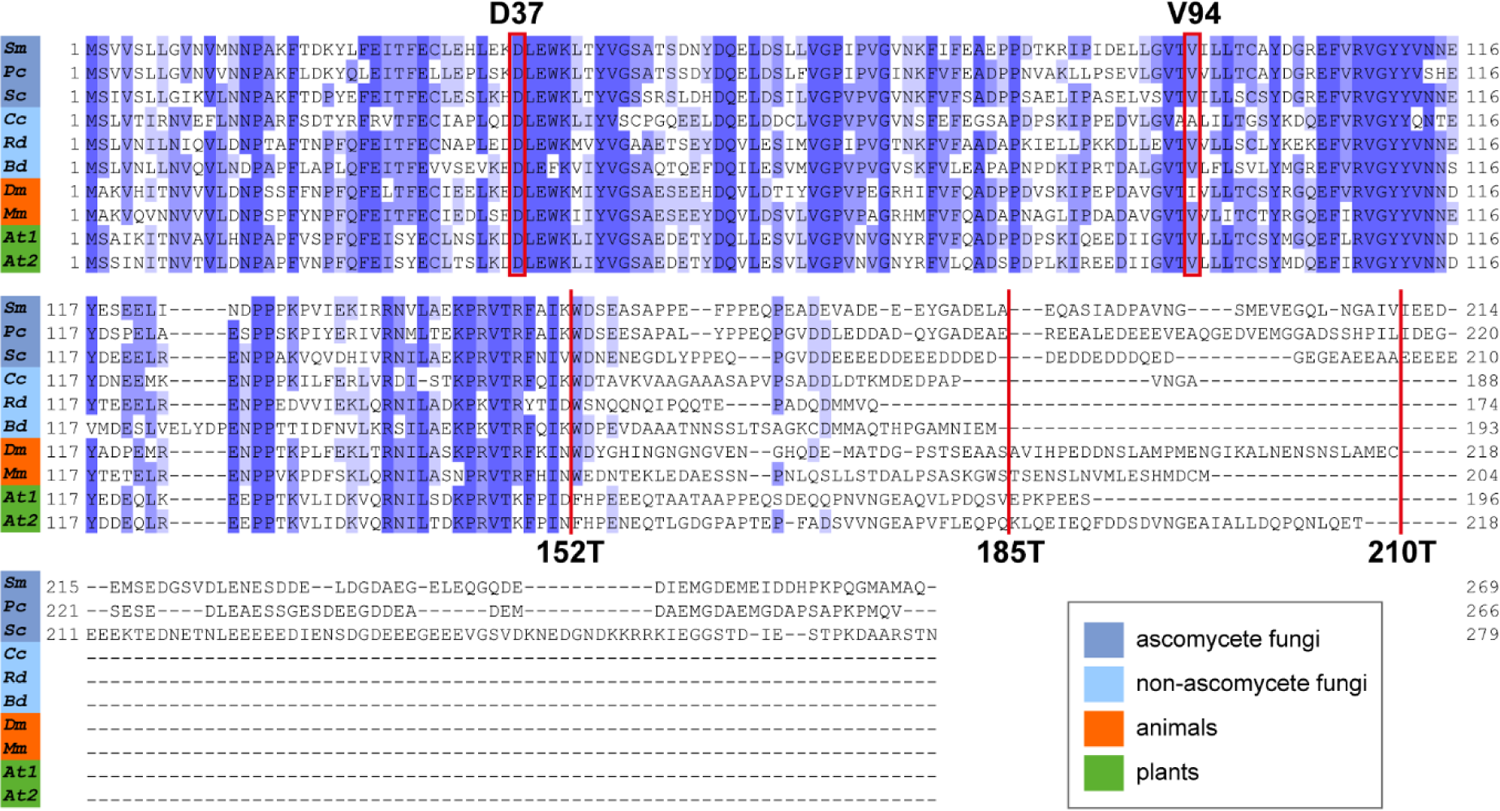
Alignment of ASF1 amino acid sequences from fungi, animals and plant. The first 152 amino acids are highly conserved. The C-terminal region is more divergent and parts of it exist only in ascomycete fungi. Positions D37 and V94 were substituted for functional analysis in this study. D37 is known as a HIRA interaction site in *Schizosaccharomyces pombe* (57), while V94 has been established as the H3-H4 interaction site (7, 19). Positions 152, 185 and 210 were targeted to create truncated ASF1 variants. Abbreviations and accession numbers: Ascomycetes: *Sm*, *Sordaria macrospora* (KAA8632619.1); *Pc*, *Pyronema confluens* (ADZ55332.1); *Sc*, *Saccharomyces cerevisiae* (NP_012420.1). Basidiomycetes: *Cc*, *Coprinopsis cinerea* (XP_001834011.1). Mucoromycetes: *Rd*, *Rhizopus delemar* (EIE83079.1). Chytridiomycetes: *Bd*, *Batrachochytrium dendrobatidis* (KAJ8331661.1). Animals: *Dm*, *Drosophila melanogaster* (NP_524163.1); *Mm*, *Mus musculus* (AAH27628.1). Plants: *At*, *Arabidopsis thaliana* (NP_176846.1 and NP_198627.1). The taxonomic groups are color coded according to the legend in the figure.

## Materials and Methods

### Strains, crosses and growth conditions

All strains used in this study have been summarized in Supplemental Table 1 and were cultivated on either solid or liquid cornmeal medium (BMM) or complete medium (CM) at 25 °C as previously described (20, 21). To perform genetic crosses, the spore color mutant fus was used as a partner to allow the identification of recombinant asci (22). Transformation of *S. macrospora* was performed as previously described (21).

### Cloning procedures, oligonucleotides and plasmids

Plasmids encoding ASF1 variants with amino acid substitutions for complementation and interaction analysis were generated by PCR mutagenesis and yeast recombinant cloning (23). Golden Gate cloning was used to assemble plasmids encoding truncated ASF1 variants (24). Oligonucleotides for the generation of plasmids and integration tests have been listed in Supplemental Table 2. All plasmids used in this study are listed in Supplemental Table 3.

### Histone interaction assay by Co-Immunoprecipitation (Co-Ip)

Plasmids coding for eGFP-tagged ASF1 variants were co-transformed together with plasmids encoding FLAG-tagged histones H3 or H4 into the *S. macrospora* wild type strain SN1693 by protoplast transformation (25). The resulting strains were grown for 4 d at 27 °C in 20 ml liquid BMM medium with 50 µg / ml Nourseothricin. The mycelium was harvested by filtration and frozen in liquid nitrogen before being ground into powder and mixed with protein extraction buffer (26). Co-Immunoprecipitation was performed according to the manufacturers’ instructions using GFP-Trap® Beads (Chromotek) and ANTI-FLAG® M2 Beads (Sigma Aldrich). Raw and elution fractions were used for SDS-PAGE (27) before being transferred to a PVDF membrane by western blotting. Co-Ip results were generated with two biological replicates each for eGFP trapping and FLAG trapping. Signals for eGFP and FLAG were detected by using Mouse Anti-GFP JL-8 (Clontech), Mouse Anti-Flag (Sigma Aldrich) and Goat Anti-Mouse, HRP-linked (Cell Signaling) antibodies according to the manufacturers’ instructions.

### Complementation of *S. macrospora* with ASF1 variants and MMS tests

Plasmids for the expression of ASF1 variants were generated from wild type genomic DNA with specific primers amplifying *asf1* with specific substitutions or a shortened C-terminus and cloned by Golden Gate Cloning (24) or via yeast homologous recombination (28). The *asf1* deletion strain SN1983 was transformed with the respective plasmids by protoplast transformation resulting in ectopic integration of the plasmid (25). Complementation strains that did not gain fertility on their own were crossed against the wild type, and ascospores were isolated to obtain homokaryotic strains, while fertile transformants were used to isolate ascospores directly (22). Phenotypical characterization was performed by documenting the growth and fertility on BMM plates with a Stemi 2000-C stereomicroscope (Zeiss). To evaluate the resistance against methyl methanesulfonate (MMS) all strains were inoculated on BMM plates with 0.007 % MMS and growth was observed over a span of 7d. The validity of the MMS assay was tested by checking the persistence of the effect of MMS on *asf1* deletion mutants on 3 days old plates over a period of 7 days.

### Microscopic analysis

For analysis of the subcellular localization of ASF1-EGFP fusion proteins, strains were grown on glass slides together with strains expressing H3-MRFP fusion proteins as described (29). Light and fluorescence microscopy was performed with an AxioImager microscope (Zeiss, Jena, Germany) with a Photometrix Cool SnapHQ camera (Roper Scientific). EGFP fluorescence was detected with Chroma (Bellows Falls, VT, USA) filter set 41017 (HQ470/40, HQ525/50, Q495lp) and mRFP fluorescence was detected with set 49008 (EG560/40x, ET630/75 m, T585lp). Images were further handled by using MetaMorph (Molecular Devices).

### Extraction and sequencing of genomic DNA from *S. macrospora*

For DNA preparation for nanopore sequencing, the wild type strain SN1693 was grown for 2 d at 27 °C in 20 ml liquid CM medium (25) in petri dishes, harvested by filtration and ground in liquid nitrogen. 500 µg of ground mycelium was used for extraction of high molecular weight DNA with the NucleoBond HMW-Kit (Macherey-Nagel, Düren, Germany) according to the manufacturer’s instructions. A sequencing library with wild type genomic DNA was prepared using the Nanopore DNA Ligation Sequencing Kit (SQK-LSK109, Oxford Nanopore Technologies, Oxford, UK) according to the manufacturer’s instructions. Sequencing was performed on an Oxford Nanopore GridION Mk1 sequencer using an R9.4.1 flow cell, which was prepared according to the manufacturer’s instructions. Basecalling was performed using guppy v5.0.11 with the super-accurate basecalling model. A total of 290927 reads were obtained with an average read length of 15874 bp and an N50 of 23018 bp.

For DNA preparation for Illumina sequencing of the Δasf1 mutant, the Δasf1 strain SN1983 was grown for 3 d at 27 °C in 20 ml liquid CM medium in petri dishes, harvested by filtration and ground in liquid nitrogen. DNA preparation was performed as described previously for *Pyronema confluens* (30). Library preparation and Illumina sequencing of a 300 bp insert library (150 bp paired-end sequencing) were performed at Novogene (Cambridge, UK).

### Genome assembly and annotation

Nanopore reads were assembled with SMARTdenovo (available at https://github.com/ruanjue/smartdenovo, accessed on 2021/06/29) resulting in nine contigs with a total size of 39.4 Mb. The initial assembly was subjected to four rounds of correction with Racon (31) using the nanopore reads for correction, and subsequently three rounds of correction with pilon (v1.24) (32) based on previously generated *S. macrospora* genomic Illumina reads (33) (NCBI SRA database, accession number SRR5749461). BLAST comparisons (34) with a previously generated *S. macrospora* genome assembly (35) showed that the smallest contig (161 kb) most likely contained mitochondrial DNA. This contig was removed, and the remaining eight contigs were analyzed for putative telomeric repeats (sequence TTAGGG) using a custom-made perl program. Telomeric repeats were found at both ends of five contigs, and at one end each in the remaining three contigs. Of the latter, one contig contained the rDNA repeats at the end without telomeric repeats. The other two non-telomeric contig ends were compared to each other and found to have an overlap of 2.2 kb with 100 % sequence identity, making it likely that these contigs were part of the same chromosome. The contigs were fused manually, and the junction was checked for continuity by mapping the nanopore reads to the assembly with graphmap (36). The resulting assembly consists of 7 contigs with a total size of 39.4 Mb and an N50 of 5.7 Mb. Comparison to the genome assembly of the closely related ascomycete *Neurospora crassa* (FungiDB version 52) (37, 38) was performed with nucmer from the MUMmer package (v4.0.0rc1) (39).

Genome annotation of protein-coding genes was performed with MAKER (v3.01.03) (40) based on transcripts and proteins from a previous annotation of the *S. macrospora* genome (33) as well as on the *N. crassa* proteins (38) from FungiDB version 52 (37). tRNAs were annotated with tRNAscan-SE (v2.0.8) (41). rDNA repeats were annotated manually based on BLAST comparisons. *S. macrospora* locus tags from a previous annotation (33) were mapped onto the predicted gene models based on BLAST results. Approximately 1200 gene models were manually corrected in the genome browser Artemis (42) based on RNA-Seq data generated previously (18, 43) (NCBI GEO accession numbers GSE33668 and GSE92337) that were mapped to the assembly with Hisat2 (v2.2.1) (44).

### Western blot analysis of histone modifications and chromatin immunoprecipitation

Potential targets for ChIP-seq experiments were selected based on semiquantitative western blot screenings with antibodies against histone modifications previously suspected to be influenced by ASF1. The wild type strain SN1693, the Δasf1 strain SN1983, and complementation strains expressing *asf1* variants with amino acid substitutions or a truncated C-terminal region, were cultivated in liquid CM medium (25) for 3d and 4d, respectively, at 27 °C, harvested by filtration and washed with an isotonic buffer established for protoplasts (PPP) (21). Samples were frozen in liquid nitrogen and ground into a fine powder. Proteins were extracted by mixing the powder with extraction buffer (50 mM Tris/HCl pH 7.5, 250 mM NaCl, 0.05 % NP-40, 0.05 % β-mercaptoethanol, 0,3 % (v / v) Protease Inhibitor Cocktail Set IV (Calbiochem)) and 20 min centrifugation. Protein concentrations were determined by Bradford assays (45) and equal amounts were loaded and separated by SDS gelelectrophoresis and transferred to a PVDF membrane by western blotting. Antibodies against target histone modifications were used to detect respective bands (Anti H3K27Me3 #9733 Cell Signaling, Anti H3K9Me #ABE101 Merck Millipore, Anti H3K56Ac #61061 Active Motif, Anti H3K9Ac #39038 Active Motif, Anti H3 #9715 Cell Signaling), detection protocols have been summarized in Supplemental Table 4. Band strengths were compared by using a ChemiDoc XRS+ system (BioRad) with Image Lab 4.0 software (BioRad). Normalization for the total amounts of protein in the samples was based on Coomassie-stained gels run in parallel. The level of histone H3 was assessed as an internal control. Modifications that exhibited significant differences between wild type and deletion mutant were selected for ChIP-seq experiments. ChIP-samples were cultivated similar to the samples for western blotting and crosslinking was performed in PPP buffer with 1% formaldehyde for 15 min at room temperature, followed by a quenching time of 15 min by an excess amount of glycine. After three PPP washing steps, lysis was conducted in extraction buffer (50 mM Tris/HCl pH 7.5, 250 mM NaCl, 0.05 % NP-40, 0.05 % β-mercaptoethanol, 0,3 % (v/v) Protease Inhibitor Cocktail Set IV (Calbiochem)) for 15 min on ice by frequent vortexing. After centrifugation, chromatin solution was obtained and mixed with MNase buffer (18) and proteinase inhibitors. Chromatin digestion was conducted by incubation with 200 U/ml MNase (Thermo Fisher Scientific) for 105 min at 37 °C under constant shaking and stopped by adding EDTA to a concentration of 20 mM. Aliquots were removed as input samples, and immunoprecipitation performed over night at 4 °C on a spinning wheel by adding 10 µl of the respective antibody (Anti H3K27Me3 #9733 Cell Signaling, Anti H3 #2650 Cell Signaling, Anti H3K56Ac #61061 Active Motif). Afterwards, 4 h of incubation on Protein-A agarose beads (Santa Cruz) were followed by 2 washing steps in low salt (50 mM Tris-HCL pH 7.5, 10 mM EDTA, 10 mM NaCl) and 2 washing steps in high salt buffer (50 mM Tris-HCL pH 7.5, 10 mM EDTA, 500 mM NaCl). Each washing step was performed for 5 min at 4 °C on a spinning wheel and followed by 2 min of centrifugation at 2500 rpm to separate the beads from the buffer (46). Chromatin was washed off the beads by 2 h of incubation in TE buffer with 1 % SDS at 65 °C and de-crosslinked over night at 65 °C after adding 2 mg/ml Proteinase K (Genaxxon bioscience). Standard phenol/chloroform extraction was performed to isolate DNA from the ChIP and input samples. Two biological replicates each were created for H3, H3K27me3 and H3K56ac. Fragment sizes were evaluated on a Bioanalyzer 2100 (Agilent) according to the requirements for ChIP-seq samples set by Novogene. Samples that contained at least 20 ng of fragments in the range of 150 to 500 bp were selected and further processed. Size selection for fragments between 150 and 500 bp and library generation was performed at Novogene (Cambridge, UK), as well as Illumina sequencing (150 bp paired-end) of ChIP and input samples. The detailed ChIP-seq protocol can be found in Supplemental Text 1.

### ChIP-seq peak detection

At least 30 million paired-end reads per sample were sequenced at Novogene (Cambridge, UK). The obtained reads were trimmed with fastp 0.23.2 (47) and mapped to the reference genome of *S. macrospora* SN1693 by using BWA-MEM 0.7.17 with default parameters (48). Deduplication and input corrected peak detection were performed for the samples of H3, which were subsequently used together with the native input samples as control for H3K27me3 and H3K56ac by using MACS2 callpeak 2.2.7.1 in paired end mode. For both histone marks, broad regions were identified with a 0.01 minimum FDR (false discovery rate) cutoff and an effective genome size of 39 Mb (49, 50). Differential peak analysis was conducted by discarding overlapping peaks in wildtype and Δasf1 using BEDTools 2.30.0 with a minimum overlap of 10 % (51). Peaks were annotated to the reference genome using HOMER annotatePeaks 4.9.1 (52).

### Hi-C sample preparation

Young wild type samples of SN1693 where cultivated for 2 d at 27 °C in liquid CM medium (25). Samples for the older wild type were grown for 5 d and Δasf1 SN1983 for 3 d under the same conditions. Mycelium was harvested by filtration, washed with PPP (21) and frozen in liquid nitrogen. The Omni-C library was prepared using the Dovetail® Omni-C® Kit according to the manufacturer’s protocol with additional changes to enable the use on *S. macrospora*. Briefly, the chromatin was fixed with 1 % formaldehyde for 10 min at room temperature on a spinning wheel and quenched by bringing the solution up to 200 mM glycine. The cross-linked chromatin was then digested in situ with DNase I according to the manufacturer’s instructions. Following digestion, the cells were lysed in 1 % SDS to extract the chromatin fragments and the chromatin fragments were bound to Chromatin Capture Beads. Next, the chromatin ends were repaired and ligated to a biotinylated bridge adapter followed by proximity ligation of adapter-containing ends by using the reagents supplied by the Dovetail® Omni-C® Kit. After proximity ligation, the crosslinks were reversed at 68 °C in Dovetail® Crosslink Reversal Buffer and 1 µg / µl Proteinase K overnight. The DNA was purified with SPRIselect™ Beads (Beckmann Coulter) and 150 ng were converted into a sequencing library using Illumina-compatible adaptors by using the Dovetail® Library Module for Illumina®. Biotin-containing fragments were isolated using streptavidin beads prior to PCR amplification. Illumina sequencing (150 bp paired-end) was performed by Novogene (Cambridge, UK).

### Generation of Hi-C maps

Hi-C maps were created by following the Dovetail® Omni-C® Pipeline with adjustments for the genome of *S. macrospora*. Reads were trimmed with fastp 0.23.2 (47) and mapped to the reference genome of *S. macrospora* SN1693 by using BWA-MEM 0.7.17 with default parameters (48). Valid ligation events were recorded by using pairtools 1.02 (53) parse with a min-map quality of 40 and a max-inter-align-gap of 30 before being sorted by pairtools sort. PCR duplicates were removed by using pairtools dedup and the pairs file was generated by pairtools split. The hic matrix was generated by using juicertools 1.6.2 (54) with standard parameters.

### Data availability

Nanopore reads and the chromosome-resolved genome sequence for the *S. macrospora* wild type strain SN1693 were deposited in the NCBI databases under BioProject ID PRJNA744349 (http://www.ncbi.nlm.nih.gov/bioproject/744349, SRA accession number SRR15901728 for the nanopore reads, DDBJ/ENA/GenBank accession numbers CP083903-CP083909 for the seven chromosomes). ChIP-seq and Hi-C data generated in this study were submitted to the NCBI GEO database as a superseries under accession number GSE216634 (https://www.ncbi.nlm.nih.gov/geo/query/acc.cgi?acc=GSE216634, accession numbers GSE216629 and GSE228529 for ChIP-seq data and GSE216632 for Hi-C data). Illumina reads from the Δasf1 strain were deposited in the NCBI SRA database under BioProject ID PRJNA922040 (https://www.ncbi.nlm.nih.gov/sra/PRJNA922040).

## Results

### A chromosome-level genome assembly of *S. macrospora*

The nuclear genome of *S. macrospora* was previously shown by pulsed-field gel electrophoresis to consist of seven chromosomes (21). Prior to our study, three genome assemblies of *S. macrospora* have been generated with increasing continuity (33, 35, 55); however, even the latest genome assembly still consists of 584 scaffolds (33) and would therefore be unsuited for analyses at chromosome level. Therefore, to aid in the planned Hi-C and ChIP-seq analyses, we generated a chromosome-level genome assembly of *S. macrospora* based on nanopore sequence reads. The new assembly (*S. macrospora* genome version 04) consists of seven gapless contigs with a total size of 39.4 Mb (Figure S1) which is similar in size to previous assemblies and to estimates by pulsed-field gel electrophoresis of 39.5 Mb (21, 33, 35, 55). Six of the seven contigs have telomeric repeats at both ends, whereas one contig has telomeric repeats at one end and 10 copies of the ribosomal RNA (rRNA) genes at the other end with the repetitive nature of the rRNA genes most likely preventing assembly of telomeric repeats at that end. This indicates that the seven contigs represent the seven chromosomes of *S. macrospora*. A comparison with the genome assembly of the closely related ascomycete *Neurospora crassa* showed a high degree of synteny for the seven chromosomes of each species, consistent with previous analyses of smaller genomic regions of the two fungi (35, 56) (Figure S1). Thus, a chromosome-level assembly for *S. macrospora* is now available and can be used for analyses at chromosome level.

### *S. macrospora* ASF1 depends on V94 and large parts of the C-terminus to accomplish histone binding

The function and structure of ASF1 is highly conserved in all eukaryotes and it has been described as a central histone chaperone that binds the histones H3 and H4 as a dimer and individually. The first 155 amino acids form a globular core in which acidic patches allow for the interaction with both histones (7). Especially the area surrounding position V94 has been shown to be essential for ASF1-histone binding in yeast (19). However, little is known about the functions of the C-terminal tail. This region is much more divergent than the globular core and parts of it exist in fungi, but not in animals or plants, with the region after amino acid 210 being restricted to ascomycetes (Figure 1). To analyze if V94 is required for histone binding of *S. macrospora* ASF1, we performed co-immunoprecipitation experiments with an ASF1 variant that includes a V94R substitution and a variant with a D37A substitution. The latter is a mutation of an interaction site with the histone chaperone HIRA that was identified in *S. pombe* (57). In addition, ASF1 variants with truncations after positions 152, 185 and 210 were generated to gain insight into possible involvement of the C-terminal region in the process of histone binding. In GFP- and FLAG-trap experiments, the V94R variant proved to be incapable of pulling down the histones H3 and H4, while the D37A variant performed like the wild type ASF1 (Figure 2, Supplemental Figures 2 and 3). Thus, the conserved V94 is required for histone interactions in *S. macrospora*, whereas the putative HIRA interaction site D37 is not. Truncated ASF1 variants were also unable to bind H3 and H4 when the truncations occurred after positions 152 or 185. Truncation after position 210 had no detectable effect and did not impair the capability to pull down the histones (Figure 3, Supplemental Figures 4 and 5).

**Figure 2.**
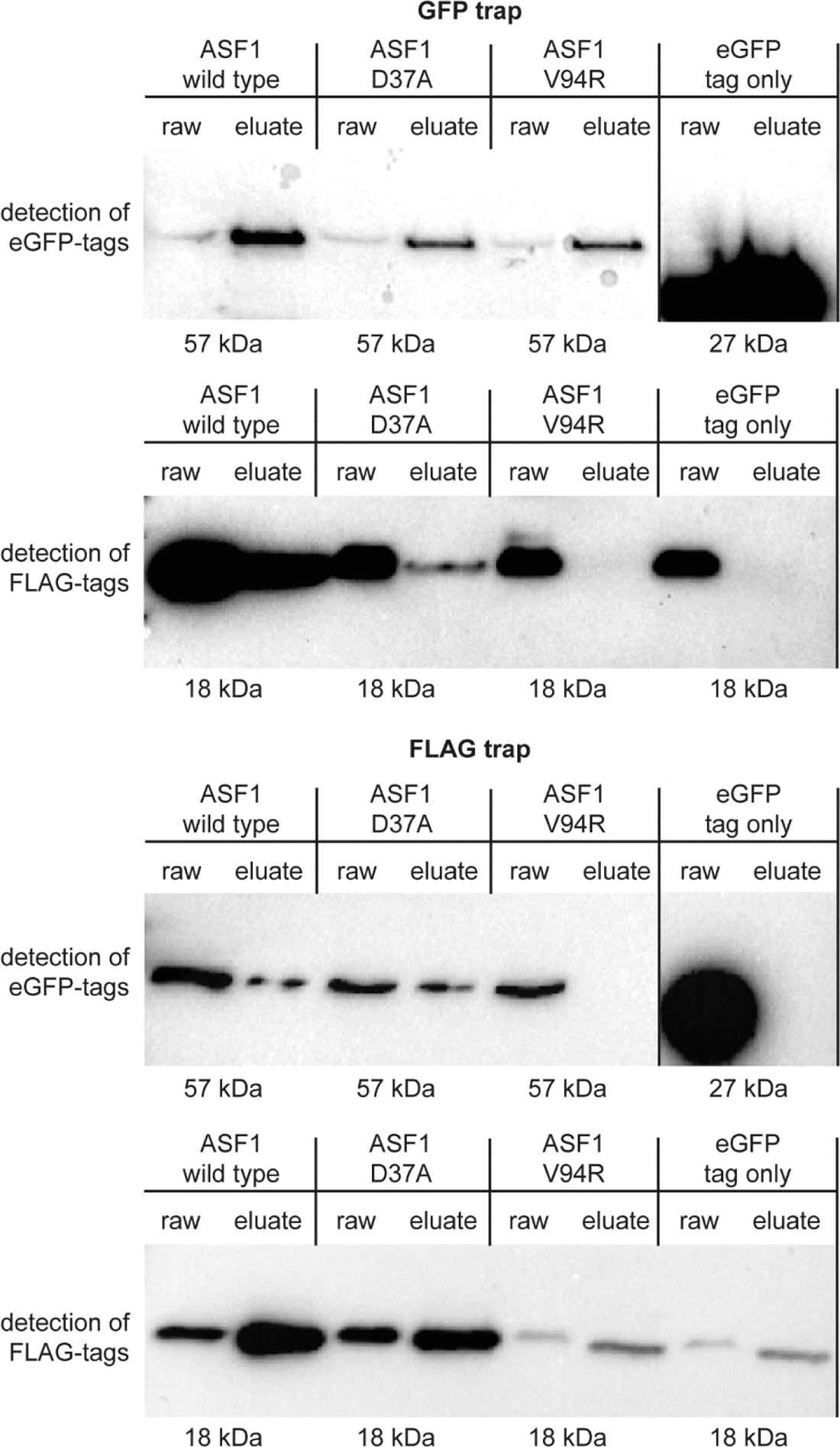
Co-immunoprecipitation results for ASF1 variants with amino acid substitutions and histone H3. GFP-tagged ASF1 wild type and variants D37A and V94R were used as potential interaction partners for Flag-tagged histone H3 in a GFP trap and a Flag trap. Results were checked by western blot analysis with antibodies against GFP and Flag tags. ASF1 wild type and the D37A variant showed signals for bait and prey proteins in the raw and eluate sample, indicating interaction, whereas the V94R variant showed the signal for the prey protein only in the raw sample. Strains expressing non-fused GFP with the corresponding Flag-tagged H3 were used as a negative control. Results for histone H4 are shown in Supplemental Figure 2. Uncropped blots are shown in Supplemental Figure 3A.

**Figure 3.**
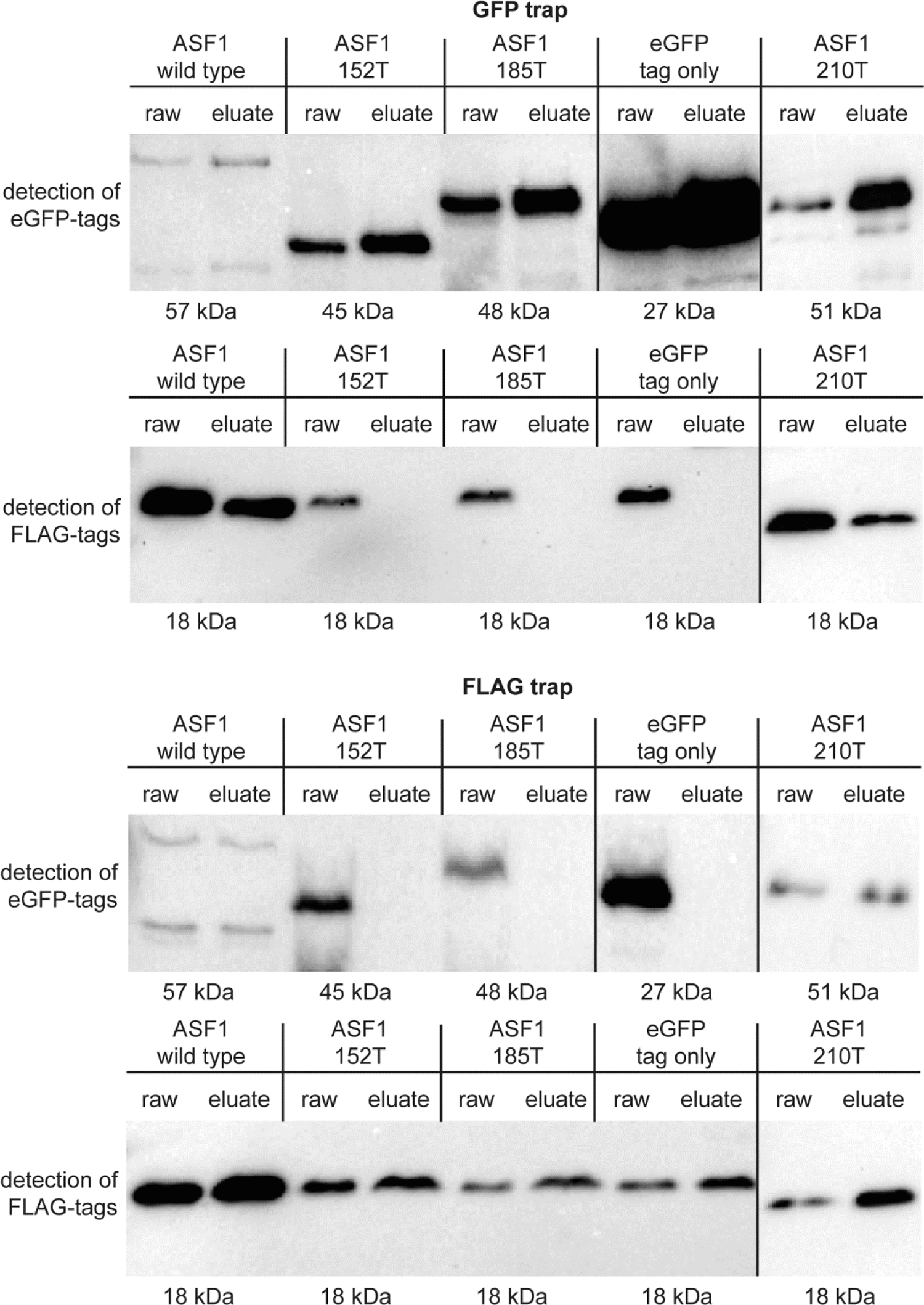
Co-immunoprecipitation results for truncated ASF1 variants and histone H3. GFP-tagged ASF1 wild type and variants 152T, 185T and 210T were used as potential interaction partners for Flag-tagged histone H3 in a GFP trap and a Flag trap. Results were checked by western blot analysis with antibodies against GFP and Flag tags. ASF1 wild type and the 210T variant showed signals for bait and prey proteins in the raw and eluate sample, indicating interaction, whereas the 152T and 185T variants showed the signal for the prey protein only in the raw sample. Strains expressing non-fused GFP with the corresponding Flag-tagged H3 were used as a negative control. Results for histone H4 are shown in Supplemental Figure 4. Uncropped blots are shown in Supplemental figure 5A.

### The histone binding function of ASF1 is essential for sexual development

We used the *asf1* variants described above carrying substitutions in the histone binding site V94, the putative HIRA interaction site D37 and variants with truncated C-terminal regions for complementation analysis in a previously constructed *asf1* deletion strain (6). By correlating the ability of these variants for rescuing the phenotype of the deletion mutant with information about their ability to bind histones, more insights into the role of ASF1-histone interaction during multicellular development can be gained. The *asf1* deletion mutant was transformed with plasmids coding for the respective variants, and homokaryotic strains were obtained by the isolation of ascospores from self-fertile perithecia or from genetic crosses. PCR screenings confirmed the integration of *asf1* variants in the resulting transformants (Supplemental Figure 6). The phenotype of the resulting strains was documented after growth for a week on BMM medium*. S. macrospora* Δasf1 is sterile and mostly consists only of vegetative mycelium, but is sometimes able to generate small protoperithecia. Introducing the wild type *asf1* gene back into the deletion mutant led to a complete reversal of the deletion phenotype and yielded strains indistinguishable from the wild type, gaining full fertility. Strains expressing the D37A variant also regained fertility, but showed slightly reduced complementation rates (Table 1). These strains were able to form perithecia, but exhibited slight aberrations with a tendency to grow aberrant perithecia and giant protoperithecia (Supplemental Figure 7). A reintroduction of an ASF1 variant with a V94R substitution did not complement the deletion mutant and the resulting strains exhibited the same phenotype as Δasf1 strains (Figure 4, Table 1). Since ASF1 V94R does not complement the sterile phenotype of deletion mutants and is unable to bind histones, a connection between histone binding and sexual development seems likely. ASF1 D37A proved to have no disruption in histone binding and regained fertility, but the substitution in a putative HIRA-binding site still had impact on the life cycle of *S. macrospora*, as evident by the formation of some aberrant perithecia. However, histone binding capability was enough to reach fertility. Truncations of ASF1 at positions 152 and 185 led to strains that resembled the deletion mutant, but a truncation at position 210 still allowed for full complementation (Figure 4, Table 1). Thus, although the C-terminal tail of ASF1 is less conserved, parts of it are essential for histone binding and sexual development. Leaving only the core intact or shortening the protein to 185 amino acids seems to have the same effect on both processes as a full deletion. A truncation after position 210 removes the part of ASF1 that is only present in ascomycete fungi and does not affect its overall functionality. To rule out the possibility of protein mislocalization as a cause for the inability to complement the deletion mutant, the eGFP-tagged variants were tracked by fluorescence microscopy. All of them co-localized with an mRFP-tagged H3 in the nucleus (Supplemental Figure 8), similar to what was previously shown for wild type ASF1 (6).

**Figure 4.**
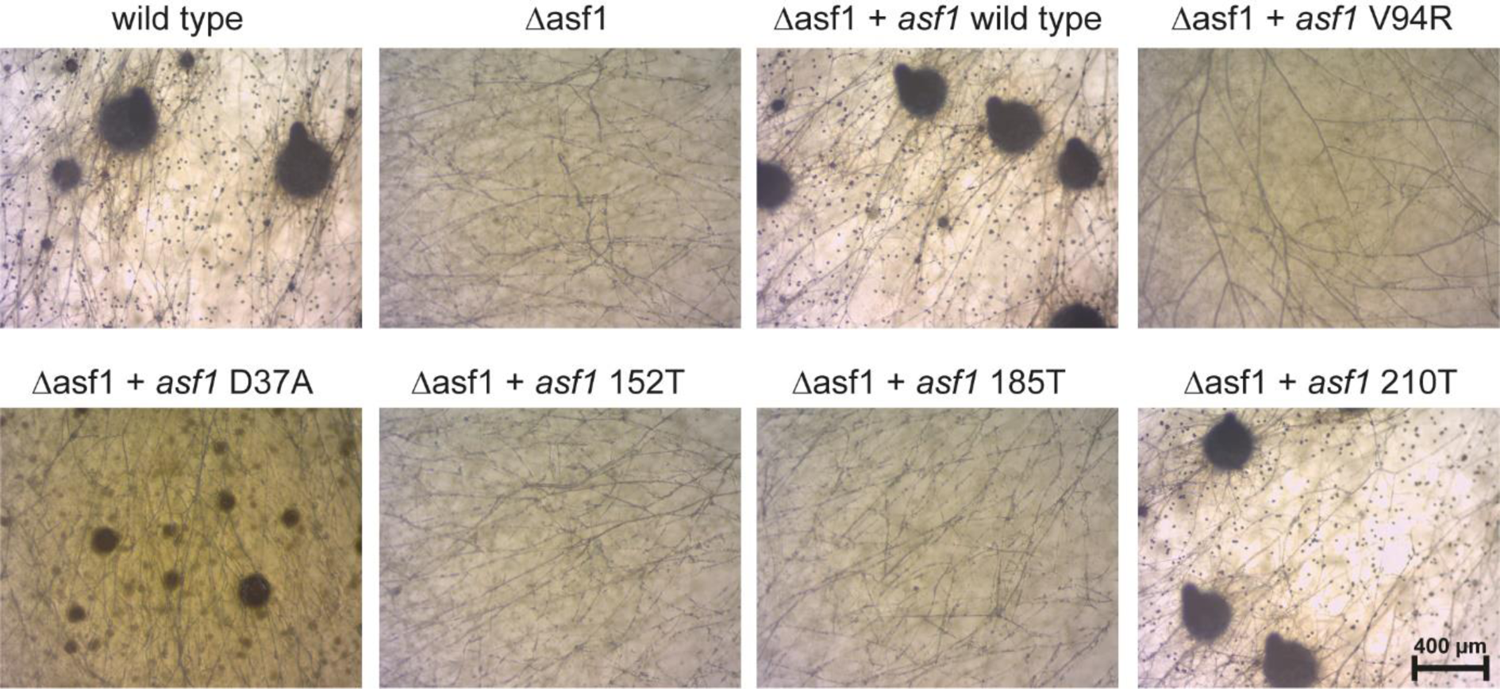
Phenotypes of strains from Δasf1 complementation experiments. The wild type generates mature perithecia in 7 days, while the development of the deletion mutant is blocked at the stage of early protoperithecia and mostly generates only vegetative mycelium. Reintroducing the *asf1* wild type gene into the deletion mutant fully complements the phenotype and lead to strains that are indistinguishable from the wild type. Transforming Δasf1 with the *asf1* V94R variant did not complement the mutant strain resulting in strains with the sterile phenotype of Δasf1. A D37A variant partially complements the mutant, such strains are fertile, but show distinct developmental aberrations like a tendency to generate mostly large protoperithecia and a brown coloration of the mycelium. Transformation with truncated *asf1* variants did not lead to complementation with variants 152T and 185T, but the longer 210T variant led to strains that show a wild type phenotype again.

**Table 1.** Transformants in complementation experiments of Δasf1 with ASF1 variants.

| Encoded ASF1 variant | No. of primary transformants | Fertile transformants | Sterile transformants | Percentage of fertile transformants |
| --- | --- | --- | --- | --- |
| Wild type | 53 | 47 | 6 | 88.7 |
| D37A | 61 | 41 | 20 | 67.2 |
| V94R | 41 | 0 | 41 | 0 |
| 152T | 19 | 0 | 19 | 0 |
| 185T | 22 | 0 | 22 | 0 |
| 210T | 18 | 15 | 3 | 83.3 |

### ASF1 can contribute to DNA damage protection even without histone binding capabilities

Even after confirming the correct localization of tested ASF1 variants, a possible complete loss of function of the protein, i.e. not specifically related to histone binding, could not be completely ruled out. Thus, we looked for a way to confirm functionality of the variants aside from histone binding *in vivo*. The complex network of ASF1 regarding chromatin modification is still poorly understood and direct functions beyond nucleosome assembly are not widely known. It was previously shown in *S. cerevisiae* that ASF1 is involved in DNA damage protection, since Δasf1 mutants are unable to grow on the DNA damage-inducing agent MMS (58). However, in *S. cerevisiae* C-terminal truncations had no impact on DNA damage protection abilities of ASF1 (59, 60). Therefore, we tested *S. macrospora* Δasf1 and the strains expressing the variants for their reaction to the DNA damaging agent MMS, which causes DNA single- and double strand breaks. At a concentration of 0.007 % MMS (v/v), the wild type was able to grow normally on BMM medium, while the mutant did not grow at all (Figure 5). Complementation strains expressing ASF1 variants D37A and 210T showed wild type-like resistance. Although the V94R, 152T and 185T variants of ASF1 are indistinguishable from the deletion strain under normal conditions, an increased resistance against MMS compared to the Δasf1 strain was clearly visible (Figure 5).

**Figure 5.**
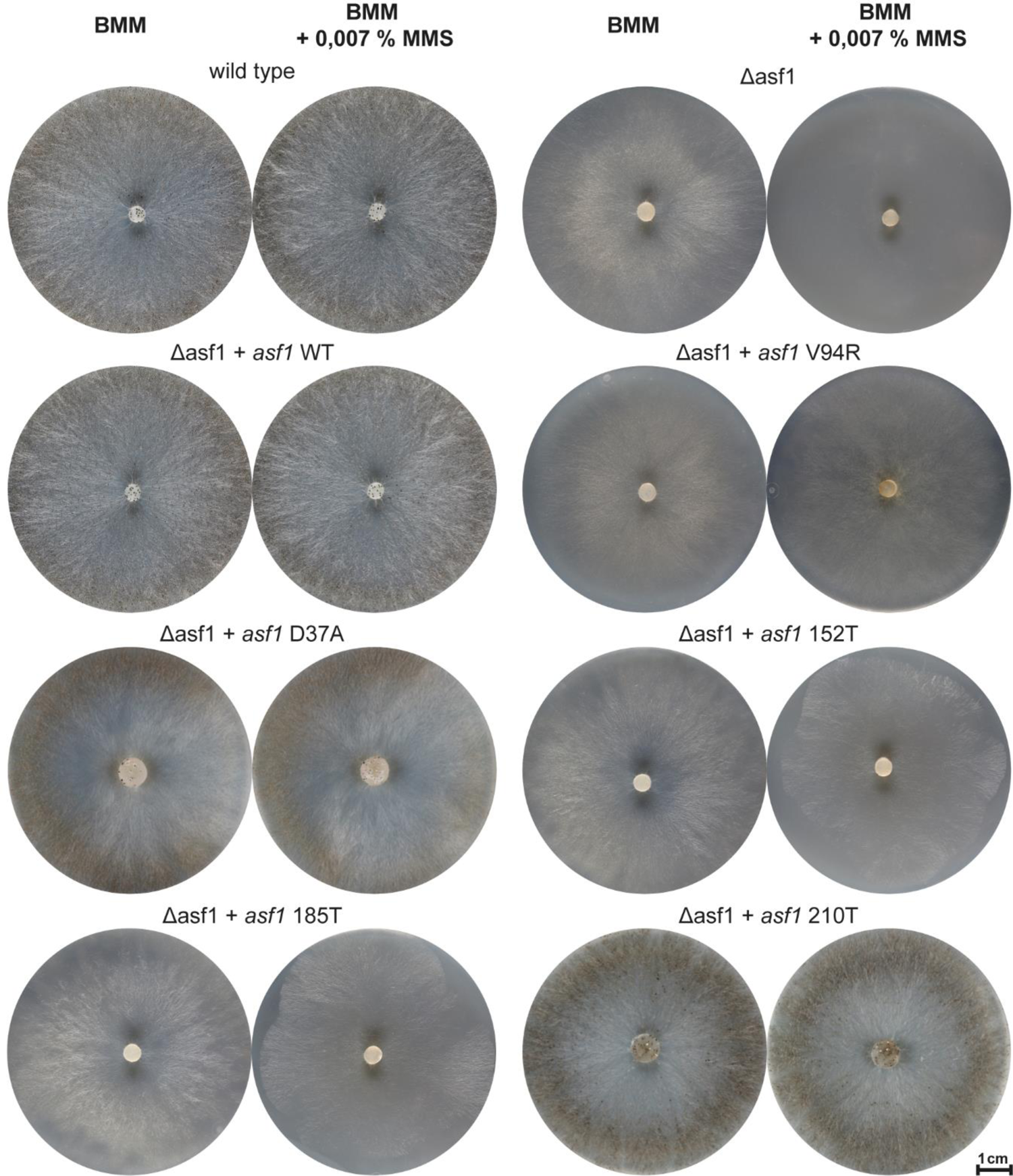
Genotoxic stress resistance of *S. macrospora* strains expressing ASF1 variants. At the given MMS concentration, the wild type was able to grow normally, while the *asf1* deletion mutant exhibited severe sensitivity (no growth). Reintroducing the wild type *asf1* into the mutant restored the resistance. All strains that expressed any kind of ASF1 variant showed at least partial resistance to genotoxic stress, even variants that were shown to be unable to interact with histones and do not complement the developmental defects of *S. macrospora* Δasf1.

Since MMS is known to be a highly reactive chemical and could be inactivated in the media after a short time, we performed a persistence test for our plates. MMS plates were inoculated with the sensitive Δasf1 strain two days after their production and growth was observed for 7 days (Supplementary Figure 9). After the plates reached an age of 7 days, weak growth of the sensitive strain was observed. After this time, Δasf1 appeared to be able to grow on the MMS plates. Therefore, we conclude that although 0.007% MMS plates are volatile and cannot be used for a long time, but during our observation period of 4 days on freshly prepared plates, MMS can be used to assess the resistance of *S. macrospora* to genotoxic stress.

The difference between *S. macrospora* strains expressing these ASF1 variants and the deletion mutant suggests that an uncharacterized function of ASF1 in DNA damage protection exists and these variants are functional in this regard even though some of them are not able to bind histones. Thus, the data suggest that the role of ASF1 in DNA damage protection does not depend on histone interaction.

### Deletion of *asf1* changes global histone modification levels in *S. macrospora*

The functions of ASF1 as a central histone chaperone and its multiple interaction partners during chromatin modification suggest a participation in processes like histone modification. Indeed, ASF1 has been shown to be involved in the acetylation of H3 on the lysines 9 and 56 in *S. cerevisiae* (9) and changes in the deposition of H3K27me3 and H3K9me3 have been observed in *M. musculus* cells during ASF1 depletion conditions (10). To gain insight into the role of ASF1 in histone modification, we performed semi-quantitative western blot screens with protein extracts from *S. macrospora* wild type and Δasf1 for histone modifications H3K27me3, H3K9me, H3K56ac, and H3K9ac. The comparison of the band strength after incubation with antibodies specific for these modifications indicated significant changes in global histone modification levels for H3K27me3 and H3K56ac in the Δasf1 mutant (Figure 6). To gain further insight into the relevance of histone binding for the establishment of histone modifications in ASF1 affected pathways, we also performed semiquantitative screens for ASF1 variants with amino acid substitutions and a truncated C-terminal tail (Figure 6). Our observations mimic our results for complementation capacity and histone binding, as variants that were unable to restore the wild type phenotype and bind histones also showed alterations in H3K27me3 and H3K56ac that were comparable to the deletion mutant. Because of the clear effect of an *asf1* deletion on H3K27me3 and H3K56ac levels, we chose these histone modifications as candidates for ChIP-Seq experiments to verify the results of the screening and to gain information about the distribution of both histone marks. Establishing this method for older mycelium of *S. macrospora* presented challenges regarding the crosslinking and shearing process, as crosslinking agents appeared to create an agglomeration of inaccessible cells. A combination of high input amounts, soft lysis conditions and DNase based shearing allowed us to conduct reproducible ChIP-seq experiments on older *S. macrospora* samples for the first time (Supplemental Text 1, Supplemental Figure 11). All sequenced samples resulted in more than 30 million paired-end reads that showed minimal sequence- or mapping based duplication rates after mapping to the reference genome (Supplemental Figure 12). Duplicates were discarded by MACS2 and broad peak detection was performed for both histone marks (50). The fraction of reads in peaks surpassed the 0.01 threshold (Supplemental Table 5) determined by the ENCODE project for successful ChIP-seq experiments for all samples (61). Repeatability was assessed by Pearson and Spearman correlation and a positive correlation was detected for our replicates (Supplemental Figures 13 and 14). Consistent with the results of Western blot analysis, ChIP-seq revealed that the number of unique ChIP peaks as well as the number of transcriptional start sites and their surrounding 2 kb regions exhibiting H3K27me3 was increased in the deletion mutant whereas H3K56ac appeared to be strongly decreased (Figure 7A, Supplemental Table 6). Although global levels of these histone marks were visibly affected by the deletion of *asf1*, the increase or decrease did not lead to a significant shift of positioning of these marks with regards to the deposition at exons, introns, TSS, TTS or intergenic regions when observed on a relative level (Figure 7B). Therefore, the changes in histone modification strength seem to be more general and not restricted to exons or other functional genomic regions. Representative peaks for exonic regions are shown in Figure 7C, additional representative regions are presented in Supplemental Figure 15.

**Figure 6.**
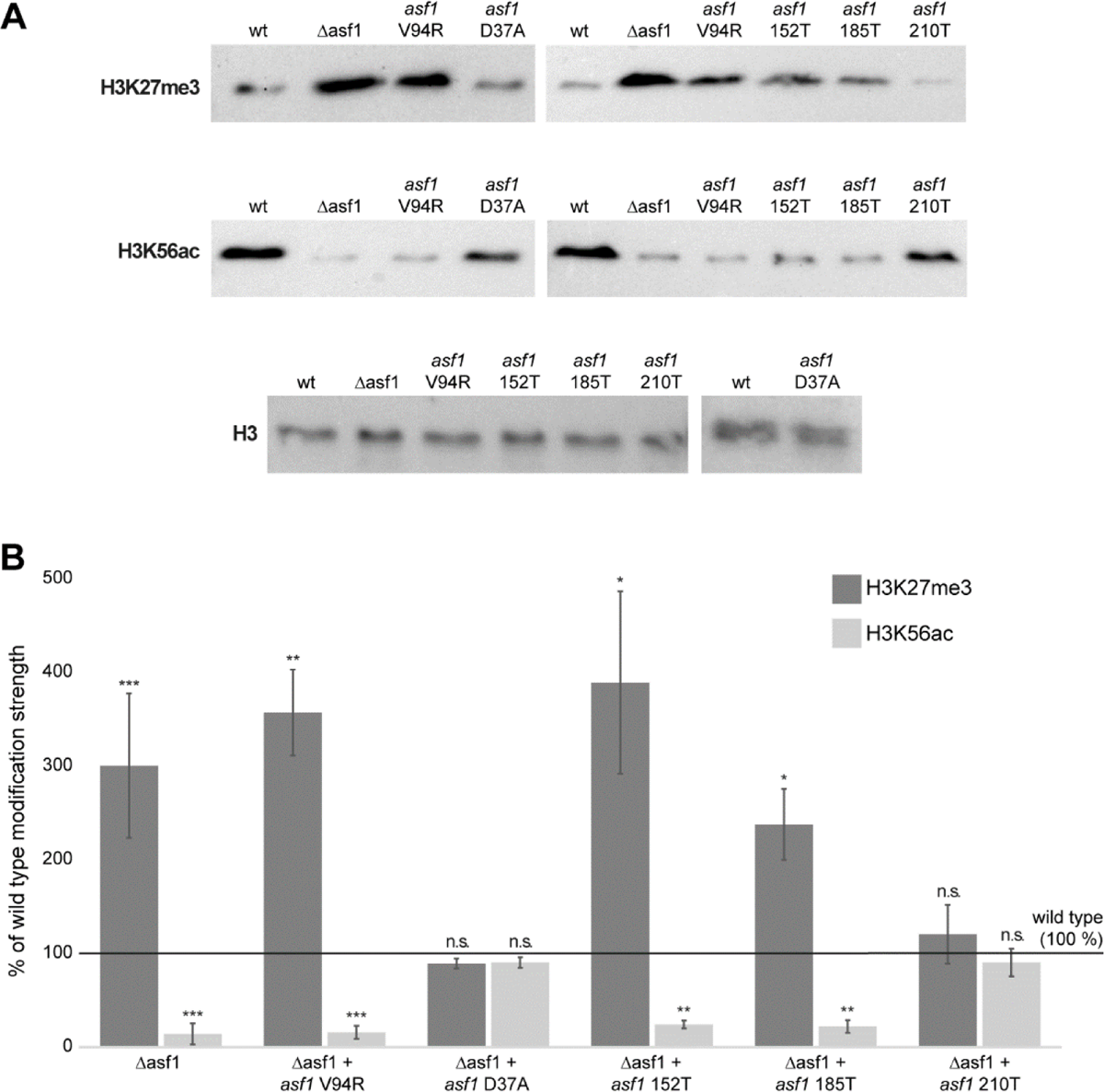
Semiquantitative screening for histone modifications affected by ASF1. **A.** Western blots with antibodies against the indicated histone modifications were used to compare the band strength of equal amounts of protein from wild type, Δasf1 and strains expressing *asf1* variants. H3 levels were measured as an internal control. Uncropped blots are included in Supplemental Figure 10. **B.** Band intensity in Western blots was measured densitometrically and the ratio of mutants to wild type was calculated. The deletion mutant as well as the non-histone binding variants showed a significant increase in H3K27me3 and a significant decrease in H3K56ac. N wild type and Δasf1 = 9; N *asf1* variants = 3. Student’s t-test for significance: * = p-value < 0.05; ** = p-value < 0.01.

**Figure 7.**
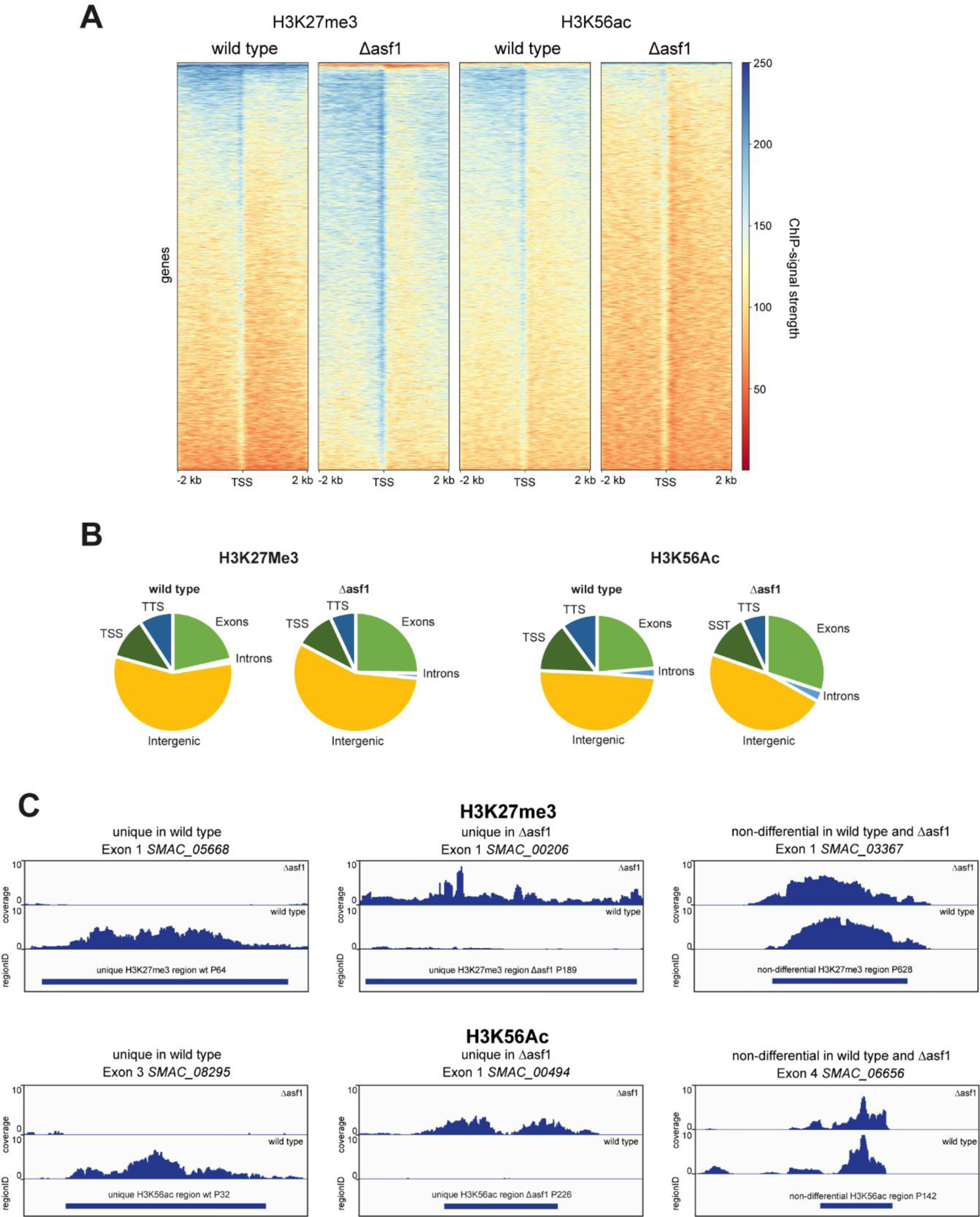
Analysis of histone modifications in the wild type, Δasf1 and *asf1* variants. **A.** Heatmap showing ChIP signal strength at regions surrounding transcription start sites (TSS) in wild type and deletion mutant. *S. macrospora* Δasf1 shows an increase in H3K27me3 and a decrease in H3K56ac compared to the wild type. **B.** Relative distribution of H3K27me3 and H3K56ac in annotated regions of *S. macrospora* in wild type and Δasf1. Although the general amount of both marks underwent significant changes in the deletion mutant, we did not observe a significant change in the relative positioning of H3K27Me3 or H3K56ac, although a slight shift into exonic regions appears likely. **C.** Representative exon regions exhibiting H3K27me3 and H3K56ac. The coverage in the wild type and deletion mutant is shown in the respective area for a unique region in the wild type and the deletion mutant, as well as for a non-differential exon region. Representative regions for introns, promoter regions, downstream regions, and intergenic regions are shown in Supplemental Figures 15 A-C.

### Hi-C reveals a 600 kb duplication in *asf1* deletion mutants that might be linked to their survival

To study the effect of *asf1* and sexual development on overall chromatin structure, we performed Hi-C experiments to map the abundance of DNA-DNA interactions in mycelium at two different developmental stages of the wild type and in mycelium of Δasf1. Samples for the young wild type were grown for 2 d, thus most of the mycelium consisted of vegetative tissue and sexual development was in its early stages. Samples for older mycelium were grown for 5 d, at this stage fully melanized protoperithecia and some early perithecia can be expected. The deletion mutant was cultivated for 3 d (since it is growing somewhat slower than the wild type), but is blocked in its sexual development at the stage of very young protoperithecia. The chromatin of the samples was fixed with formaldehyde and treated with DNase to obtain crosslinked pairs of DNA that are in close proximity at that time. Ends were filled with biotin and re-ligation was performed under high dilution conditions, so ligation events of crosslinked strands are more likely. A streptavidin pulldown was used to generate samples containing interacting DNA fragments of a sample at the given time. By using Dovetail® Dual index primers, Illumina® libraries of around 50 ng were obtained and used for high throughput paired end sequencing. Almost all samples generated over 50 million paired end reads that were mapped to the reference genome and used for the generation of Hi-C maps. Hi-C contacts appeared to be quite low, averaging around 11 %, indicating low complexity (Supplemental Table 7). All recorded interaction events were plotted against their position on the *S. macrospora* genome, each one represented by a red dot. The intensity of red dots at a specific area indicates more recorded DNA-DNA interactions in the sample at the tested point in time. The characteristic diagonal line is an indicator of the high interaction frequency of genomic regions that are always in close proximity to each other, because of their vicinity on the DNA strand itself (Figure 8A). While the resolution of the generated Hi-C maps was too low to make out specific compartments of the young and old wild type, a significant reduction of the general resolution in the sexual mycelium might hint at the development of multiple cell types with different chromatin conformations. During sexual development *S. macrospora* generates multiple different cell types (4), therefore it seems likely that different cell types are characterized by different chromatin conformations. The presence of more than one chromatin conformation in a sample could lead to a loss of signal for mappable DNA-DNA interactions at a specific point.

**Figure 8.**
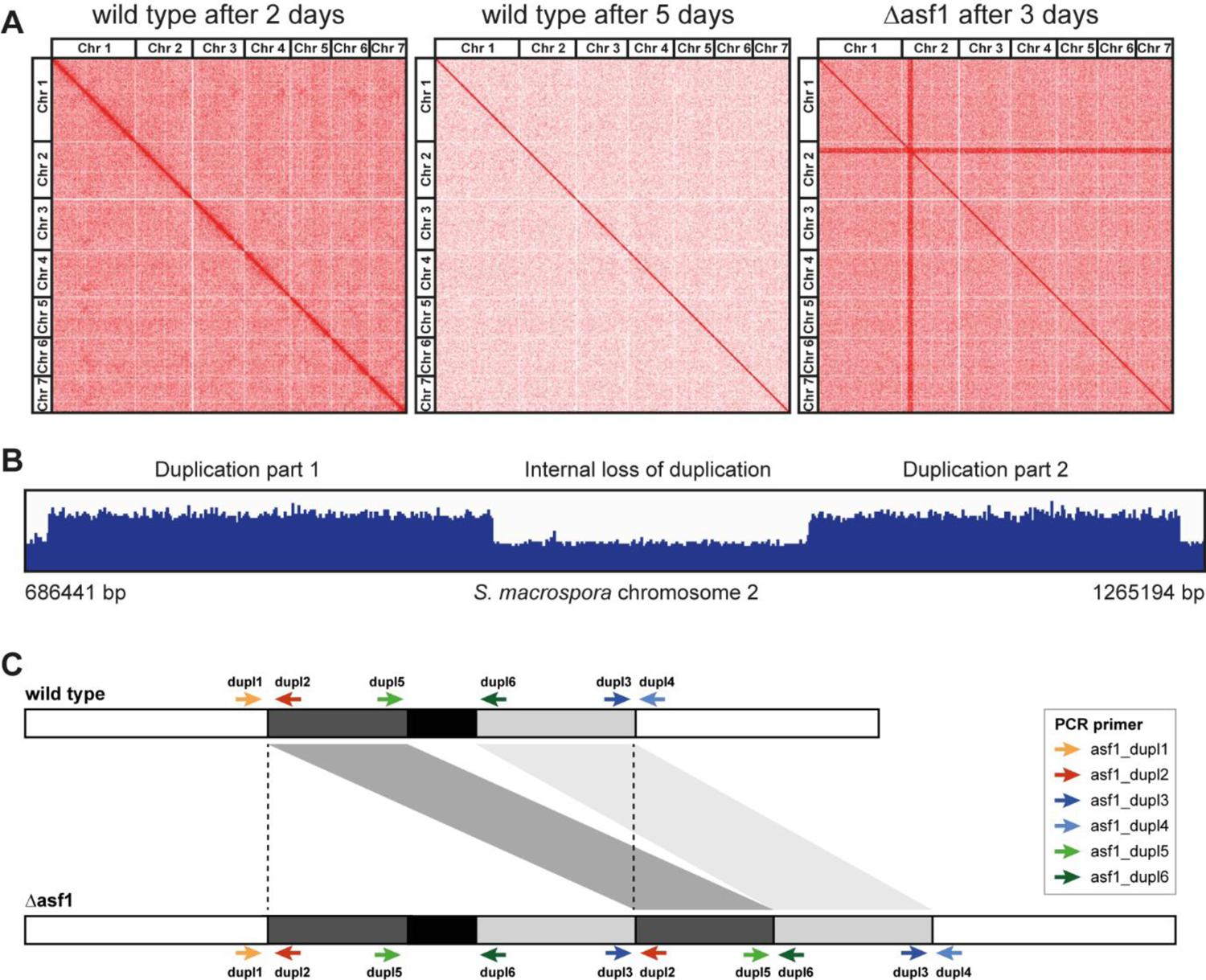
Detection of a 600kb duplication in *S. macrospora* Δasf1 by Hi-C and genome sequencing. **A.** Hi-C analysis of *S. macrospora* wild type and Δasf1. DNA-DNA interactions recorded in the samples at the given time were plotted against both interacting positions in the genome. Each red dot represents an interaction event. The characteristic red diagonal indicates highly increased interaction frequencies of adjacent positions that are always in close proximity to each other. Even in the younger, mostly vegetative state, the resolution was not high enough to make out specific compartments of the genome, even though some areas of heightened interaction frequencies can be observed. The resolution was much lower in the older sample that has progressed further in the developmental cycle. This might be an indicator of different chromatin states that could be present in cells that are only generated during sexual development. Strong horizontal and vertical lines were detectable in the deletion mutant that appeared to indicate higher interaction frequencies originating from a 600 kb are on chromosome 2 and were subsequently shown to be caused by a duplication of a region on chromosome 2. **B.** Whole genome sequencing of *S. macrospora* Δasf1 confirmed the presence of a duplication in a region of around 600 kb on chromosome 2. The alignment of the reads from sequencing of the mutant genome to the wild type reference genome shows that the duplication consists of two parts, indicating either two separate duplication events or the loss of the internal area of the duplicated region. The duplication contains more than 100 genes (Supplemental Table 8). **C.** Schematics for a putative duplication on chromosome 2 of *S. macrospora* Δasf1. Whole genome sequencing proved the presence of a duplication. To test our hypothesis of a tandem duplication with an internal loss event, PCR tests with primers amplifying the genomic borders in the wild type or the newly created, duplication-specific junctions were designed. While the wild type should only yield PCR products for the external borders of the area of interest (asf1_dupl1 x asf1_dupl2, asf1_dupl3 x asf1_dupl4), strains possessing a tandem duplication should also generate products for the new junctions of the duplicated area (asf1_dupl3 x asf1_dupl2) and of the internal region lost in the duplication (asf1_dupl5 x asf1_dupl6).

The Hi-C map of the deletion mutant appeared to show a strong increase of DNA-DNA interactions of a specific region of 600 kb on chromosome 2 with all other genomic regions (enhanced horizontal and vertical bars in Figure 8A). However, such an explanation is biologically unlikely, and a more likely explanation would be that this genomic region in the Δasf1 mutant differs from the wild type genome that was used as a reference for mapping. In this case, this strong increase hinted at a duplication of this region in Δasf1, which would result in twice the number of reads, which then all mapped to the single copy of the wild type genome that was used as a reference for the generation of the Hi-C map. This would also lead to twice the number of (spurious) interactions with other regions of the genome, which would appear as the observed enhanced horizontal and vertical bars in the Hi-C contact map. We detected read patterns in the Hi-C data that match the internal border of a putative tandem duplication of the area of interest. To test this, we sequenced the genome of the Δasf1 mutant by Illumina sequencing. Sequencing reads were mapped to the wild type reference genome, which proved the existence of the duplication in the deletion mutant (Figure 8B, Supplemental Table 8). The detected duplication consists of two compartments, with the middle region of the area of interest not duplicated, hinting at a loss event or two independent duplication events.

PCR tests with primers amplifying the newly created internal junctions of the duplicated area (Figure 8C) revealed the existence of the duplication in all used Δasf1 strains even those that resulted from backcrosses against the wild type. Since the *asf1* gene is located on chromosome 6, whereas the duplication is present on chromosome 2, they should segregate independently resulting in approximately 50 % of Δasf1 strains with and 50 % without the duplication in a cross with a wild type strain. Therefore, we suspected that the viability of the *asf1* deletion might depend on the presence of the duplication. Crossing experiments were performed to analyze the correlation between viable Δasf1 strains and the duplication. The crosses with the spore color mutant fus, which carries the wild type *asf1* allele, resulted 69 strains with an *asf1* wild type allele, of which 21 contained the duplication, matching expectations for free recombination. The 42 Δasf1 mutant strains all possessed the duplication, suggesting that the viability of Δasf1 strains is indeed linked to the occurrence of the duplication on chromosome 2 (Table 2, Supplemental Table 9). While screening for duplication-free Δasf1 strains, we also screened strains carrying *asf1* variants with amino acid substitutions that we used in the complementation analysis, and found the absence of the duplication in several strains containing ASF1 variants V94R, D37A, 185T and 152T (Table 2, Supplemental Table 9). While the absence of the duplication in D37A strains is not surprising since they are fertile and able to interact with histones, the V94R, 185T and 152T variants are unable to bind histones and do not complement the developmental phenotype of *asf1* deletions. This suggests that the presence of an ASF1 protein even without histone binding ability is sufficient to overcome the potential lethality of the complete absence of ASF1, whereas the complementation studies described above show that ASF1 without histone binding ability is not sufficient for fruiting body development. The presence of the duplication does not appear to have any additional effect on the strains carrying it, since all strains show the same phenotype whether the duplication is present or not (fertile for wild type and D37A, sterile for V94R, 185T and 152T, Supplemental Table 9).

**Table 2.** Presence of the duplication on chromosome 2 in wild type, *asf1* deletion strains and strains expressing *asf1* variants. Strains were obtained from crosses of Δasf1 with the spore color mutant fus, which carries the wild type *asf1* gene. The full list of tested strains has been added as Supplemental table 9.

| Genotype | No. of tested strains | No. of strains with duplication | Percentage of strains with duplication |
| --- | --- | --- | --- |
| Wild type | 69 | 21 | 30.4 |
| $\Delta asf1$ | 42 | 42 | 100 |
| $\Delta asf1 + asf1$ D37A | 7 | 2 | 28.6 |
| $\Delta asf1 + asf1$ V94R | 15 | 5 | 30 |
| $\Delta asf1 + asf1$ 152T | 5 | 2 | 40 |
| $\Delta asf1 + asf1$ 185T | 9 | 2 | 22.2 |

## Discussion

### *S. macrospora* ASF1 needs an intact C-terminal region to perform histone binding, an essential function for sexual development

Next to *A. thaliana* (17), *S. macrospora* is one of only two known multicellular organisms that survive a full deletion of *asf1* (even though this appears to be contingent on the presence of the duplication on chromosome 2 as described above), making it a valuable model for ASF1 *in vivo* research, especially when its fast vegetative growth and sexual cycle are taken in account. Therefore, we have the unique opportunity to study the importance of *asf1* for multicellular development. *asf1* deletion strains have been shown to exhibit a slowed vegetative growth rate, and sexual development is blocked at the stage of young protoperithecia (6).

The ability to interact with histones can be regarded as the central requirement of a histone chaperone to perform its functions. ASF1 is known to interact with the histones H3 and H4 and deposits them during nucleosome assembly during transcription, replication and DNA-repair. The general interaction mode of ASF1 with H3 and H4 has been analyzed by *in vitro* experiments with *S. cerevisiae* proteins. The central acidic patch in the core of ASF1 and especially the amino acid V94 are essential in this regard (7). Since the first 155 amino acids of ASF1 are highly conserved in all eukaryotes, similar functions can be assumed. In fact, our co-immunoprecipitation experiments showed a clear dependency of *S. macrospora* ASF1 on V94 to establish interactions with H3 and H4. However, the C-terminal region of ASF1 is quite divergent, with some parts existing in fungi but not in animal or plant models, so information about the functional relevance of this region is sparse. Although experiments in *S. cerevisiae* showed a contribution of the C-terminal region of ASF1 to histone interaction and gene silencing, the specific interaction mode still remains elusive (59). Since the unicellular nature of yeasts does not allow for the analysis of multicellular developmental processes and the specific region of ASF1 is quite divergent, experiments in multicellular eukaryotes like *S. macrospora* might shed more light on the functions of this part of ASF1. Co-immunoprecipitation showed that despite of this region’s divergent nature it is essential for establishing a connection between ASF1 and H3-H4 up until amino acid 210. Thus, a function of the C-terminal tail during this interaction is highly likely. The acidic nature of the C-terminal tail might contribute to facilitating the connection between ASF1 and the histones. A physical interaction between the C-terminal region and H3-H4 has not yet been detected, but might be difficult to prove with structural analysis, because of the usually unfolded C-terminus (58). ASF1 is known to bind histones H3 and H4, but prior to this study, it was not clear if histone binding is required for sexual development. We therefore analyzed different ASF1 variants for their ability to bind histones as well as their ability to support fruiting body formation. Since all ASF1 variants that did not co-immunoprecipitate the histones H3 and H4 were unable to restore the deletion phenotype, the histone binding function of ASF1 appears to be fundamental for sexual and therefore complex multicellular development. Complementation experiments with ASF1 variants that were shown to be able to bind histones led to fully fertile strains, underscoring the importance of this function for normal development. With the exception of ASF1 D37A, these variants generated *S. macrospora* strains that resembled the wild type. D37A strains exhibited a light brown coloration of the mycelium and a tendency to generate aberrant perithecia that resembled giant protoperithecia and exhibited a slightly reduced fertility. Since D37 is the putative interaction site of ASF1 with the histone chaperone complex HIRA (19), one might speculate about an involvement of HIRA during melanin biosynthesis and sexual development.

### Histone binding is not essential for the function of ASF1 during DNA-damage repair

A role of ASF1 during DNA-damage repair without usage of its histone chaperone activity has often been suspected and indeed, in recent studies with human ASF1A, a connection to double strand repair mechanisms independent of histone binding has been discovered (62). In *S. macrospora*, strains expressing ASF1 variants defective in histone binding were able to survive genotoxic stress, whereas Δasf1 proved to be highly sensitive. It can therefore be concluded that the constructed ASF1 variants are at least partially functional in DNA damage repair and that the observed defects in fruiting body development are specifically related to the ability to perform canonical histone chaperone functions. One hypothesis might be that ASF1 is not only a histone chaperone, but also functions as a chromatin modifier hub that regulates and supports the activity of other chromatin-modifying proteins. Regarding the involvement in DNA damage repair, multiple interaction partner candidates are conceivable. A histone-independent interaction with RIF1 has already been demonstrated in human cell lines and contributes to determining the choice of the appropriate double strand break repair mechanism (62). Another candidate involved in DNA damage protection and possibly activated by ASF1 is the histone acetyltransferase RTT109. RTT109 is responsible for H3K9 and K56 acetylation, modifications that are known to be important for DNA damage resistance (63). Interaction between ASF1 and RTT109 has been characterized for *S. cerevisiae* (9) and an activating effect on the histone acetyltransferase was shown, although the exact mechanism remains elusive (64). A defect in acetylation of H3 in *asf1* deletion mutants has been also demonstrated in our ChIP-Seq experiments, therefore a connection between the increased sensitivity of *S. macrospora* Δasf1 to DNA damage and lowered activity of RTT109 seems possible, but the results from H3K56ac screenings for histone-binding defective *asf1*-variants demonstrate that this effect cannot be solely responsible for the increased sensitivity of Δasf1 strains to DNA damage. Strains expressing variants unable to interact with histones, as shown in our interaction study, showed resistance in the DNA damage assay, even though the reduction in H3K56ac was highly similar to the deletion mutant. A combination of multiple factors therefore appears likely. Reduction of H3K56ac might a contributing factor to the MMS sensitivity of the mutant, but additional ASF1-dependent factors might be defective without this central histone chaperone.

Another ASF1-mediated factor for DNA stability is known in *S. cerevisiae* as the checkpoint kinase RAD53 and is involved in DNA-double strand break repair mechanisms. RAD53 needs to be inactivated and bound to ASF1 after repair has been completed, and lack of ASF1 could lead to an overabundance of active RAD53 and subsequent disturbances in the cell cycle and further damage repairs (65). All in all, ASF1 seems to play a crucial role during multiple DNA-damage protection and repair pathways, not all of them relying on ASF1-H3/H4 interactions.

### ASF1 is involved in the histone modification landscape of *S. macrospora*

The histones are targets of various posttranslational modifications. The best known are acetylation, methylation, phosphorylation, sumoylation, and ubiquitination. These modifications vary throughout the genome and shape the histone code, which in turn influences access to the DNA for transcription and the recruiting of transcription factors. The histone landscape of a cell is a significant determinant of the transcriptome (66). ASF1 as a central eukaryotic histone chaperone has been reported to affect multiple histone modifications, such as acetylation of H3K56 (9) and methylation of H3K27 (10). These modifications undergo significant changes after deletion of *asf1* in *S. macrospora*. For H3K27me3, a well-known heterochromatin modification (67), the *asf1* deletion mutant showed 1142 unique ChIP peaks compared to only 401 unique peaks in the wild type. The increase in unique H3K27me3 positions did not seem to be site-specific, as the relative distribution was comparable in the mutant and the wild type. In general, we did not observe a significant change of H3K27me3 in specific functional regions, like transcriptional start sites or gene bodies, instead the global level of this modification was affected as a whole. The increase of H3K27me3 in *asf1* deletion strains could either be a direct consequence of a disruption of ASF1-dependent pathways, or a compensatory reaction to the absence of ASF1. ASF1 is known to be involved in the recycling of histones during DNA replication (7). H3K27me3 is an inheritable histone modification that can persist during replication and the *de novo* integration of histones by ASF1 and CAF1 in parental cells was shown to heavily dilute the inheritance of this mark in nascent chromatin in *A. thaliana* (68). Therefore, a reduction in *de novo* nucleosome assembly through the deletion of a major histone chaperone like ASF1 could reduce general recycling of histones and cause the already established H3K27me3 to persist and accumulate. Especially genomic regions that do not carry H3K27me3 all the time under wild type conditions would show a visible increase under such circumstances, matching our observations of significantly more uniquely marked positions in the deletion mutant. Our observations for non-histone binding ASF1 variants and their deletion mutant-like increase in H3K27me3 suggest a link between the histone interaction ability of ASF1 and the level of H3K27me3 and support the idea that an ASF1, even if present as a protein but unable to interact with histones and therefore probably unable to contribute to nucleosome recycling, may not be able to prevent the persistence of H3K37me3. The blocking of nucleosome recycling and the resulting more permanent marking of histones with H3K27me3 might be an explanation for the severe developmental phenotype of *S. macrospora* Δasf1. Genome wide redistribution of H3K27me3 has been linked to defects in sexual development in *N. crassa*, a close relative of *S. macrospora* (69), so the persistence of this mark in regions where it would exist only temporarily in the wild type might cause similar effects. However, the changes in H3K27me3 distribution could also be a reaction to the reduced DNA methylation observed in Δasf1 (18). In addition to a global reduction of DNA methylation, the deletion of the *S. macrospora asf1* leads to an increase in transcription of many usually weakly expressed genes (18). DNA methylation and H3K27me3 are mostly considered to be marks of transcriptionally silenced regions, but are often regarded as antagonistic to each other (70). Thus, the observed global increase in H3K27me3 in a deletion mutant that also shows a decrease in DNA methylation could be the result of a compensation mechanism for the loss of a transcriptional silencing mechanism. The rising transcription levels resulting from the DNA methylation loss might be kept at least in an acceptable range by using other transcriptional silencers, such as H3K27me3 to keep the deletion mutant alive. This effect might therefore be one of the reasons why *S. macrospora* survives full *asf1* deletion and other multicellular organisms do not.

Histone acetylations on the other hand are known as euchromatin markers, their negative charge can lower the overall positive charge of histones and weaken their binding to the DNA. This can facilitate a more relaxed and open chromatin state that can be accessed by RNA polymerases (71). The higher transcription rates of usually weakly expressed genes after the deletion of *asf1* in *S. macrospora* might suggest that euchromatin marks could be in higher abundance in such a strain. For H3K56ac, our Western blot and ChIP-Seq results showed that at least for this mark, the opposite is the case and the mutant exhibits less H3K56 acetylation on a global scale. As with the heterochromatin mark H3K27me3, we were not able to detect specific patterns of acetylation loss, the overall abundance of the mark appeared to be reduced. Strikingly, we still detected H3K56ac during western blot screenings, as well as ChIP-Seq experiments, while it has been reported that the presence of ASF1 is required to generate this mark in *S. cerevisiae* (72). Interestingly, our screens showed that the establishment of H3K56ac seems to be related to the ability of ASF1 to interact with histones. Histone-binding defective variants showed the same reduction in this mark as Δasf1 strains. Since this mark is generated by the histone acetyltransferase RTT109, which is activated by ASF1 in *S. cerevisiae* (64), it can be speculated that the lower abundance of H3K56ac results from a lack of ASF1-RTT109 interaction. It is speculated that this interaction takes place in the form of a multi-protein complex consisting of ASF1, RTT109 and the histones H3 and H4 together with additional factors (9). This may explain why, even in the presence of ASF1, disruption of histone binding leads to loss of H3K56ac. Additionally, the presence of H3K56ac in *S. macrospora* Δasf1, even though ASF1 was expected to be required for the generation of this mark, hints at either an unknown second activation pathway for RTT109, or an undiscovered way to establish H3K56ac in multicellular ascomycetes. Taken together with the detected increased levels of the heterochromatin mark H3K27me3, lower euchromatic H3K56ac levels would suggest an all-around lower general transcriptional activity of *S. macrospora asf1* deletion mutants. Since the opposite has been documented (18), it can be hypothesized that the contribution of ASF1 to regulating the transcriptome is either not mainly mediated by histone modifications, or is even more complex than previously assumed.

### Hi-C analysis reveals a 600 kb duplication in *S. macrospora* Δasf1 which might be linked to its viability

Methods such as Hi-C allow the analysis of the three-dimensional organization of the chromatin and its effects on the transcriptome and gene regulation. The relationship between chromatin structure and gene regulation seems to be quite complex and possibly both processes can regulate each other (73). In this study, we created DNA-DNA interaction maps of the young, mostly vegetative state of *S. macrospora* and compared it to older mycelium, which has progressed in the cycle of sexual development and an *asf1* deletion mutant with the aim of identifying differences in the chromatin state of those samples. The major challenge was the application of a method that has been fine tuned for single cell assays and fungal protoplasts to older, more mature fungal mycelium. The resulting map of the young wild type showed some areas of the chromatin that suggested higher condensation than other areas, but in general the resolution was not high enough to confidently identify specific interaction compartments. An even lower resolution was visible for the Hi-C map of the older, sexual sample. *S. macrospora* mycelium grows thicker as it ages and sexual structures become reinforced by melanization, which might further complicate the preparation of appropriate Hi-C samples. Crosslinking might be less effective for such samples and the extraction of intact Hi-C pairs might be impaired. The lower DNA-DNA interaction could also be a sign of multiple different chromatin states. Entering sexual development, *S. macrospora* begins generating more specialized cell types, at least 13 different cell types have been identified during fruiting body formation of this fungus (4). Different functions of cells require different transcriptomic processes, which can depend on specific changes of the three-dimensional chromatin state. Higher differentiation of cells in a sample and therefore more divergent chromatin states would reduce the signal-to-noise ratio and would lead to lower detection rates of DNA-DNA interactions. Significant changes in the transcriptome during development are well documented in *S. macrospora* (3). Other studies have demonstrated a strong connection between detectable changes of chromatin architecture and gene expression (74), so it can be assumed that the transcriptional changes of *S. macrospora* during sexual development might be accompanied by significant changes of DNA-DNA interaction frequency. A lower resolution of regularly detectable interactions in the older wild type that is generating multiple cell types supports these assumptions and might be another sign of chromatin state changes that accompany multicellular development. The Hi-C analysis of the Δasf1 mutant revealed a large duplicated region on chromosome 2. We hypothesized that the presence of this duplication might not be just a coincidence, but could be linked to the surprising viability of the Δasf1 mutant in *S. macrospora*. Genetic crosses confirmed co-segregation of the Δasf1 allele on chromosome 6 with the duplication on chromosome 2, whereas the wild type *asf1* allele and the duplication were able to recombine freely. These findings support the hypothesis that the viability of the Δasf1 mutant is coupled to the presence of the duplication. Which specific role this region on chromosome 2 plays in the network of ASF1 is still unclear. So far, no documented interaction partners or parts of the ASF1 network seem to be encoded in this region. The duplication itself seems to be irrelevant for strains expressing *asf1*, even if the ASF1 variant is non-functional with respect to histone binding and sexual development. Strains expressing *asf1* with V94R substitution or truncations after amino acids 185 or 152 were viable but sterile with or without the duplication, and wild type and D37A strains were fertile and did not show additional phenotypes when the duplication was present. One hypothesis might be that the duplication somehow contributes to DNA damage repair or protection, since MMS resistance is the only phenotype that V94R, 185T and 152T confer compared to the *asf1* deletion mutant.

Summarizing our work, we discovered that the ability of ASF1 to interact with histones is essential for sexual development of *S. macrospora*, a function that also depends on the presence of large part of the less conserved C-terminus of ASF1. Histone binding however, proved not to be necessary to allow ASF1 to contribute to the genotoxic stress response. ChIP-seq experiments showed that ASF1 is involved in the establishment of H3K56ac and the repression of H3K27me3. Furthermore, we discovered a 600 kb duplication in the deletion mutant that appears to be linked to the viability of *S. macrospora* Δasf1. Future work might be able to shed light on how ASF1 regulates gene expression by histone modification and which genes of the duplicated area are necessary to stabilize *asf1* deletion mutants.

## Acknowledgements

The authors would like to thank Silke Nimtz, Susanne Schlewinski and Tanja Rollnik for excellent technical assistance and Christopher Grefen for support at the Department of Molecular and Cellular Botany. The authors acknowledge the NGS team of the Omics CF NGS Unit (in development) and CeBiTec. This work was funded by the German Research Foundation (DFG, grant NO407/7-2 to MN). We acknowledge support by the Open Access Publication Funds of the Ruhr-Universität Bochum

