## Supplementary Material for "Histone binding of ASF1 is required for fruiting body development, but not for genome stability in the filamentous fungus *Sordaria macrospora*"

This document contains the following supplemental data:

- Supplemental Tables 1-5, 7, and 9
- Supplemental Figures 1-15
- Supplemental Text 1

Supplemental Tables 6 and 8 are provided separately as Excel files.

**Supplemental Table 1.** *S. macrospora* strains used in this study.

| **Name of strain** | **Genotype** | **Comments** |
|---|---|---|
| SN1693 | Wild type | Reference strain |
| SN1891 | Fus1-1 | Spore color mutant |
| SN1983 | Δasf1::*hph*; sterile | *asf1* deletion mutant |
| SJM 25.4.2 | Δasf1::*hph* + pDS23-ASF1-WT | *asf1* complementation strain with wild type gene |
| SJM 26.8.5 | Δasf1::*hph* + pDS23-ASF1-D37A | *asf1* complementation strain with D37A substitution |
| SJM 27.4.1 | Δasf1::*hph* + pDS23-ASF1-V94R | *asf1* complementation strain with V94R substitution |
| JB 41.1.1 | Δasf1::*hph* + pGG-ASF1-152T | *asf1* complementation strain with truncation in pos. 152 |
| JB 42.1.2 | Δasf1::*hph* + pGG-ASF1-185T | *asf1* complementation strain with truncation in pos. 185 |
| JB 51.1.13 | Δasf1::*hph* + pGG-ASF1-210T | *asf1* complementation strain with truncation in pos. 210 |
| SJM 44.1 | wild type + pGG_H3_Flag + pDS23_asf1_WT | Performance of CoIP experiments with ASF1-eGFP wild type construct and H3-Flag |
| SJM 45.1 | wild type + pGG_H3_Flag + pDS23_asf1_D37A | Performance of CoIP experiments with ASF1-eGFP D37A construct and H3-Flag |
| SJM 35.8 | wild type + pGG_H3_Flag + pDS23_asf1_V94R | Performance of CoIP experiments with ASF1-eGFP V94R construct and H3-Flag |
| SJM 47.1 | wild type + pGG_H3_Flag + pDS23_eGFP | Performance of CoIP experiments with eGFP construct and H3-Flag |
| SJM 48.5 | wild type + pGG_H4_Flag + pDS23_asf1_WT | Performance of CoIP experiments with ASF1-eGFP wild type construct and H4-Flag |
| SJM 49.2 | wild type + pGG_H4_Flag + pDS23_asf1_D37A | Performance of CoIP experiments with ASF1-eGFP D37A construct and H4-Flag |
| SJM 50.2 | wild type + pGG_H4_Flag + pDS23_asf1_V94R | Performance of CoIP experiments with ASF1-eGFP V94R construct and H4-Flag |
| SJM 51.1 | wild type + pGG_H4_Flag + pDS23_eGFP | Performance of CoIP experiments with eGFP construct and H4-Flag |
| JB 25.2 | wild type + pGG_H3_Flag + pGG-ASF1-152T | Performance of CoIP experiments with ASF1-eGFP 152T construct and H3-Flag |
| JB 35.2 | wild type + pGG_H3_Flag + pGG-ASF1-185T | Performance of CoIP experiments with ASF1-eGFP 185T construct and H3-Flag |
| JB 52.2 | wild type + pGG_H3_Flag + pGG-ASF1-210T | Performance of CoIP experiments with ASF1-eGFP 210T construct and H3-Flag |
| JB 34.3 | wild type + pGG_H4_Flag + pGG-ASF1-152T | Performance of CoIP experiments with ASF1-eGFP 152T construct and H4-Flag |
| JB 36.5 | wild type + pGG_H4_Flag + pGG-ASF1-185T | Performance of CoIP experiments with ASF1-eGFP 185T construct and H4-Flag |
| JB 53.2 | wild type + pGG_H4_Flag + pGG-ASF1-210T | Performance of CoIP experiments with ASF1-eGFP 210T construct and H4-Flag |

**Supplemental Table 2.** Oligonucleotides used in this study.

| **Primer** | **Sequence 5'-3'** | **remarks** |
|---|---|---|
| asf1_1 | TCATCGCAGCTTGACTAACAGCTACATGTCTGTC GTTTCGCTTCTCGGGG | *asf1* from start downstream with 5' overlap with plasmid pDS23 |
| asf1_2 | ACAGCTCCTCGCCCTTGCTCACCATTTGAGCCAT GGCCATACCCTGCGGT | *asf1* from end upstream with 5' overlap with plasmid pDS23 |
| asf1_3 | TGGGCGTCACGAGGATTTTGTTGACGTGCGC | for mutagenesis of *asf1* V94R |
| asf1_9 | CGTCAACAAAATCCTCGTGACGCCCAGAAGCTCG | for mutagenesis of *asf1* V94R |
| asf1_5 | GTTCGCGTACCAGCCCTCGAGTGGAAGCTCA | for mutagenesis of *asf1* D37A |
| asf1_6 | TCCACTCGAGGGCTGGTACGCGAACGCGACG | for mutagenesis of *asf1* D37A |
| GG-ASF1-152T-FW | GGCTACGGTCTCGTGGTATGTCTGTCGTTTCGCTTC | for amplification of *asf1* 152T |
| GG-ASF1-152T-RV | GGCTACGGTCTCAATCCCTTGATGGCAAAGCGCGTAAC | for amplification of *asf1* 152T |
| GG-ASF1-185T-FW | GGCTACGGTCTCCTGGTATGTCTGTCGTTTCGCTTCTCG | for amplification of *asf1* 185T |
| GG-ASF1-185T-RV | GGCTACGGTCTCTATCCGGCAAGTTCATCGGCGCC | for amplification of *asf1* 185T |
| GG-ASF1-210T-FW | GGCTACGGTCTCTTGGTATGTCTGTCGTTTCGCTTCTC | for amplification of *asf1* 210T |
| GG-ASF1-210T-RV | GGCTACGGTCTCCATCCGACGATGGCGCCGTTCAATTG | for amplification of *asf1* 210T |
| asf1_veri_fw | TCAATCTTGTCGCTAACCG | verification of *asf1* presence in PCR tests |
| asf1_veri_rv | GTCAACACTGCCATCCTC | verification of *asf1* presence in PCR tests |
| egfp_rv | ACTTGTGGCCGTTTACGTCG | verification of *asf1* presence in PCR tests by binding to the eGFP tag of complementation vectors |
| 1751 | GCCATATTTTCCTGCTCTCC | verification of *asf1* presence in PCR tests by binding to the gpd promotor of complementation vectors |
| asf1_dupl1 | CTCACCAGACCTGAAGTTGCGTCC | verification of duplication in Δasf1 strains |
| asf1_dupl2 | TAGAGGTACACTACGCCCATCTCC | verification of duplication in Δasf1 strains |
| asf1_dupl3 | TCAGCGATCATTTGTCAGTGTCCG | verification of duplication in Δasf1 strains |
| asf1_dupl4 | GTGGGTGGAAAGTCAAAAGTCTGG | verification of duplication in Δasf1 strains |
| asf1_dupl5 | ATATAGCTGGTGATCCCTTCCAGC | verification of deletion in duplication in Δasf1 strains |
| asf1_dupl6 | ATGGGGAGATTACACAGGTGTTCG | verification of deletion in duplication in Δasf1 strains |

**Supplemental Table 3.** Plasmids used in this study.

| **Plasmid** | **Characteristics** | **Comments** |
|---|---|---|
| pDS23 | 8,8 kb, P*gpd::eGFP::*TrpC*, ura, amp, lacZ* | eGFP control for CoIP experiments, cloning basis for complementation vectors, eGFP tag |
| pDS23_asf1_WT | 9,9 kb, pDS23-derivate, asf1(WT)-*egfp* | *asf1* wild type complementation and CoIP vector, eGFP tag |
| pDS23_asf1_D37A | 9,9 kb, pDS23-derivate, *asf1*(D37A)-*egfp* | *asf1* D37A complementation and CoIP vector, eGFP tag |
| pDS23_asf1_V94R | 9,9 kb, pDS23-derivate, *asf1*(V94R)-*egfp* | *asf1* V94R complementation and CoIP vector, eGFP tag |
| pGG-H3-Flag | 5,8 kb, P*trpC*::*Nat*, P*gpd*, 3xFlag, *LacZa*, T*rpC*, *amp*, pUC-ori, *SMAC_02363* (histone H3) | H3 CoIP vector, FLAG tag |
| pGG-H4-Flag | 5,8 kb, P*trpC*::*Nat*, P*gpd*, 3xFlag, *LacZa*, T*rpC*, *amp*, pUC-ori,*SMAC_02364* (histone H4) | H4 CoIP vector, FLAG tag |
| pGG-ASF1-152T | 6,6 kb, P*trpC*::*Nat*, P*gpd*, eGFP, *LacZa*, T*rpC*, *amp*, pUC-ori, *asf1*(152T)-*egfp* | *asf1* 152T complementation and CoIP vector, eGFP tag |
| pGG-ASF1-185T | 6,8 kb, P*trpC*::*Nat*, P*gpd*, eGFP, *LacZa*, T*rpC*, *amp*, pUC-ori, *asf1*(185T)-*egfp* | *asf1* 185T complementation and CoIP vector, eGFP tag |
| pGG-ASF1-210T | 6,9 kb, P*trpC*::*Nat*, P*gpd*, eGFP, *LacZa*, T*rpC*, *amp*, pUC-ori, *asf1*(210T)-*egfp* | *asf1* 210T complementation and CoIP vector, eGFP tag |

**Supplemental table 4.** Western Blot protocols used in this study.

| **H3K27me3 detection** | **H3K9me detection** | **H3K56ac detection** |
|---|---|---|
| 4 h blocking in TBST (0,1 % Tween-20) with 5 % (w/v) non-fat dry milk<br><br>10 min wash in TBST (0,1 % Tween-20)<br><br>16 h incubation Anti H3K27Me3 (Cell Signaling) 1:1000 in TBST (0,1 % Tween-20) with 5 % (w/v) non-fat dry milk<br><br>3 x 10 min wash in TBST (0,1 % Tween-20)<br><br>1 h incubation Anti-Rabbit IgG HRP linked (Cell Signaling) 1:5000 in TBST (0,1 % Tween-20) with 5 % (w/v) non-fat dry milk<br><br>3 x 10 min wash in TBST (0,1 % Tween-20) | 4 h blocking in TBST (0,1 % Tween-20) with 5 % (w/v) non-fat dry milk<br><br>10 min wash in TBST (0,1 % Tween-20)<br><br>16 h incubation Anti H3K9Me (Merck Millipore) 1:1000 in TBST (0,1 % Tween-20) with 5 % (w/v) non-fat dry milk<br><br>3 x 10 min wash in TBST (0,1 % Tween-20)<br><br>1 h incubation Anti-Rabbit IgG HRP linked (Cell Signaling) 1:5000 in TBST (0,1 % Tween-20) with 5 % (w/v) non-fat dry milk<br><br>3 x 10 min wash in TBST (0,1 % Tween-20) | 4 h blocking in TBST (0,1 % Tween-20) with 1 % (w/v) BSA<br><br>16 h incubation Anti H3K56Ac (Active Motif) 1:1000 in TBST (0,1 % Tween-20) with 1 % (w/v) BSA<br><br>10 min wash in TBST (0,1 % Tween-20)<br><br>2 h incubation Anti-Rabbit IgG HRP linked (Cell Signaling) 1:5000 in TBST (0,1 % Tween-20) with 1 % (w/v) BSA<br><br>3 x 10 min wash in TBST (0,1 % Tween-20) |
| **H3K9Ac** | **eGFP tag detection** | **FLAG tag detection** |
| 4 h blocking in TBST (0,1 % Tween-20) with 1 % (w/v) BSA<br><br>16 h incubation Anti H3K9Ac (Active Motif) 1:1000 in TBST (0,1 % Tween-20) with 1 % (w/v) BSA<br><br>10 min wash in TBST (0,1 % Tween-20)<br><br>2 h incubation Anti-Rabbit IgG HRP linked (Cell Signaling) 1:5000 in TBST (0,1 % Tween-20) with 1 % (w/v) BSA<br><br>10 min wash in TBST (0,1 % Tween-20) | 1 h blocking in PBST (0,1 % Tween-20) with 5 % (w/v) non-fat dry milk<br><br>10 min wash in PBS<br><br>1h incubation in living colors JL-8 (Clontech) 1:2000 in PBST (0,1 % Tween-20) with 5 % (w/v) non-fat dry milk<br><br>2 x 5 min wash in PBST (0,1 % Tween-20)<br><br>1 h incubation Anti-Mouse IgG HRP linked (Cell Signaling) 1:1000 in PBST (0,1 % Tween-20) with 5 % (w/v) non-fat dry milk<br><br>4 x 10 min wash in PBST (0,1 % Tween-20) | 1 h blocking in TBST (0,05 % Tween-20) with 5 % (w/v) non-fat dry milk<br><br>10 min wash in TBS<br><br>1h incubation Mouse Anti-FLAG M2 (Sigma-Aldrich) 1:2000 in TBST (0,05 % Tween-20) with 5 % (w/v) non-fat dry milk<br><br>10 min wash TBS<br><br>1 h incubation Anti-Mouse IgG HRP linked (Cell Signaling) 1:5000 in TBST (0,05 % Tween-20) with 5 % (w/v) non-fat dry milk<br><br>3 x 10 min wash in TBST (0,05 % Tween-20) |

**Supplemental table 5.** ChIP-seq fraction of reads in peaks.

| **H3K27me3** | wild type rep 1 | wild type rep 2 | Δasf1 rep 1 | Δasf1 rep 2 |
|---|---|---|---|---|
| Obtained reads | 51921558 | 47584564 | 40282052 | 47681632 |
| Reads in peaks | 9358549 | 9875478 | 12034107 | 15859431 |
| Fraction of reads in peaks | 18.02 % | 20.75 % | 29.87 % | 33.26 % |
| **H3K56ac** | wild type rep 1 | wild type rep 2 | Δasf1 rep 1 | Δasf1 rep 2 |
| Obtained reads | 30947548 | 43537509 | 34128031 | 44521758 |
| Reads in peaks | 9689591 | 11503774 | 2535030 | 2037400 |
| Fraction of reads in peaks | 31.31 % | 26.42 % | 7.43 % | 4.58 % |

**Supplemental table 7.** Hi-C ligation events.

| Strain | Genotype | Replicate | Obtained reads | Mapped pairs | Hi-C contacts | % of reads in Hi-C contacts |
|---|---|---|---|---|---|---|
| SN1693 | wild type young | 1 | 50280915 | 46429907 | 4191907 | 8.33 |
| SN1693 | wild type young | 2 | 55234668 | 51511005 | 10427734 | 18.87 |
| SN1693 | wild type old | 1 | 48173321 | 43876915 | 2534375 | 5.26 |
| SN1693 | wild type old | 2 | 149315718 | 135385996 | 18809722 | 12.59 |
| SN1983 | Δasf1 | 1 | 50056244 | 45904670 | 5244496 | 10.47 |
| SN1983 | Δasf1 | 2 | 55780155 | 50555623 | 8609712 | 15.43 |

**Supplemental table 9.** Strains tested for the duplication on chromosome 2.

| Name of strain | Genotype at the *asf1* gene locus | Duplication on chromosome 2 | Fertility |
|---|---|---|---|
| S 689 | wild type | not present | fertile |
| S 690 | wild type | present | fertile |
| S 692 | wild type | present | fertile |
| S 693 | wild type | present | fertile |
| S 694 | wild type | not present | fertile |
| S 695 | wild type | not present | fertile |
| S 696 | wild type | not present | fertile |
| S 697 | wild type | not present | fertile |
| S 698 | wild type | not present | fertile |
| S 699 | wild type | present | fertile |
| S 700 | wild type | present | fertile |
| S 701 | wild type | not present | fertile |
| S 703 | wild type | present | fertile |
| S 704 | wild type | not present | fertile |
| S 705 | wild type | not present | fertile |
| S 706 | wild type | not present | fertile |
| S 707 | wild type | not present | fertile |
| S 708 | wild type | not present | fertile |
| S 709 | wild type | not present | fertile |
| S 710 | wild type | present | fertile |
| S 896 | wild type | not present | fertile |
| S 897 | wild type | present | fertile |

| | | | |
|---|---|---|---|
| S 898 | wild type | not present | fertile |
| S 899 | wild type | not present | fertile |
| S 900 | wild type | not present | fertile |
| S 901 | wild type | not present | fertile |
| S 902 | wild type | not present | fertile |
| S 903 | wild type | not present | fertile |
| S 904 | wild type | not present | fertile |
| S 905 | wild type | not present | fertile |
| S 906 | wild type | present | fertile |
| S 907 | wild type | not present | fertile |
| S 908 | wild type | not present | fertile |
| S 909 | wild type | not present | fertile |
| S 910 | wild type | present | fertile |
| S 911 | wild type | present | fertile |
| S 912 | wild type | present | fertile |
| S 913 | wild type | not present | fertile |
| S 918 | wild type | not present | fertile |
| S 919 | wild type | present | fertile |
| S 920 | wild type | not present | fertile |
| S 922 | wild type | not present | fertile |
| S 923 | wild type | present | fertile |
| S 924 | wild type | not present | fertile |
| S 925 | wild type | present | fertile |
| S 926 | wild type | not present | fertile |
| S 937 | wild type | not present | fertile |
| S 938 | wild type | not present | fertile |
| S 939 | wild type | present | fertile |
| S 940 | wild type | not present | fertile |
| S 941 | wild type | present | fertile |
| S 942 | wild type | not present | fertile |
| S 943 | wild type | not present | fertile |
| S 944 | wild type | not present | fertile |
| S 945 | wild type | not present | fertile |
| S 946 | wild type | not present | fertile |
| S 947 | wild type | not present | fertile |
| S 948 | wild type | present | fertile |
| S 949 | wild type | present | fertile |
| S 950 | wild type | not present | fertile |
| S 951 | wild type | not present | fertile |
| S 952 | wild type | present | fertile |
| S 953 | wild type | not present | fertile |
| S 954 | wild type | not present | fertile |
| S 955 | wild type | not present | fertile |
| S 956 | wild type | not present | fertile |
| S 957 | wild type | not present | fertile |
| S 958 | wild type | not present | fertile |
| S 963 | wild type | present | fertile |
| S 28 | Δasf1 | present | sterile |
| S 40 | Δasf1 | present | sterile |
| S 49 | Δasf1 | present | sterile |

| | | | |
|---|---|---|---|
| S 50 | Δasf1 | present | sterile |
| S 54 | Δasf1 | present | sterile |
| S 57 | Δasf1 | present | sterile |
| S 691 | Δasf1 | present | sterile |
| S 702 | Δasf1 | present | sterile |
| S 790 | Δasf1 | present | sterile |
| S 807 | Δasf1 | present | sterile |
| S 914 | Δasf1 | present | sterile |
| S 915 | Δasf1 | present | sterile |
| S 916 | Δasf1 | present | sterile |
| S 917 | Δasf1 | present | sterile |
| S 921 | Δasf1 | present | sterile |
| S 927 | Δasf1 | present | sterile |
| S 928 | Δasf1 | present | sterile |
| S 929 | Δasf1 | present | sterile |
| S 930 | Δasf1 | present | sterile |
| S 931 | Δasf1 | present | sterile |
| S 932 | Δasf1 | present | sterile |
| S 933 | Δasf1 | present | sterile |
| S 934 | Δasf1 | present | sterile |
| S 935 | Δasf1 | present | sterile |
| S 936 | Δasf1 | present | sterile |
| S 959 | Δasf1 | present | sterile |
| S 960 | Δasf1 | present | sterile |
| S 961 | Δasf1 | present | sterile |
| S 962 | Δasf1 | present | sterile |
| S 964 | Δasf1 | present | sterile |
| S 980 | Δasf1 | present | sterile |
| S 981 | Δasf1 | present | sterile |
| S 993 | Δasf1 | present | sterile |
| S 1015 | Δasf1 | present | sterile |
| S 1019 | Δasf1 | present | sterile |
| S 1023 | Δasf1 | present | sterile |
| S 1025 | Δasf1 | present | sterile |
| J 3 | Δasf1 | present | sterile |
| J 14 | Δasf1 | present | sterile |
| J 37 | Δasf1 | present | sterile |
| J 71 | Δasf1 | present | sterile |
| J 78 | Δasf1 | present | sterile |
| SJM 26.8.1 | Δasf1 + pDS23-ASF1-D37A | not present | fertile |
| SJM 26.8.4 | Δasf1 + pDS23-ASF1-D37A | not present | fertile |
| SJM 26.8.5 | Δasf1 + pDS23-ASF1-D37A | not present | fertile |
| SJM 26.8.8 | Δasf1 + pDS23-ASF1-D37A | not present | fertile |
| SJM 26.8.9 | Δasf1 + pDS23-ASF1-D37A | present | fertile |
| SJM 26.8.11 | Δasf1 + pDS23-ASF1-D37A | not present | fertile |
| SJM 26.8.12 | Δasf1 + pDS23-ASF1-D37A | present | fertile |
| SJM 27.4.1 | Δasf1 + pDS23-ASF1-V94R | not present | sterile |
| JB 13.1.1 | Δasf1 + pDS23-ASF1-V94R | not present | sterile |
| JB 13.1.2 | Δasf1 + pDS23-ASF1-V94R | not present | sterile |
| JB 13.1.3 | Δasf1 + pDS23-ASF1-V94R | present | sterile |

| | | | |
|---|---|---|---|
| JB 13.1.4 | Δasf1 + pDS23-ASF1-V94R | present | sterile |
| JB 13.1.5 | Δasf1 + pDS23-ASF1-V94R | present | sterile |
| JB 13.1.6 | Δasf1 + pDS23-ASF1-V94R | not present | sterile |
| JB 13.1.7 | Δasf1 + pDS23-ASF1-V94R | not present | sterile |
| JB 13.1.8 | Δasf1 + pDS23-ASF1-V94R | not present | sterile |
| JB 13.1.9 | Δasf1 + pDS23-ASF1-V94R | not present | sterile |
| JB 13.1.10 | Δasf1 + pDS23-ASF1-V94R | not present | sterile |
| JB 13.1.11 | Δasf1 + pDS23-ASF1-V94R | present | sterile |
| JB 13.1.12 | Δasf1 + pDS23-ASF1-V94R | present | sterile |
| JB 13.1.13 | Δasf1 + pDS23-ASF1-V94R | not present | sterile |
| JB 13.1.114 | Δasf1 + pDS23-ASF1-V94R | not present | sterile |
| JB 41.1.1 | Δasf1 + pGG-ASF1-152T | not present | sterile |
| JB 41.1.3 | Δasf1 + pGG-ASF1-152T | not present | sterile |
| JB 41.1.18 | Δasf1 + pGG-ASF1-152T | present | sterile |
| JB 41.1.22 | Δasf1 + pGG-ASF1-152T | not present | sterile |
| JB 41.4.3 | Δasf1 + pGG-ASF1-152T | present | sterile |
| JB 42.1.2 | Δasf1 + pGG-ASF1-185T | present | sterile |
| JB 42.1.3 | Δasf1 + pGG-ASF1-185T | present | sterile |
| JB 42.2.1 | Δasf1 + pGG-ASF1-185T | not present | sterile |
| JB 42.2.2 | Δasf1 + pGG-ASF1-185T | not present | sterile |
| JB 42.3.1 | Δasf1 + pGG-ASF1-185T | not present | sterile |
| JB 42.4.1 | Δasf1 + pGG-ASF1-185T | not present | sterile |
| JB 42.4.3 | Δasf1 + pGG-ASF1-185T | not present | sterile |
| JB 42.5.1 | Δasf1 + pGG-ASF1-185T | not present | sterile |
| JB 42.6.1 | Δasf1 + pGG-ASF1-185T | not present | sterile |

**A**

| genome version | **v04** | **v03** |
|---|---|---|
| genome size [Mb] | 39.4 | 38.9 |
| N50 [kb] | 5713 | 180 |
| no. of contigs/scaffolds | 7 | 584 |
| no. of gaps in contigs/scaffolds | 0 | 216 |
| largest contig [kb] | 9310 | 664 |
| smallest contig [kb] | 4093 | 1 |
| GC content [%] | 52.1 | 52.0 |
| no. of protein-coding genes | 10420 | 9874 |

**B**

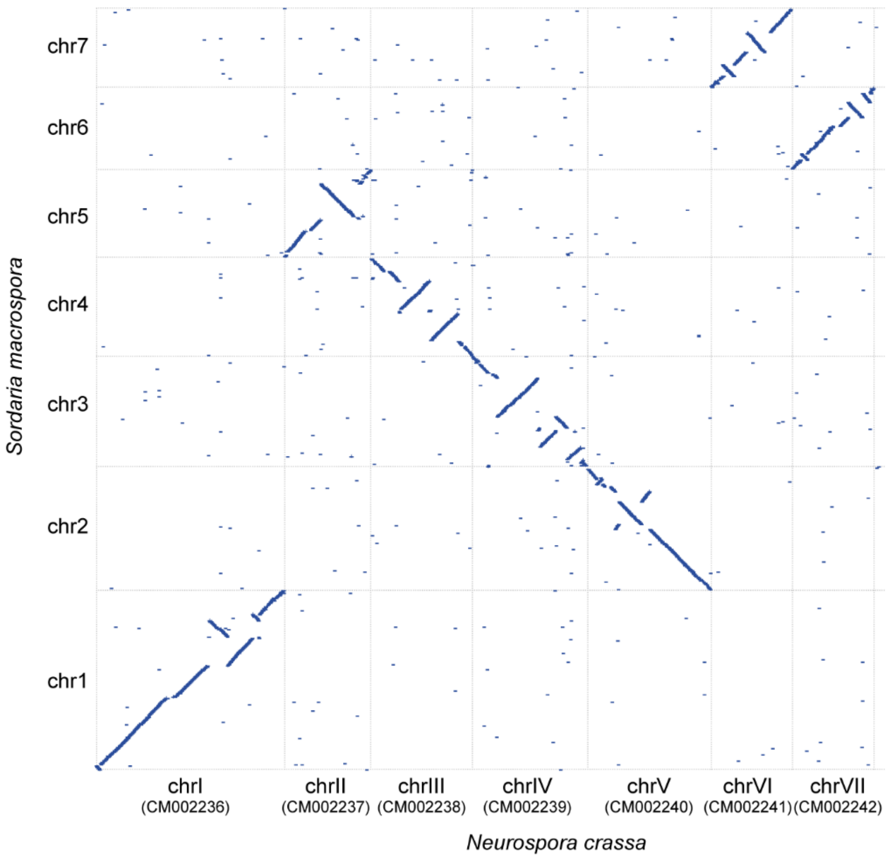


**Supplemental Figure 1.** Features of the *Sordaria macrospora* genome assembly v04. **A.** Core features of the *S. macrospora* assembly versions v04 (this study) and v03 (Blank-Landeshammer et al. 2019, mBio 10:e02367-02319). **B.** Nucmer (Kurtze et al. 2004, Genome Biol 5:R12) comparison of the genome assemblies of *S. macrospora* and its close relative *Neurospora crassa* (FungiDB version 52; Galagan et al. 2003, Nature 422:859-868; Basenko et al. 2018, J Fungi 4:39). The nuclear genomes of both ascomycetes comprise seven chromosomes that are largely syntenic except for several inversions or shuffling of regions within each chromosome.

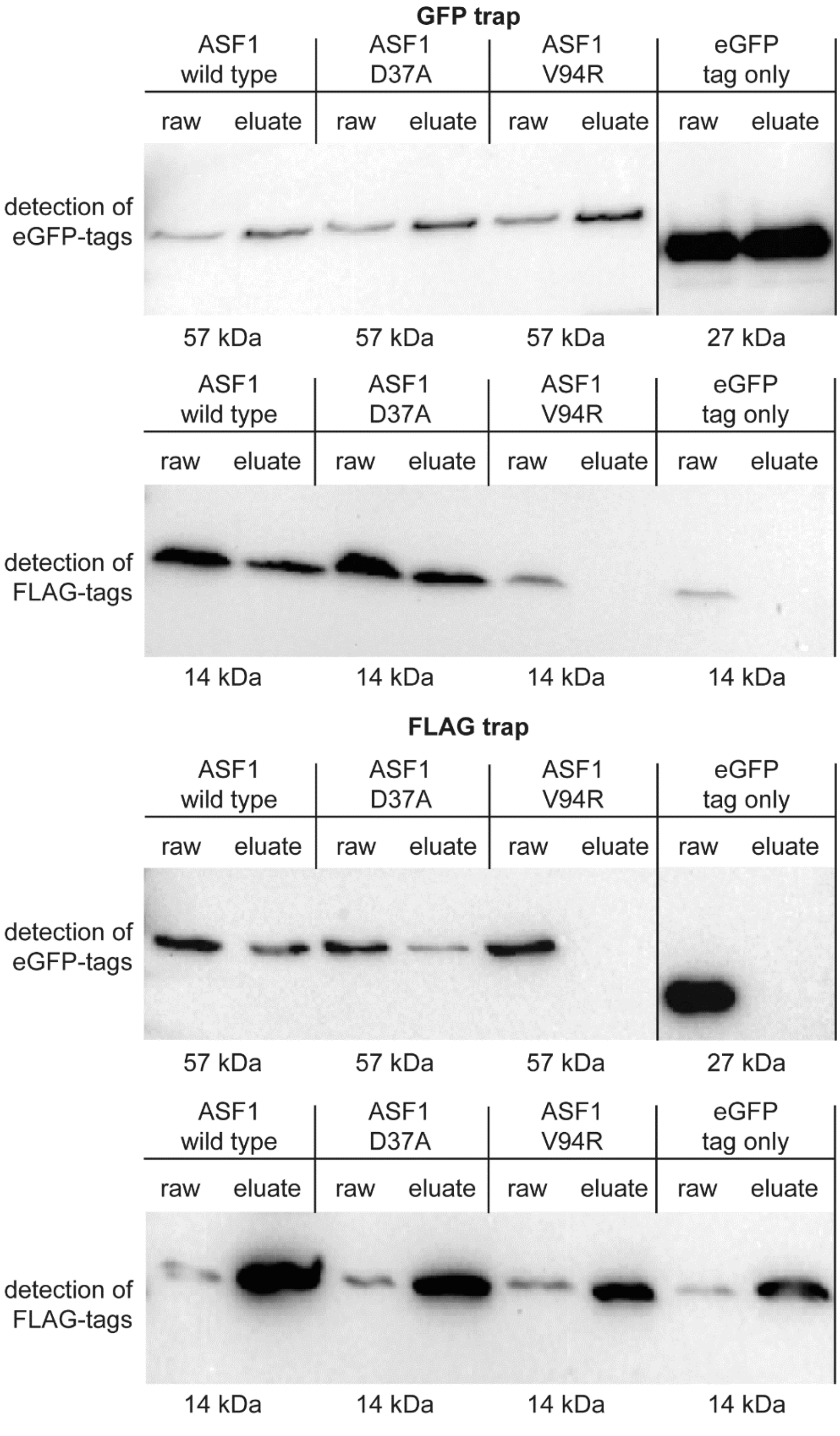


**Supplemental Figure 2.** Co-immunoprecipitation results for ASF1 variants with amino acid substitutions and histone H4. GFP-tagged ASF1 wild type and variants D37A and V94R were used as potential interaction partners for Flag-tagged histone H4 in a GFP trap and a Flag trap. Results were checked by western blot analysis with antibodies against GFP and Flag tags. ASF1 wild type and the D37A variant showed signals for bait and prey proteins in the raw and eluate sample, indicating interaction, whereas the V94R variant showed the signal for the prey protein only in the raw sample. Strains expressing non-fused GFP with the corresponding Flag-tagged H4 were used as a negative control. Uncropped blots are shown in Supplementary Figure 3B.

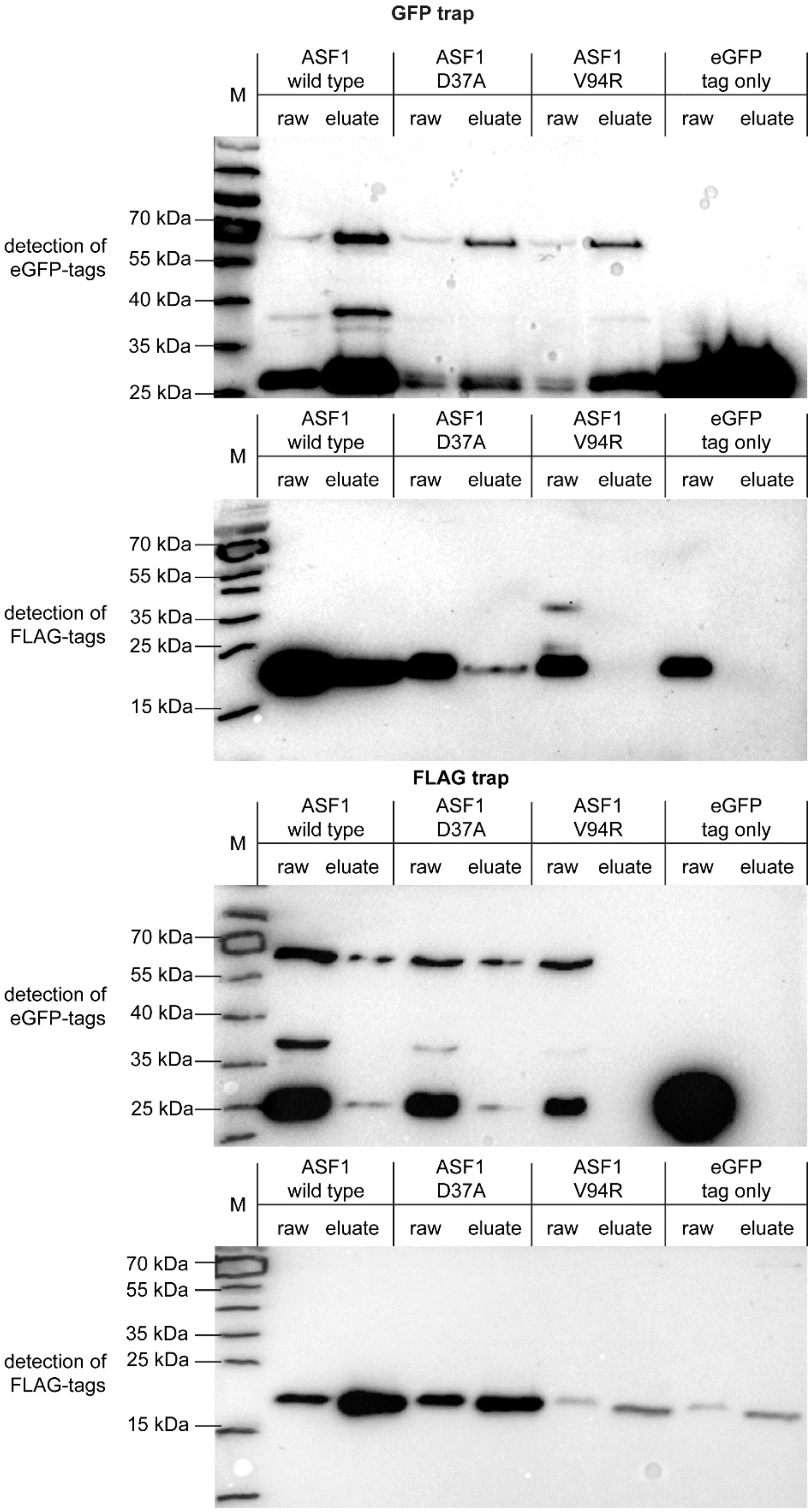


**Supplemental Figure 3A.** Uncropped blots - Co-immunoprecipitation results for ASF1 variants with amino acid substitutions and histone H3. GFP-tagged ASF1 wild type and variants D37A and V94R were used as potential interaction partners for Flag-tagged histone H3 in a GFP trap and a Flag trap. Results were checked by western blot analysis with antibodies against GFP and Flag tags. ASF1 wild type and the D37A variant showed signals for bait and prey proteins in the raw and eluate sample, indicating interaction, whereas the V94R variant showed the signal for the prey protein only in the raw sample. Strains expressing non-fused GFP with the corresponding Flag-tagged H3 were used as a negative control.

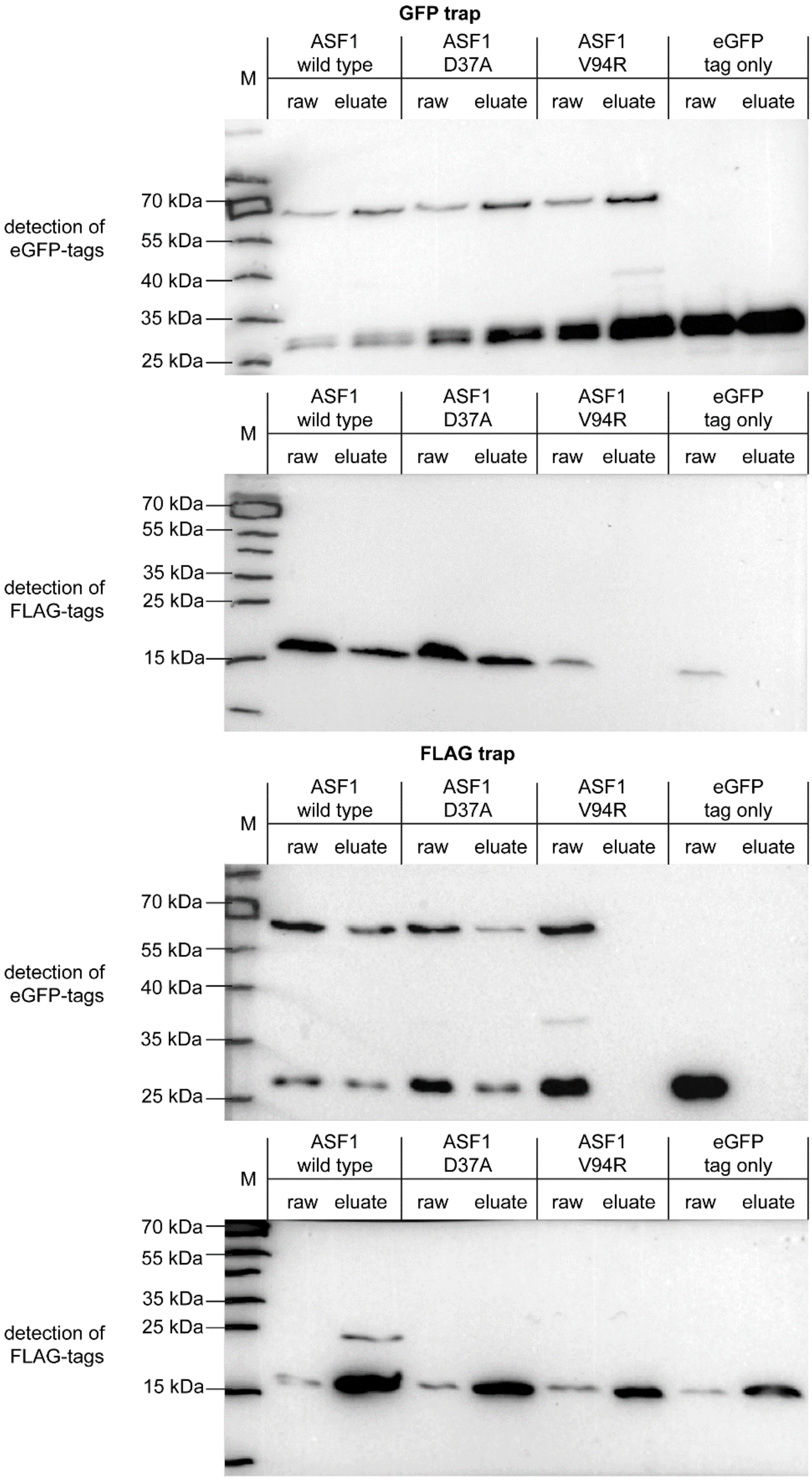


**Supplemental Figure 3B.** Uncropped blots - Co-immunoprecipitation results for ASF1 variants with amino acid substitutions and histone H4. GFP-tagged ASF1 wild type and variants D37A and V94R were used as potential interaction partners for Flag-tagged histone H4 in a GFP trap and a Flag trap. Results were checked by western blot analysis with antibodies against GFP and Flag tags. ASF1 wild type and the D37A variant showed signals for bait and prey proteins in the raw and eluate sample, indicating interaction, whereas the V94R variant showed the signal for the prey protein only in the raw sample. Strains expressing non-fused GFP with the corresponding Flag-tagged H4 were used as a negative control.

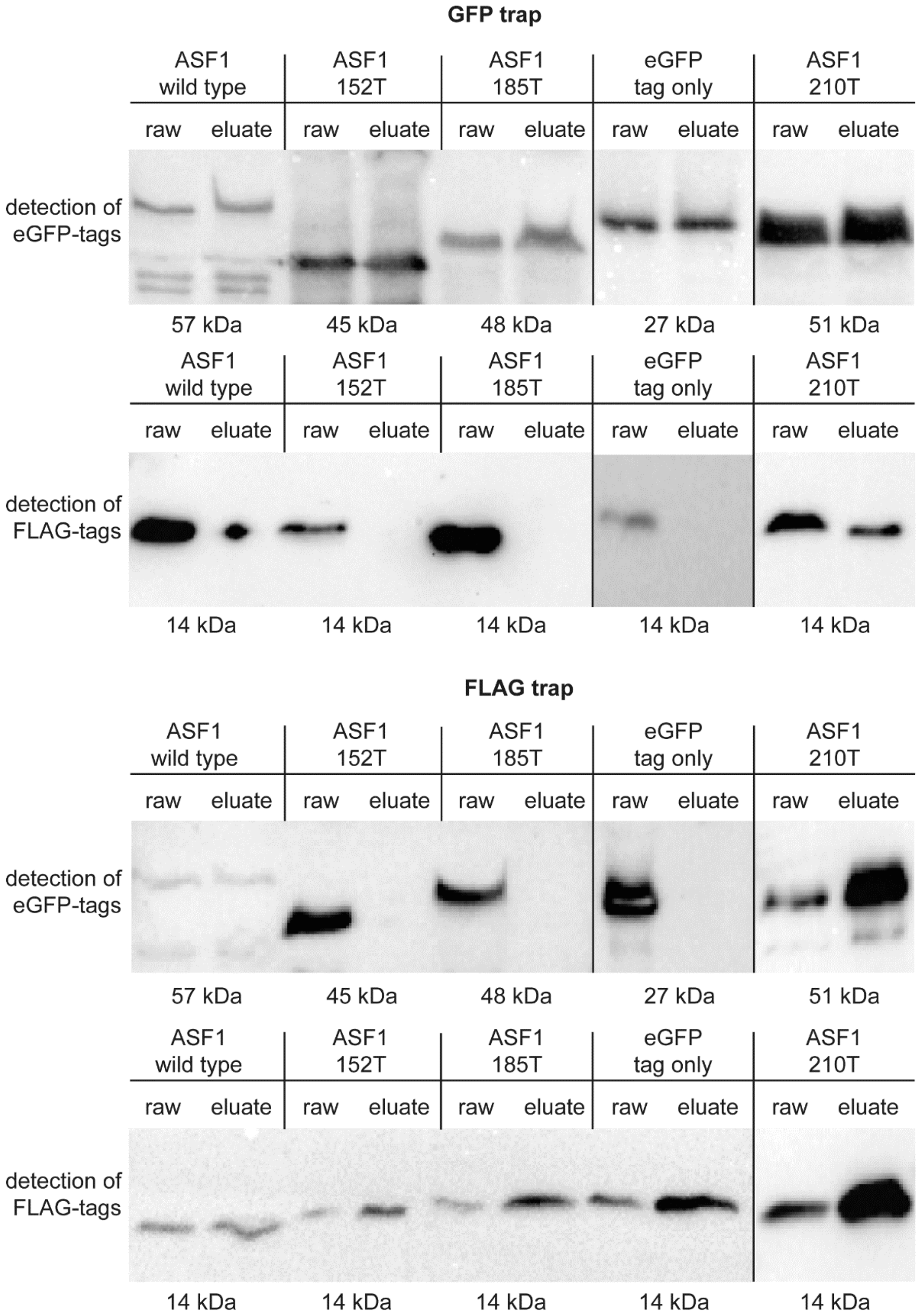


**Supplemental Figure 4.** Co-immunoprecipitation results for truncated ASF1 variants and histone H4. GFP-tagged ASF1 wild type and variants 152T, 185T and 210T were used as potential interaction partners for Flag-tagged histone H4 in a GFP trap and a Flag trap. Results were checked by western blot analysis with antibodies against GFP and Flag tags. ASF1 wild type and the 210T variant showed signals for bait and prey proteins in the raw and eluate sample, indicating interaction, whereas the 152T and 185T variants showed the signal for the prey protein only in the raw sample. Strains expressing non-fused GFP with the corresponding Flag-tagged H4 were used as a negative control. Uncropped blots are shown in Supplemental Figure 5B.

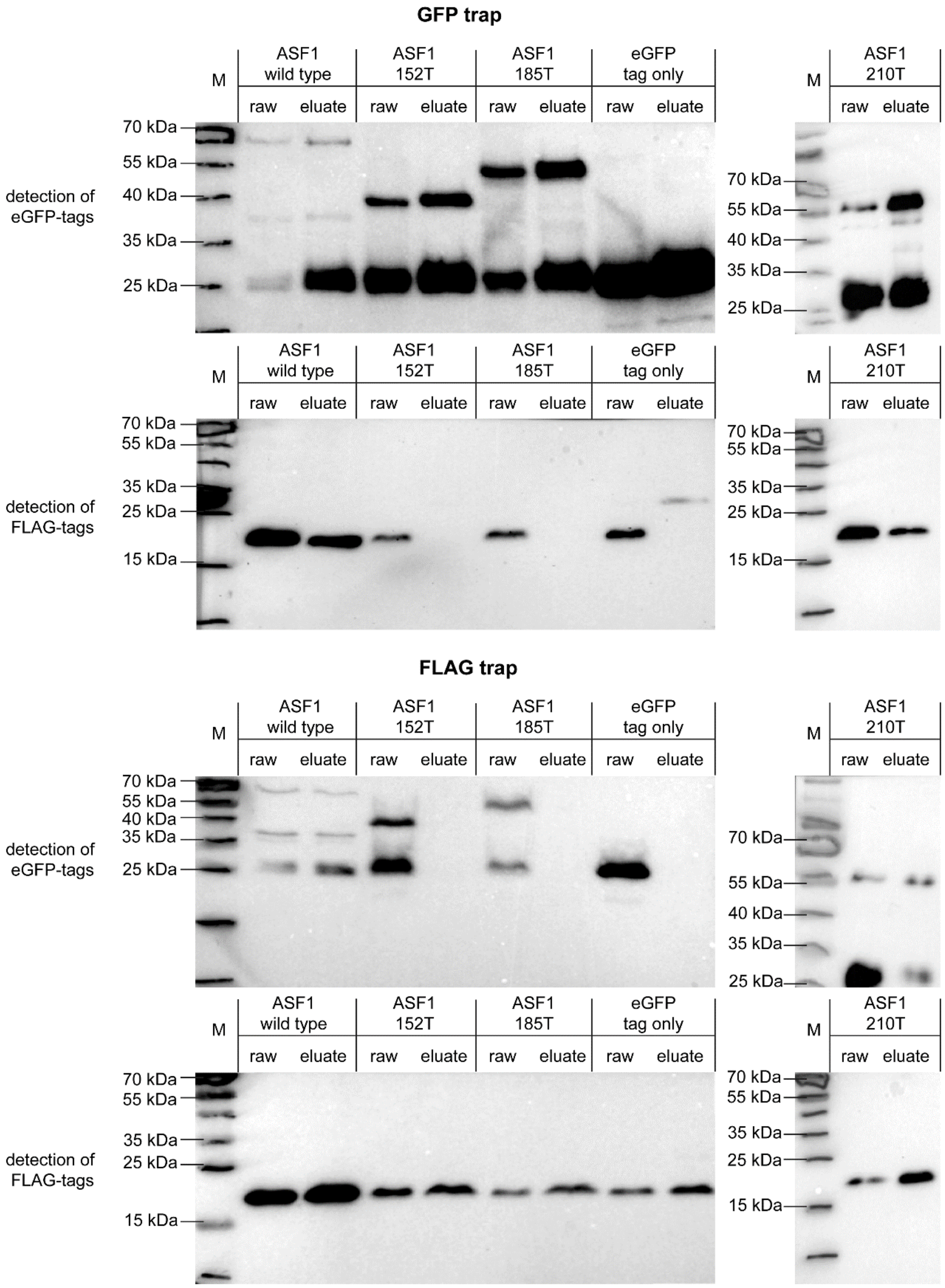


**Supplemental Figure 5A**. Uncropped blots - Co-immunoprecipitation results for truncated ASF1 variants and histone H3. GFP-tagged ASF1 wild type and variants 152T, 185T and 210T were used as potential interaction partners for Flag-tagged histone H3 in a GFP trap and a Flag trap. Results were checked by western blot analysis with antibodies against GFP and Flag tags. ASF1 wild type and the 210T variant showed signals for bait and prey proteins in the raw and eluate sample, indicating interaction, whereas the 152T and 185T variants showed the signal for the prey protein only in the raw sample. Strains expressing non-fused GFP with the corresponding Flag-tagged H3 were used as a negative control.

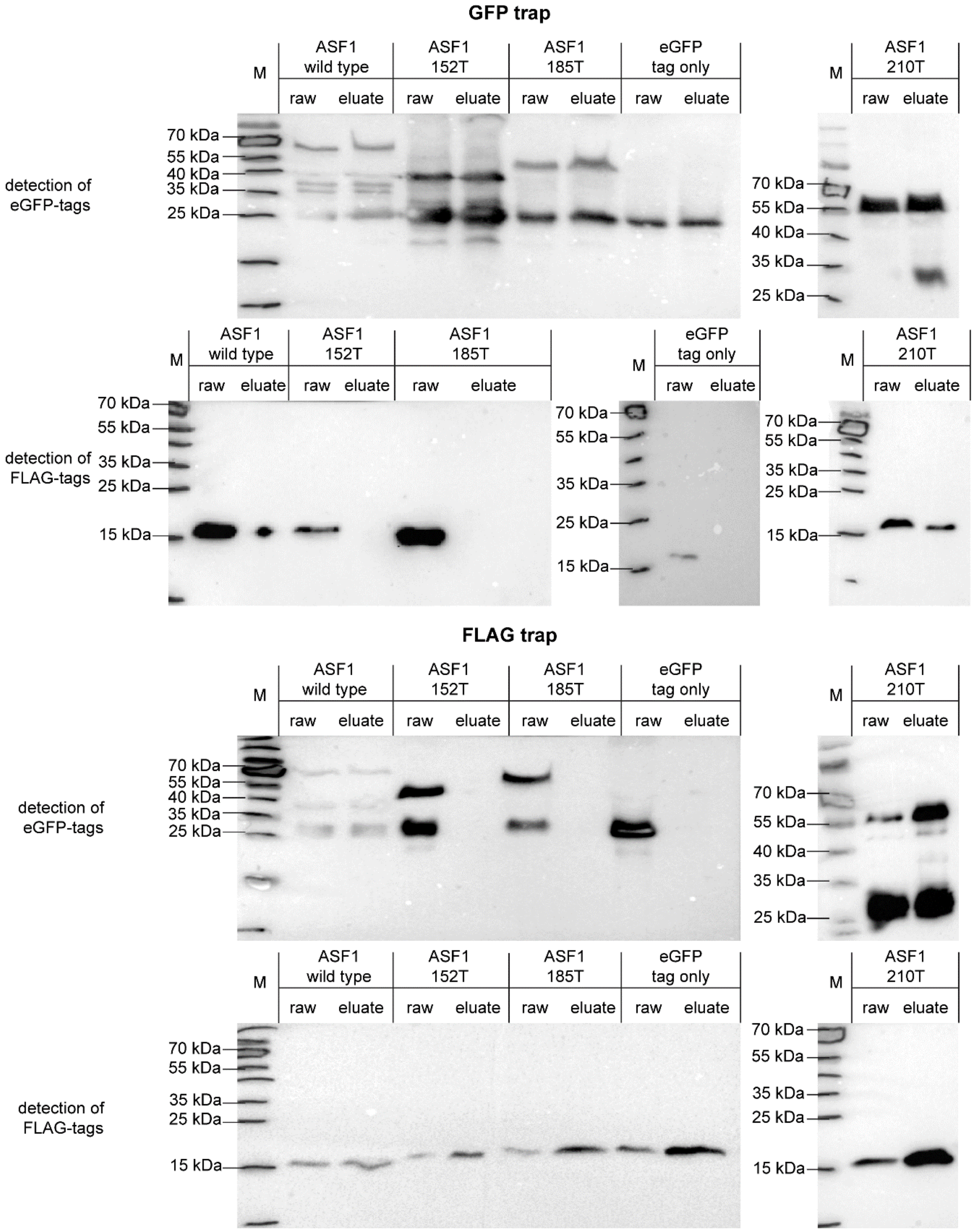


**Supplemental Figure 5B.** Uncropped blots - Co-immunoprecipitation results for truncated ASF1 variants and histone H4. GFP-tagged ASF1 wild type and variants 152T, 185T and 210T were used as potential interaction partners for Flag-tagged histone H4 in a GFP trap and a Flag trap. Results were checked by western blot analysis with antibodies against GFP and Flag tags. ASF1 wild type and the 210T variant showed signals for bait and prey proteins in the raw and eluate sample, indicating interaction, whereas the 152T and 185T variants showed the signal for the prey protein only in the raw sample. Strains expressing non-fused GFP with the corresponding Flag-tagged H4 were used as a negative control.

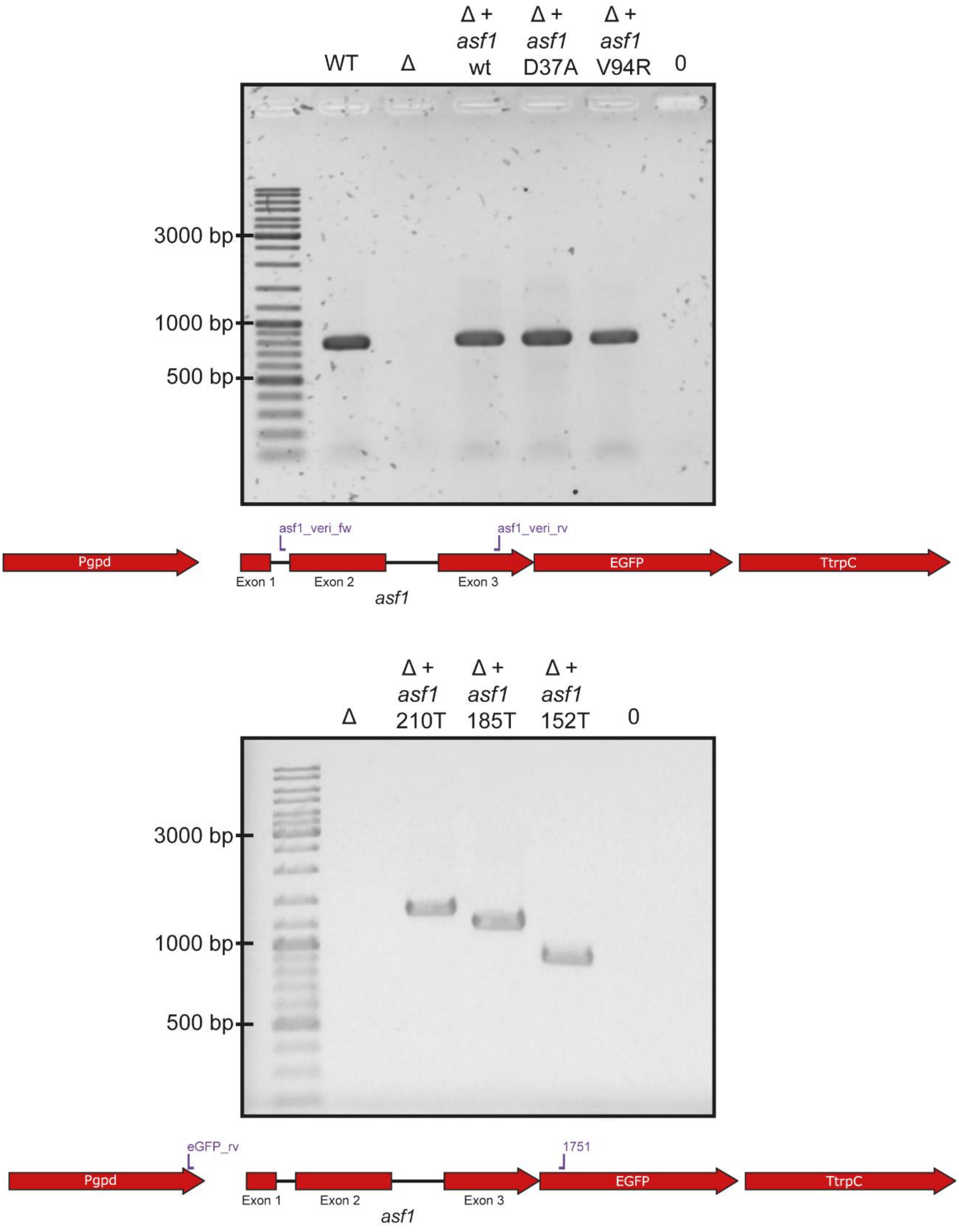


**Supplemental Figure 6.** PCR verification of transformation of Δasf1 with *asf1* variants. Primers asf1_veri_fw and asf1_very_rv were used to confirm the integration of the *asf1* wild type, D37A and V94R constructs, amplifying a 786 bp region within the *asf1* gene, which is not present in the deletion mutant. Primers 1751 and eGFP_rv were used to amplify parts of the integrated complementation vectors for *asf1* 152T, 185T and 210T variants, allowing to also verify the integration of the correct length of the truncated variants. The expected bands of 1282 bp for *asf1* 210T, 1207 bp for *asf1* 185T and 918 bp for *asf1* 152T were detected. Abbreviation: Δ = Δasf1, 0 = no template control.

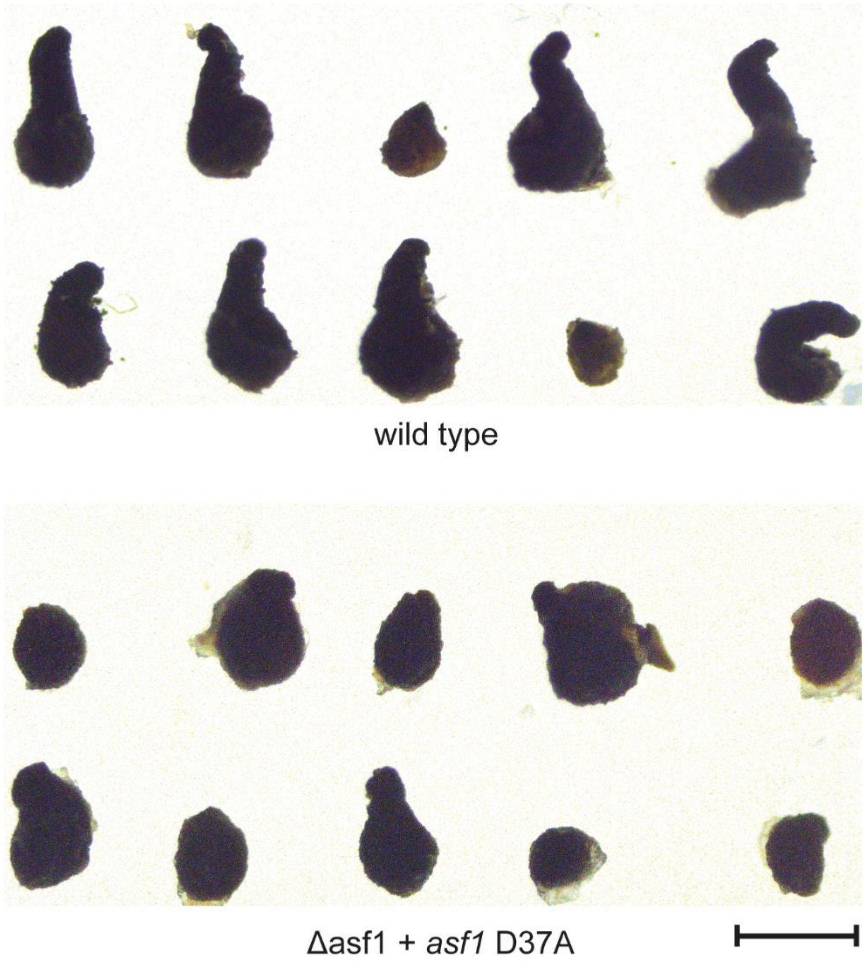


**Supplemental Figure 7.** Comparison of fruiting body morphology of wild type and Δasf1 + asf1 D37A. While the wild type produces fully grown, cone-shaped perithecia as well as protoperithecia, strains expressing asf1 with a D37A substitution tend to produce perithecia that appear more rounded and look like giant protoperithecia, although the production of normal perithecia is generally possible. The scale bar represents 300 µm.

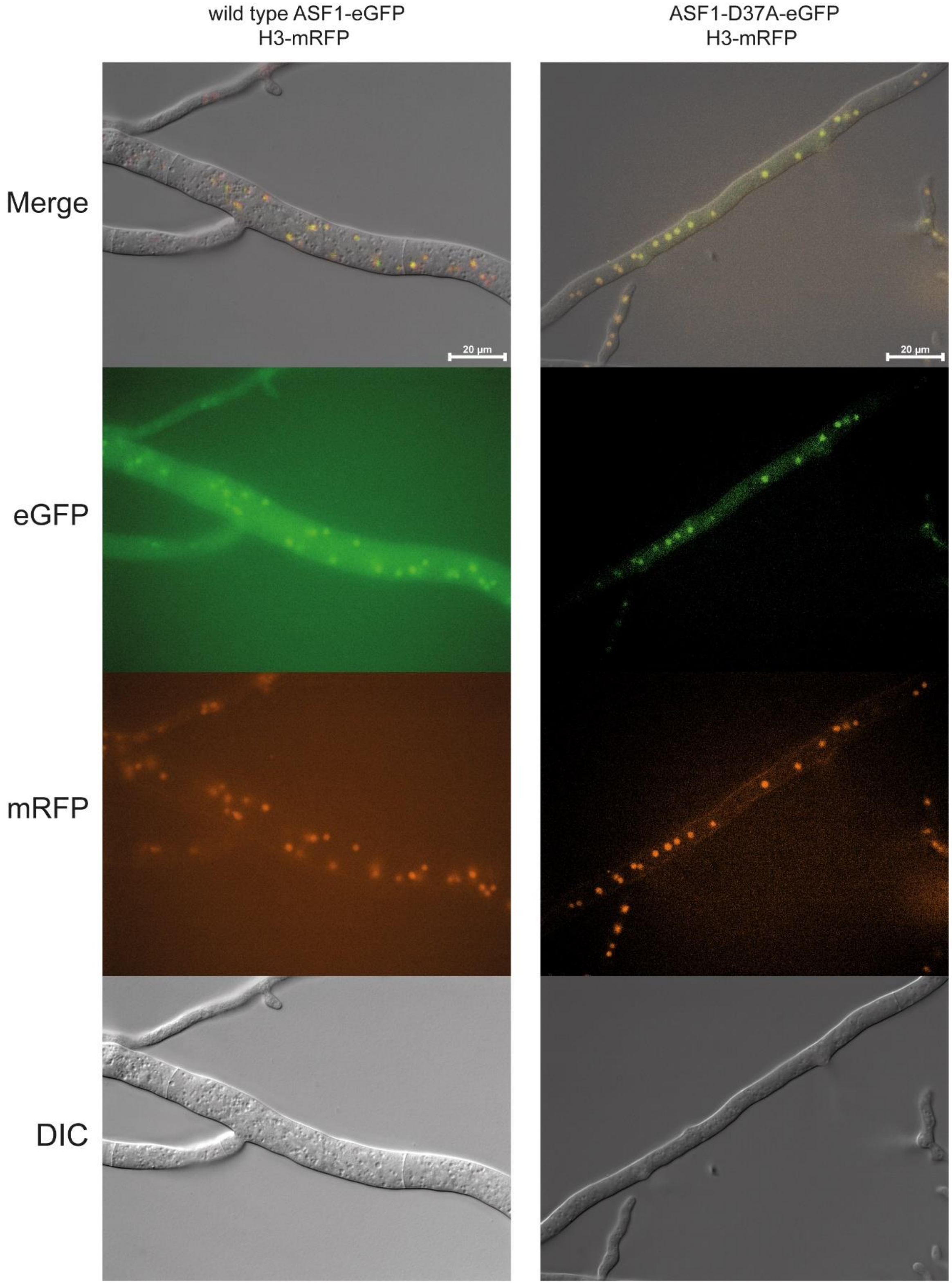


**Supplemental Figure 8A.** Localization of ASF1-variants by fluorescence microscopy. To confirm the correct localization of ASF1 variants in the nucleus, eGFP tagged variants were grown together with a control strain expressing histone H3 with an mRFP tag. Hyphal fusion of both strains leads to nuclear exchange and therefore hyphae that contain both tagged proteins. Both tags colocalized, detectable in the shown overlay images (GFP fluorescence, mRFP fluorescence and differential interference contrast to show hyphal outlines) by yellow - orange fluorescence, thus confirming the nuclear localization of all ASF1 variants.

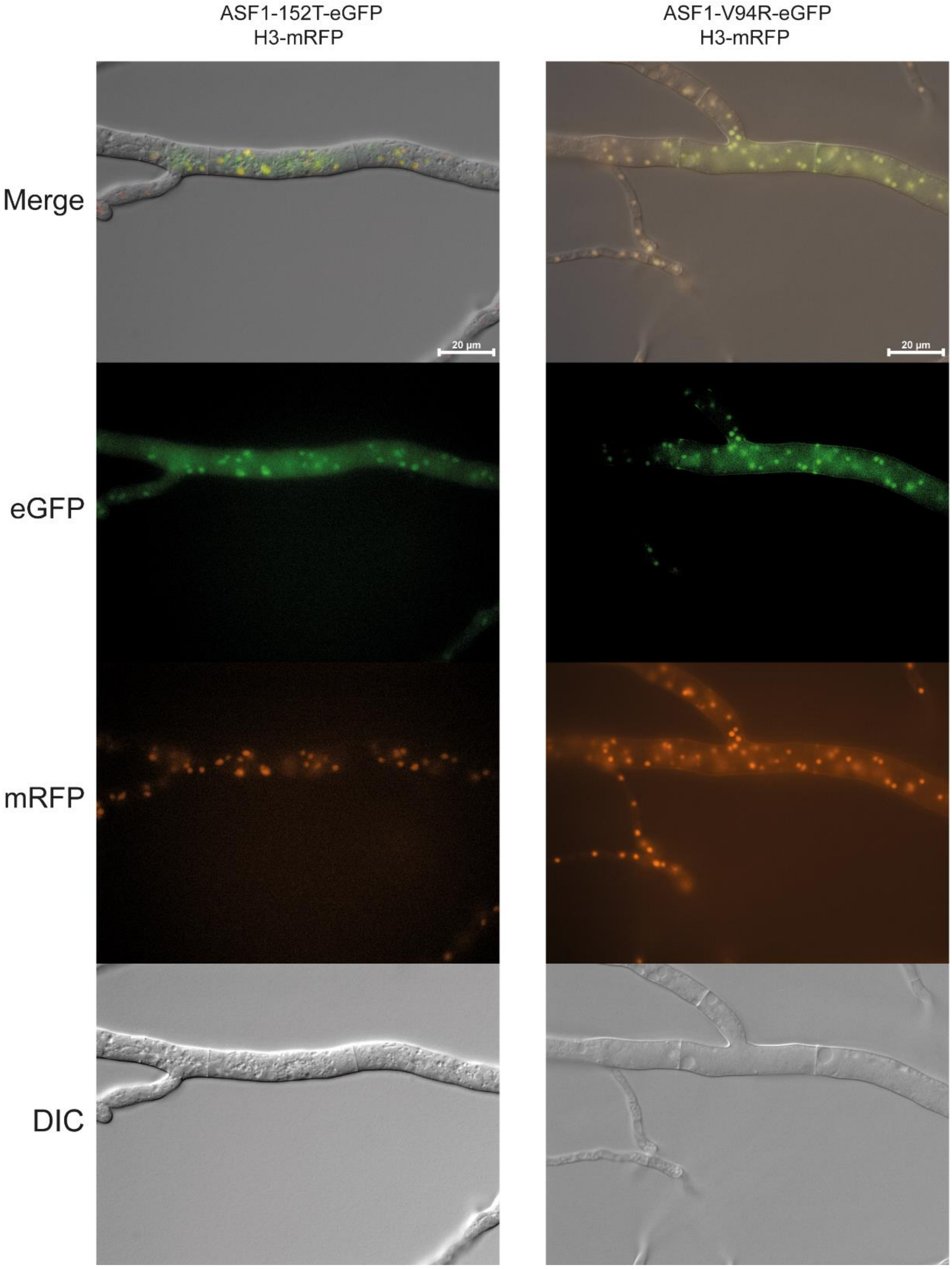


**Supplemental Figure 8B.** Localization of ASF1-variants by fluorescence microscopy. To confirm the correct localization of ASF1 variants in the nucleus, eGFP tagged variants were grown together with a control strain expressing histone H3 with an mRFP tag. Hyphal fusion of both strains leads to nuclear exchange and therefore hyphae that contain both tagged proteins. Both tags colocalized, detectable in the shown overlay images (GFP fluorescence, mRFP fluorescence and differential interference contrast to show hyphal outlines) by yellow - orange fluorescence, thus confirming the nuclear localization of all ASF1 variants.

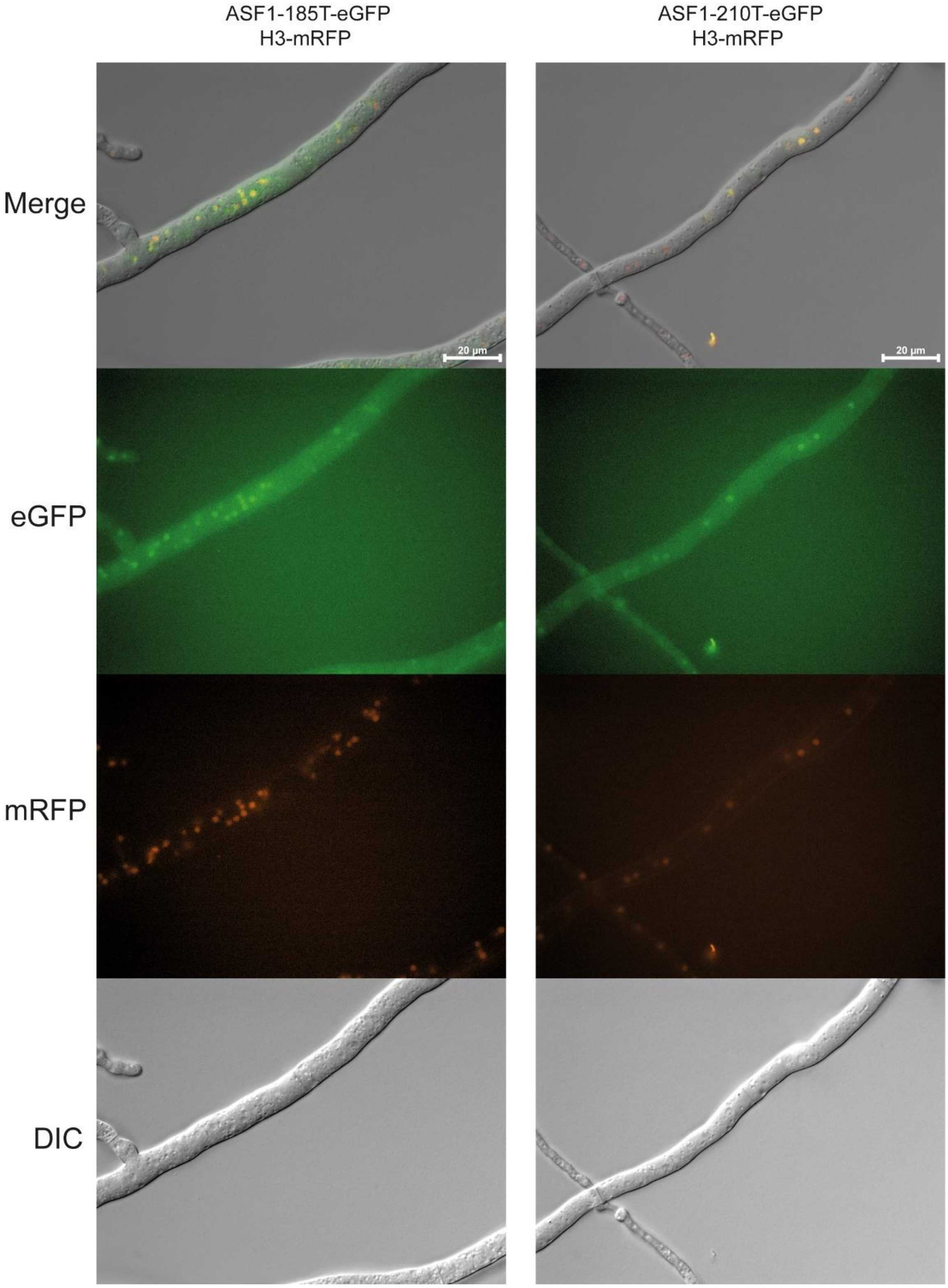


**Supplemental Figure 8C.** Localization of ASF1-variants by fluorescence microscopy. To confirm the correct localization of ASF1 variants in the nucleus, eGFP tagged variants were grown together with a control strain expressing histone H3 with an mRFP tag. Hyphal fusion of both strains leads to nuclear exchange and therefore hyphae that contain both tagged proteins. Both tags colocalized, detectable in the shown overlay images (GFP fluorescence, mRFP fluorescence and differential interference contrast to show hyphal outlines) by yellow - orange fluorescence, thus confirming the nuclear localization of all ASF1 variants.

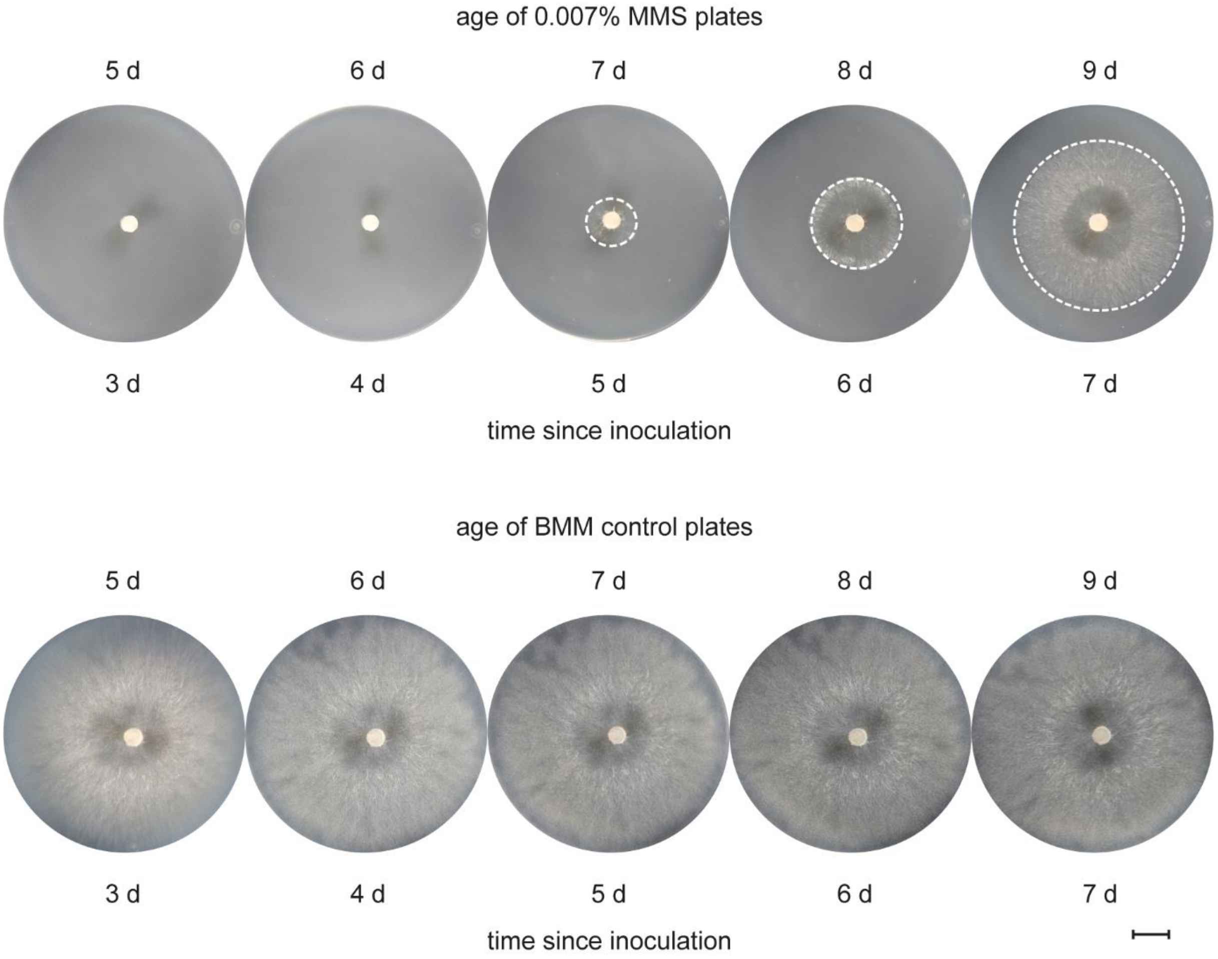


**Supplemental Figure 9.** Validation of the MMS assay. To validate the effect of the highly unstable substance MMS, the sensitive strain Δasf1 was inoculated on 5-day-old BMM plates with 0.007% MMS. While no growth was possible until day 6, small amounts of mycelia were visible by day 7. Thereafter, the strain appeared to be able to grow on the media. Freshly prepared BMM medium containing 0.007% MMS is therefore suitable for the for the 4-day observation period used in the genotoxic stress assay.

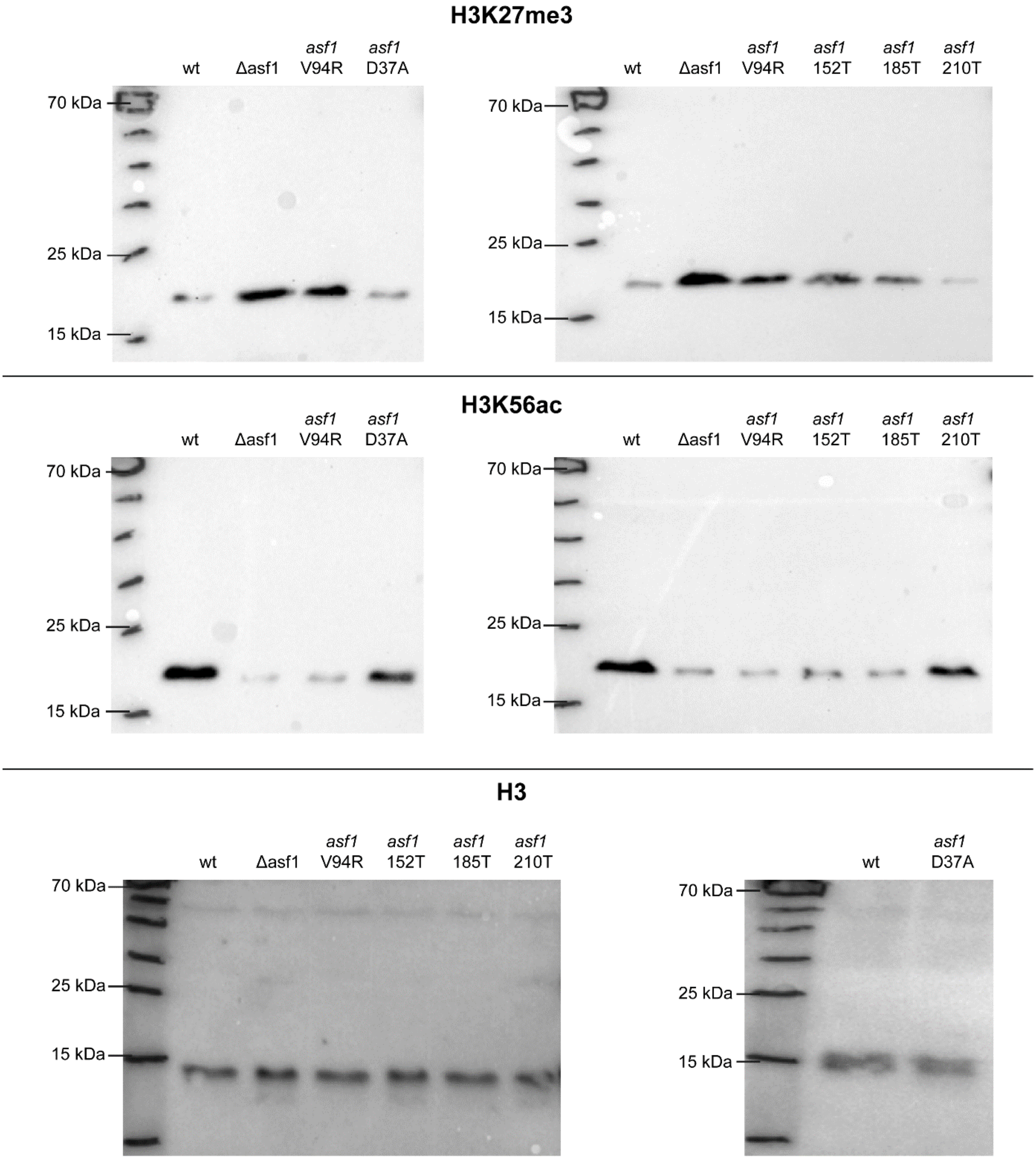


**Supplemental Figure 10.** Uncropped blots - Semiquantitative screening for histone modifications affected by ASF1. Western blots with antibodies against the indicated histone modifications were used to compare the band strength of equal amounts of protein from wild type, Δasf1 and strains expressing *asf1* variants. H3 levels were measured as an internal control.

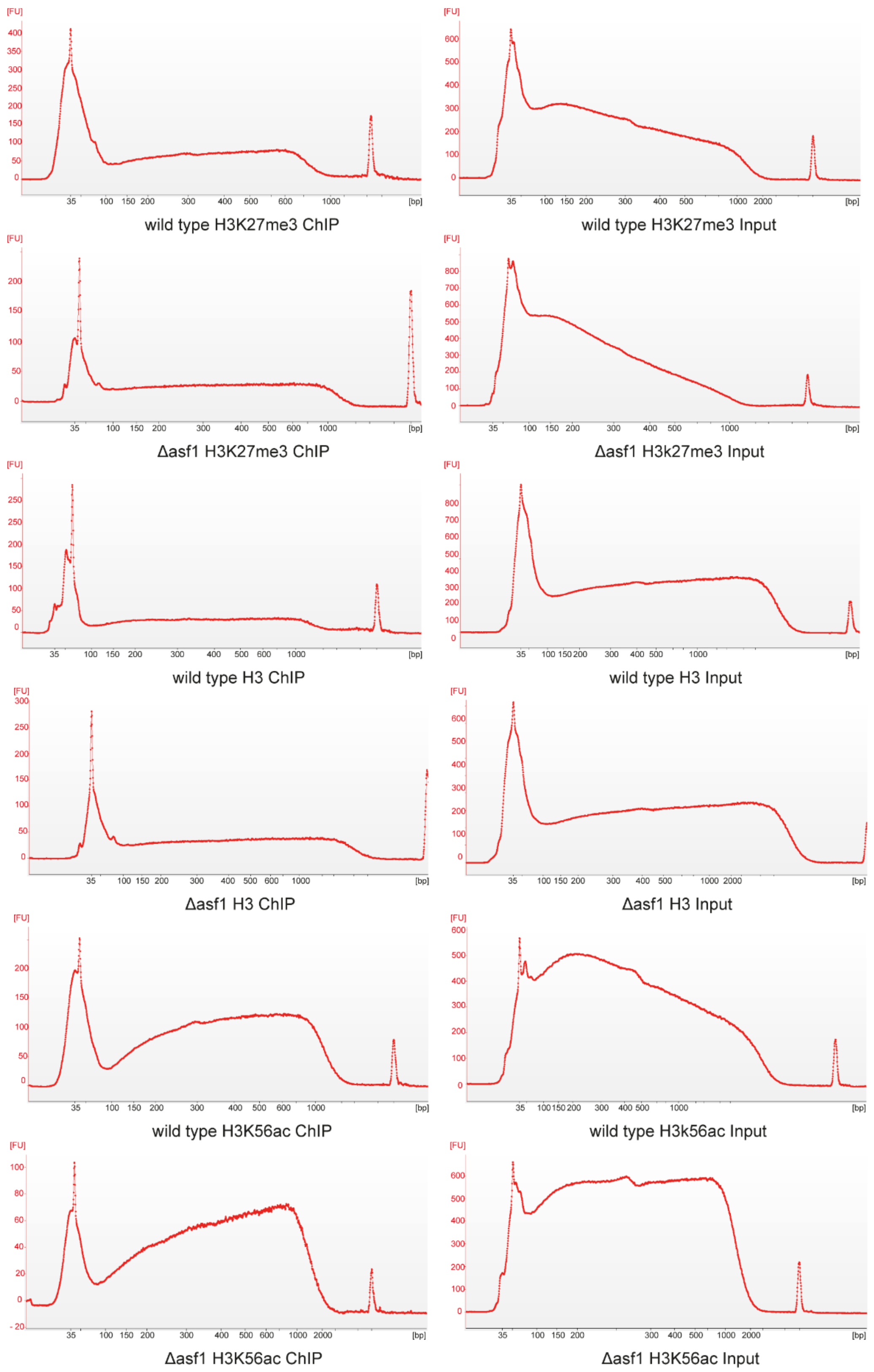


**Supplemental Figure 11.** Fragmentation of ChiP-seq samples. Samples were analyzed on an Agilent Bioanalyzer 2100 with High Sensitivity DNA Kit. If more than 20 ng were detected in the desired range of 150 to 500 bp, samples were sent to Novogene for size selection, library generation, and sequencing. Y-Axis = fluorescence level, X-Axis = basepairs.

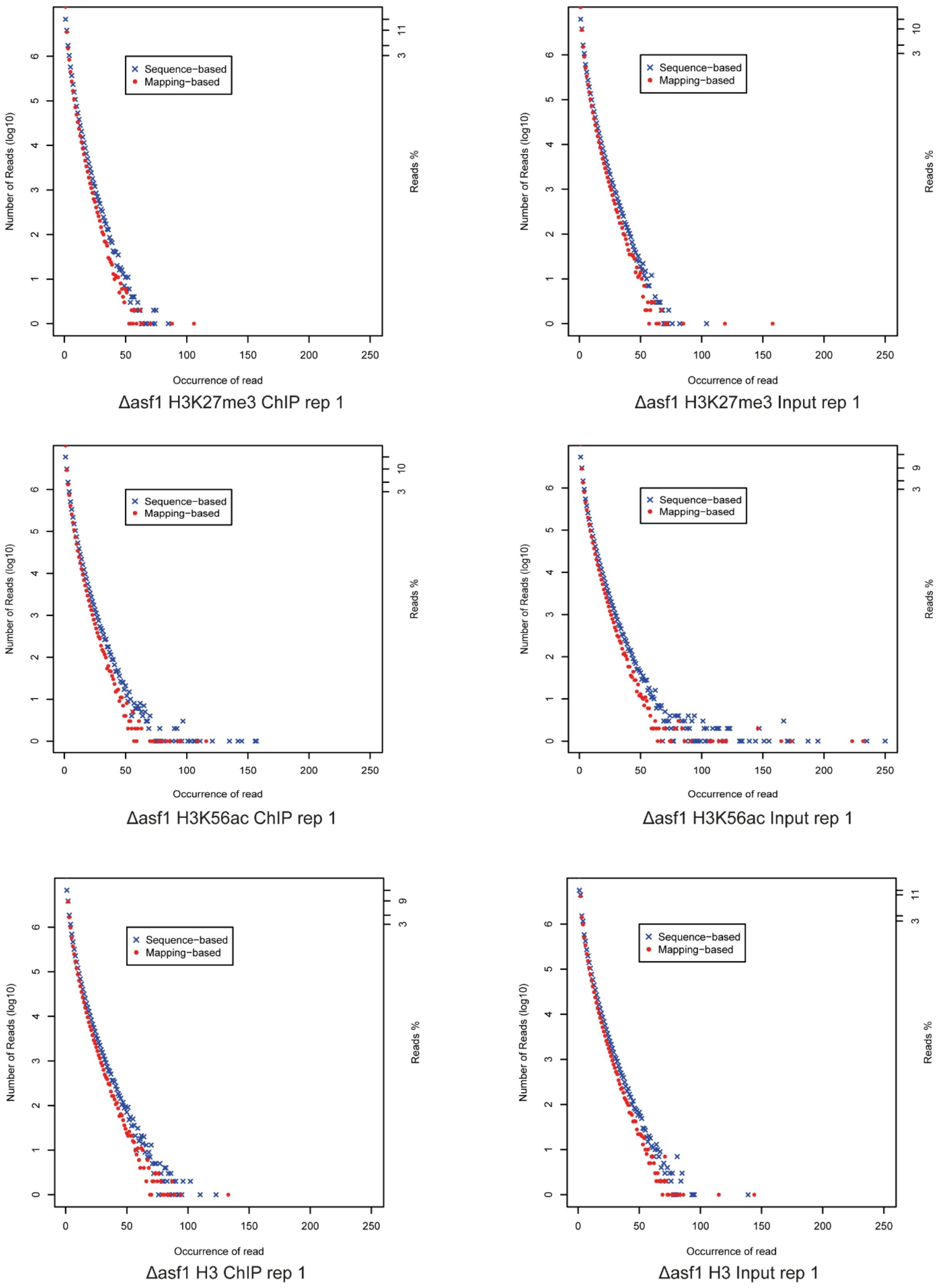


**Supplemental Figure 12A.** Duplication rates of ChIP-seq Δasf1 replicate 1 samples before deduplication. After quality control and trimming, reads were mapped to the reference genome and checked for sequence- and mapping-based duplication rates. For peak detection, all datasets were deduplicated using MACS2.

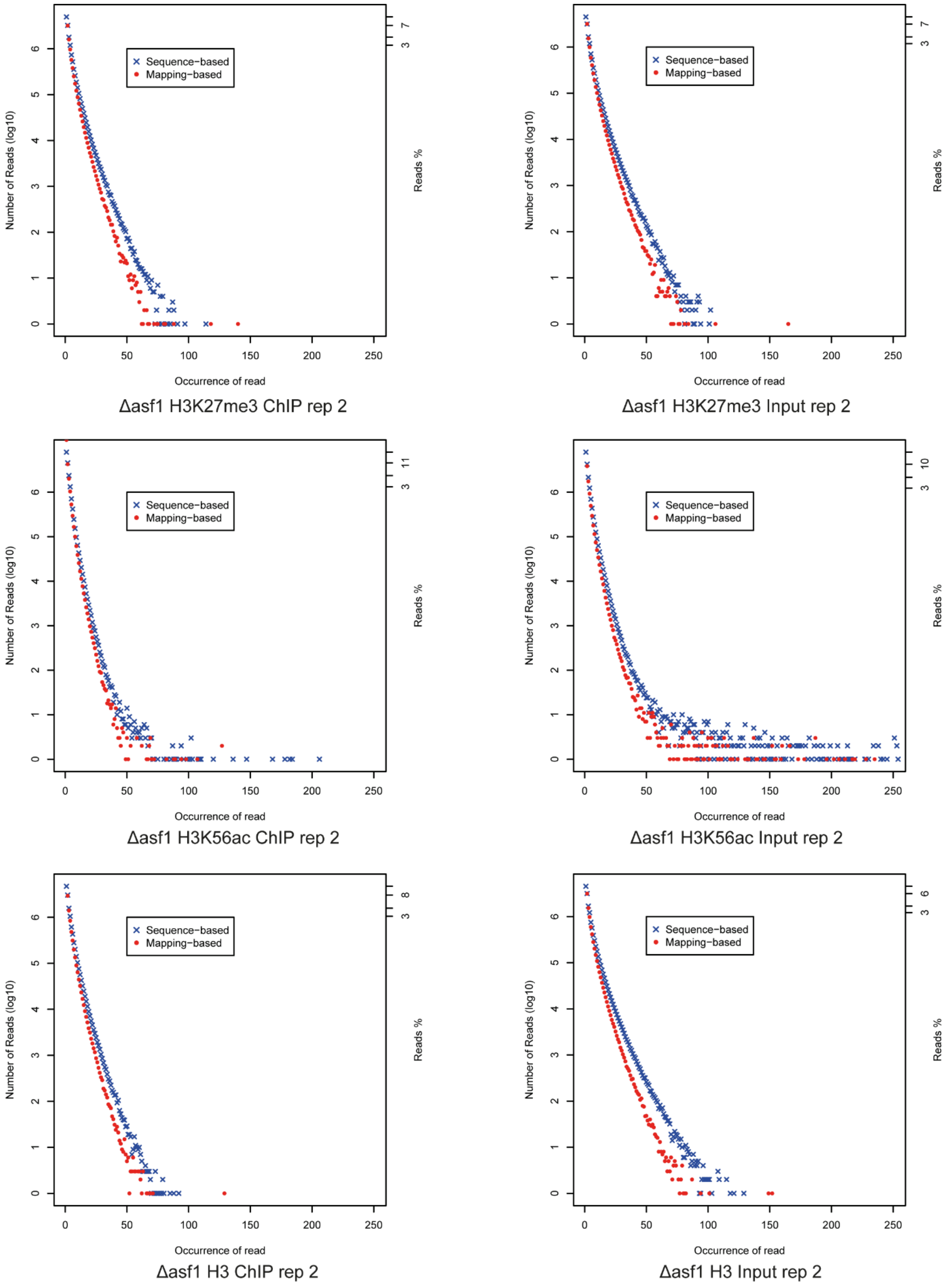


**Supplemental Figure 12B.** Duplication rates of ChIP-seq Δasf1 replicate 2 samples before deduplication. After quality control and trimming, reads were mapped to the reference genome and checked for sequence- and mapping-based duplication rates. For peak detection, all datasets were deduplicated using MACS2.

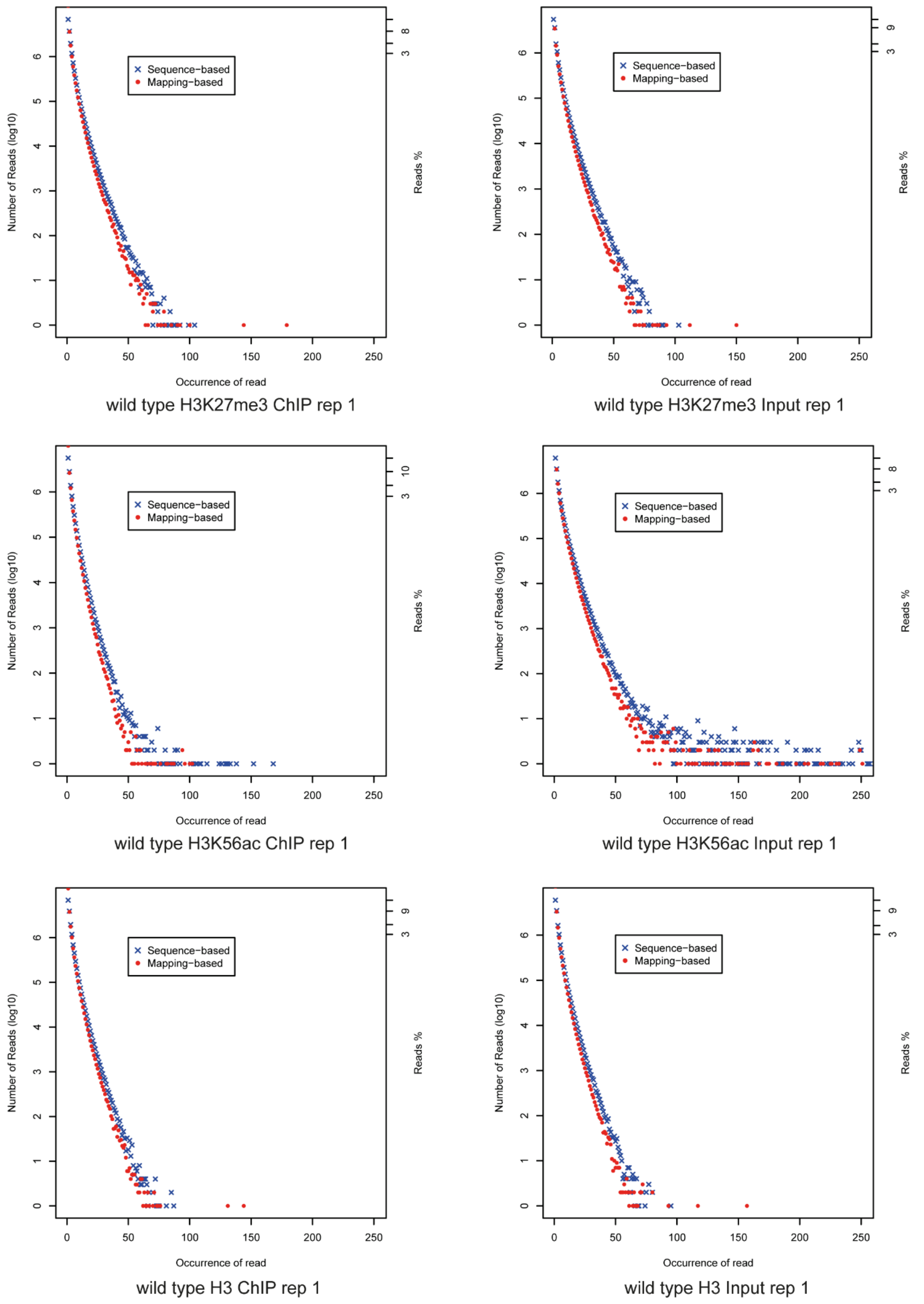


**Supplemental Figure 12C.** Duplication rates of ChIP-seq wild type replicate 1 samples before deduplication. After quality control and trimming, reads were mapped to the reference genome and checked for sequence- and mapping-based duplication rates. For peak detection, all datasets were deduplicated using MACS2.

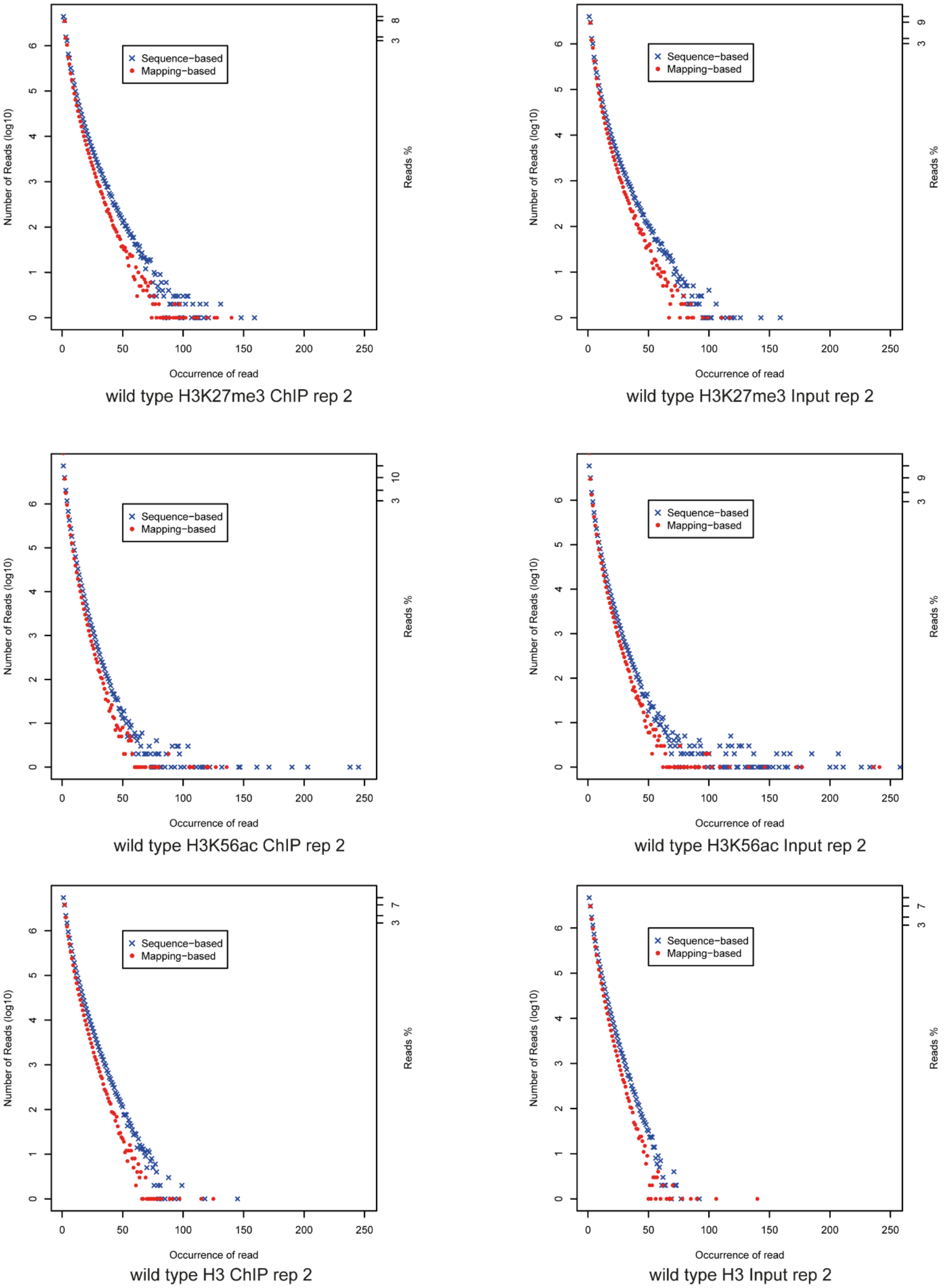


**Supplemental Figure 12D.** Duplication rates of ChIP-seq wild type replicate 2 samples before deduplication. After quality control and trimming, reads were mapped to the reference genome and checked for sequence- and mapping-based duplication rates. For peak detection, all datasets were deduplicated using MACS2.

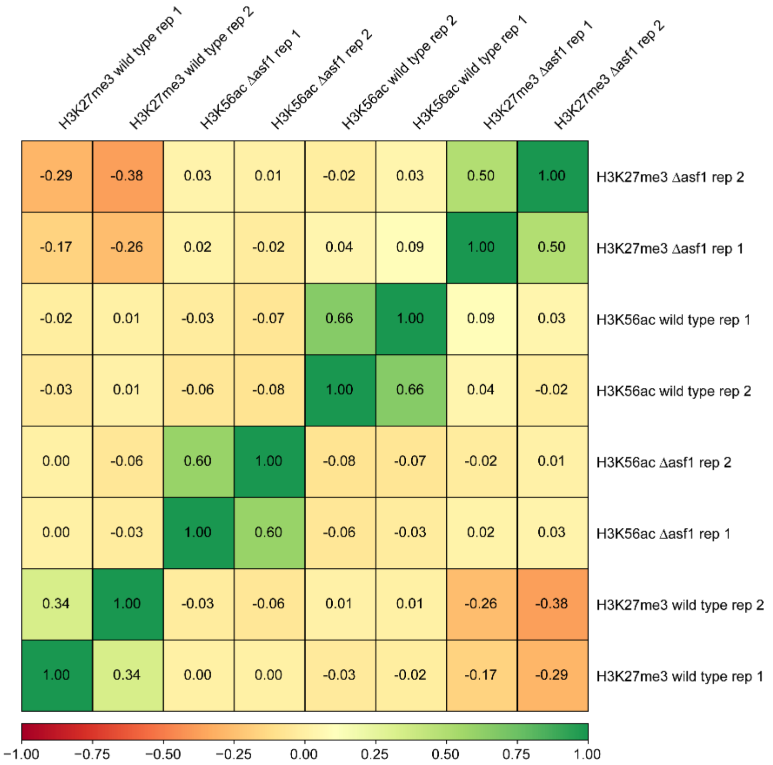


**Supplemental Figure 13.** Pearson correlation of ChIP-seq samples. Positive correlations were detected for the respective replicates of ChIP samples.

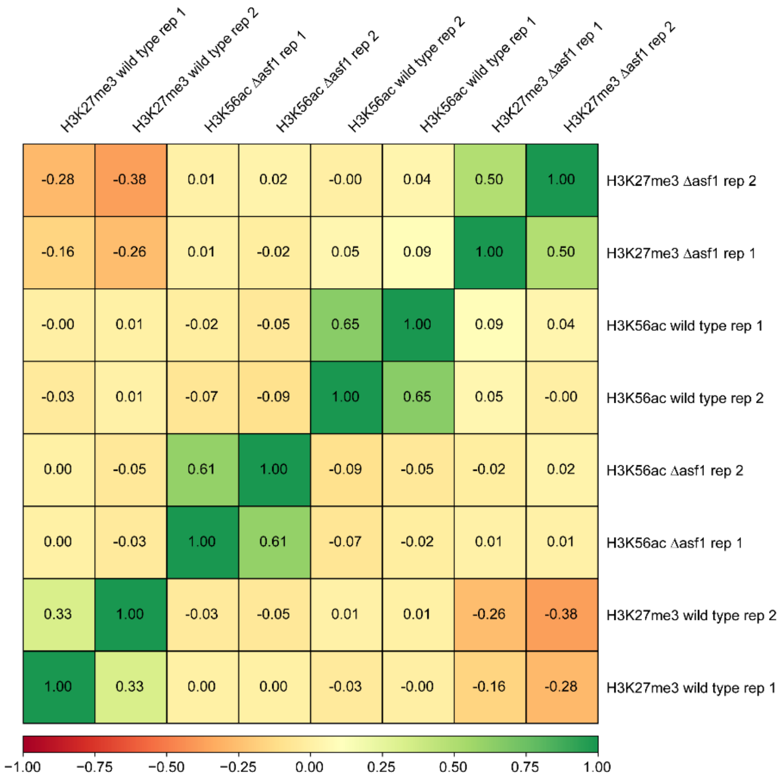


**Supplemental Figure 14.** Spearman correlation of ChIP-seq samples. Positive correlations were detected for the respective replicates of ChIP samples.

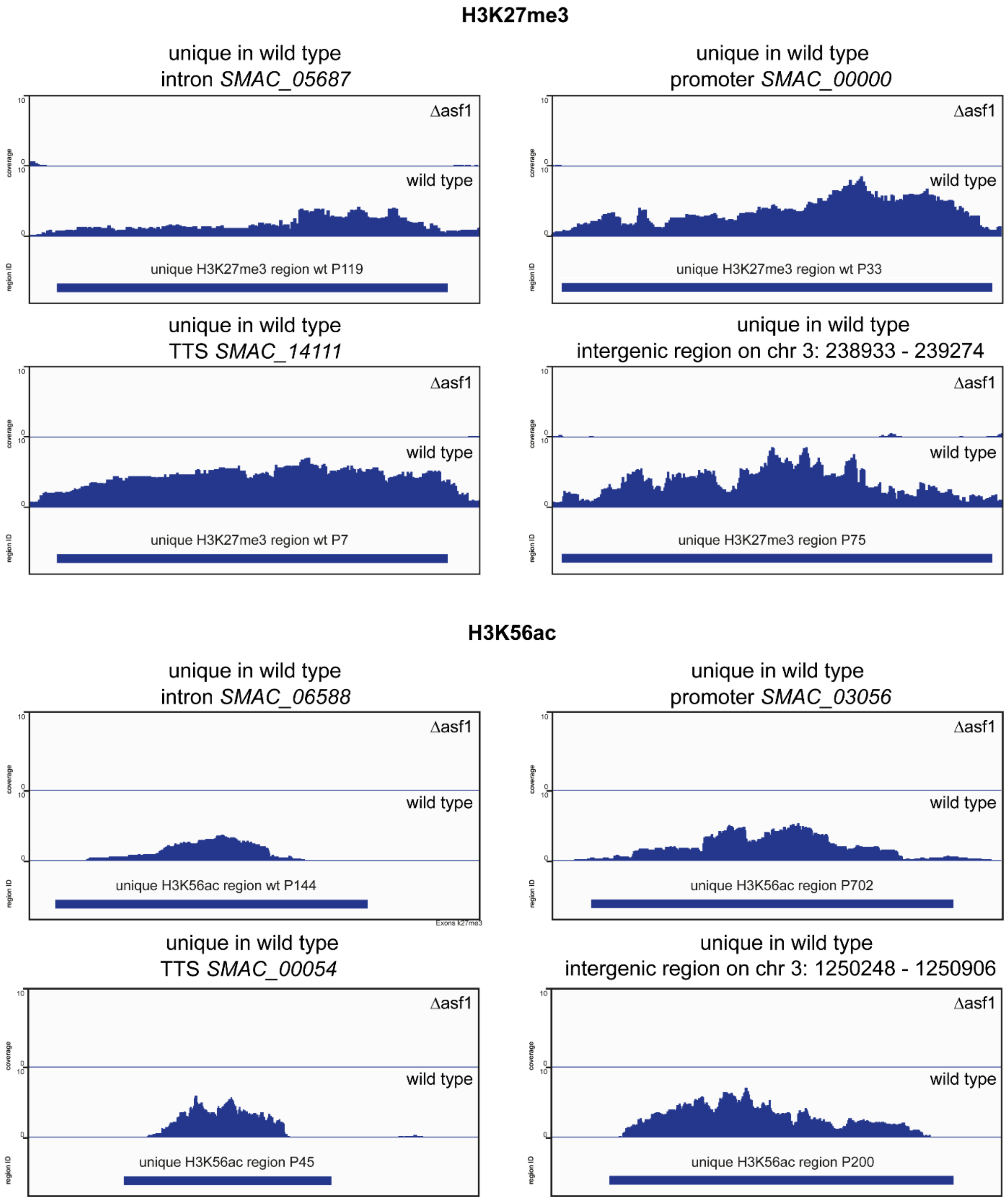


**Supplemental Figure 15A.** Representative regions exhibiting H3K27me3 and H3K56ac uniquely in the wild type. The coverage in the wild type and the deletion mutant is shown alongside the respective region.

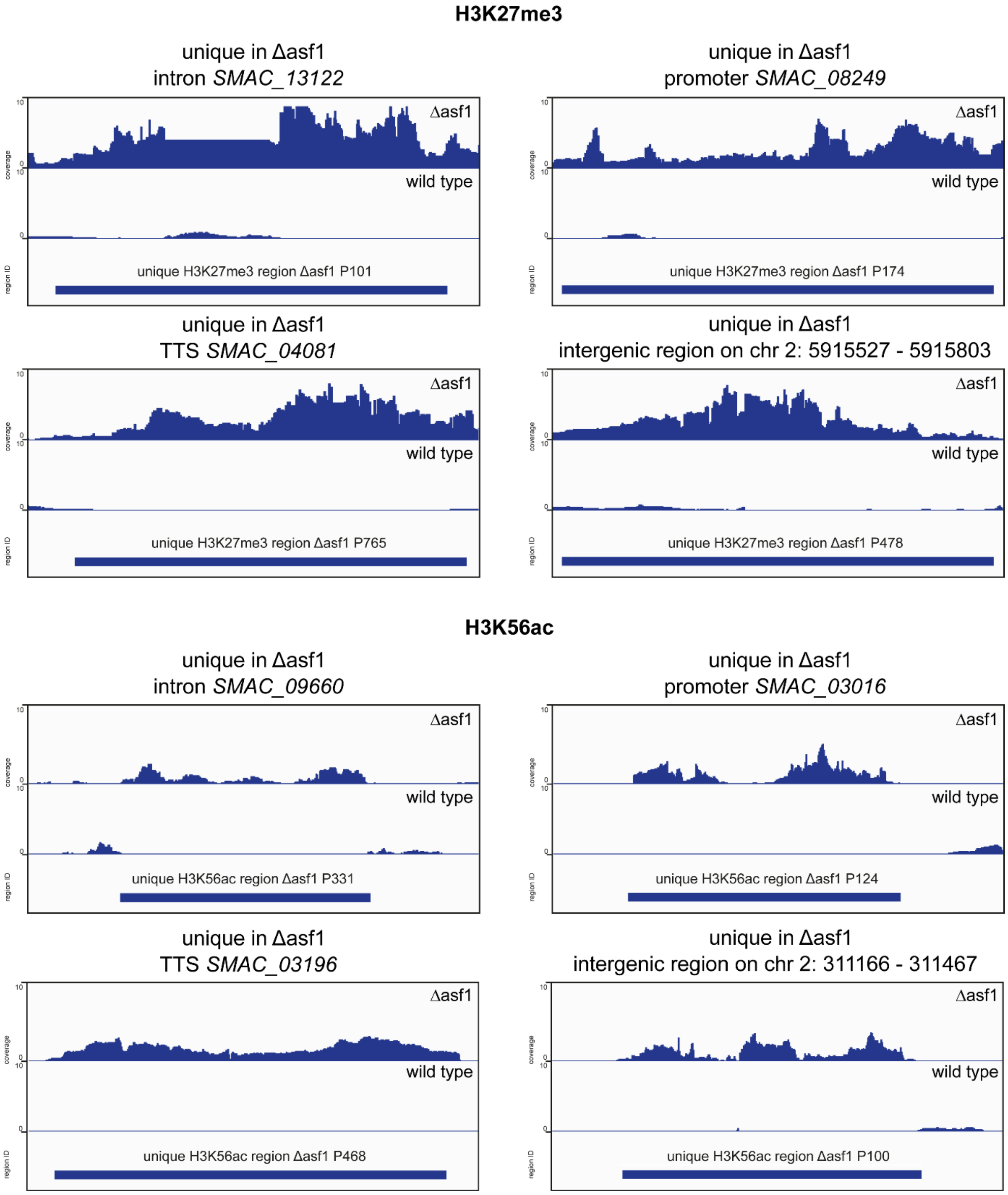


**Supplemental Figure 15B.** Representative regions exhibiting H3K27me3 and H3K56ac uniquely in the Δasf1 strain. The coverage in the wild type and the deletion mutant is shown alongside the respective region.

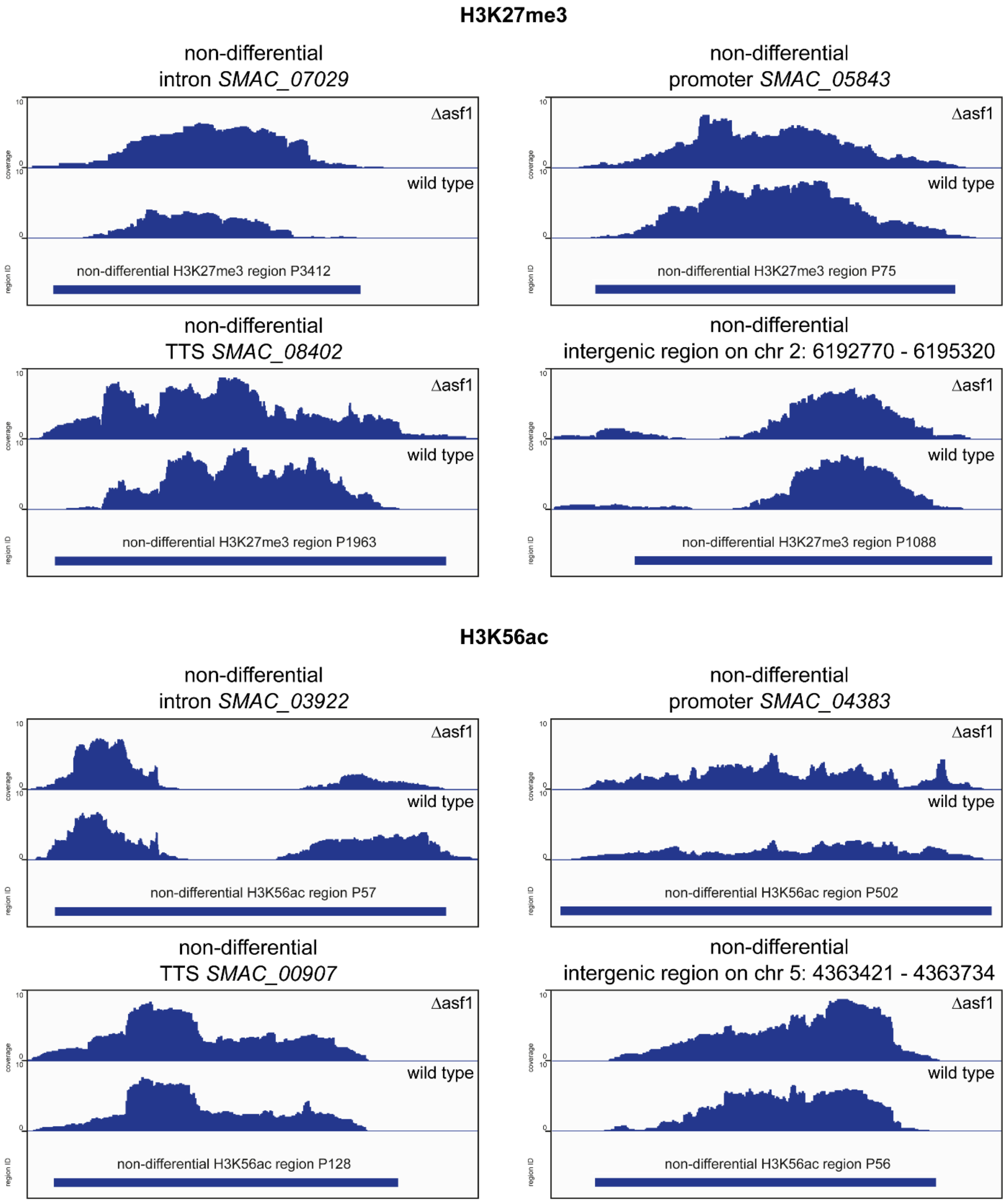


**Supplemental Figure 15C.** Representative regions exhibiting H3K27me3 and H3K56ac in the wild type and Δasf1 strain. The coverage in the wild type and the deletion mutant is shown alongside the respective region. TTS indicates a transcription termination region.

**Supplemental Text 1.** ChIP-Seq protocol used in this study.

4 Fernbach flasks with 150 ml of CM-media each were inoculated with *S. macrospora* wild type or Δasf1 and cultivated for 3 or 4 days at 27 °C. Mycelium was harvested by filtration, washed with PPP twice and frozen in liquid nitrogen. Frozen samples were ground to a fine powder under regular addition of liquid nitrogen for at least 10 min. The powder was transferred to a 50 ml falcon tube until the 30 ml mark was reached. For crosslinking, the powder was mixed with 30 ml crosslinking solution (PPP + 1 % formaldehyde) and placed on a rotating wheel for 15 min at room temperature, followed by a quenching step with 3 ml of 2.5 M glycine for 5 min. The samples were centrifugated at 2500 rpm for 5 min at 4 °C and the supernatant was discarded. The pellets containing the crosslinked samples were washed twice with PPP. Lysis was performed by adding 5 ml lysis buffer with protease inhibitors (0.3 % (v / v) Protease Inhibitor Cocktail Set IV (Calbiochem) + 100 µl PMSF) and two steel mixing balls. Samples were kept on ice for 15 min and mixed regular by intense vortexing. To obtain a chromatin solution, centrifugation was performed at 6000 rpm at 4 °C for 5 min. The supernatant was kept and transferred to multiple 2 ml Eppendorf tubes and centrifugated again at 13000 rpm at 4 °C for 5 min to get rid of cell debris and obtain a clear solution. 4 ml of the solution were transferred to a 15 ml falcon tube and mixed with 2 ml MNase buffer. Chromatin digestion was conducted by using 200 U/ml MNase (Thermo Fisher Scientific) and separating the samples in 6 batches of 1 ml in 1,5 ml Eppendorf tubes. Tubes were incubated for 105 min at 37 °C and 800 rpm in a thermomixer. The reaction was stopped by adding EDTA to a concentration of 20 mM. To confirm correct digestion, one 1 ml sample was de-crosslinked by adding SDS up to a final concentration of 1 % and 2 mg/ml proteinase K (Genaxxon bioscience) and incubation at 65 °C over night, while the other samples were frozen at -80 °C for later use. The de-crosslinked sample was used for phenol/chloroform extraction and separated on a 1 % agarose gel. The desired digestion profile should be visible by a smear that appears strongest between 500 and 100 bp. After confirmation of correct digestion, 10 µl of the respective antibody (Anti H3K27Me3 #9733 Cell Signaling, Anti H3 #2650 Cell Signaling, Anti H3K56Ac #61061 Active Motif) were added to 5 Eppendorf tubes per sample, each containing 1 ml of digested chromatin solution. The samples were incubated over night at 4 °C on a spinning wheel. Afterwards, 4 h of incubation with 50 µl Protein-A agarose beads (Santa Cruz) per tube were followed by 2 washing steps in low salt and 2 washing steps in high salt buffer. Each washing step was performed for 5 min at 4 °C on a spinning wheel and followed by 2 min of centrifugation at 2500 rpm to separate the beads from the buffer. Chromatin was washed off the beads by 2 h of incubation in TE buffer with 1 % SDS at 65 °C. The tubes containing the samples were centrifuged for 5 min at 6000 rpm to collect the supernatant that was merged in a 15 ml falcon tube and de-crosslinked over night at 65 °C after adding 2 mg/ml proteinase K. Phenol/chloroform extraction was performed to isolate DNA from the samples. Final samples were

checked on a Bioanalyzer 2100 (Agilent) with the Agilent High Sensitivity DNA Kit. A clear accumulation of fragments between 100 and 500 bp should be detectable to qualify for sequencing.

Buffers and media used in the ChIP-Seq protocol:

CM medium: 1 % (w/v) glucose, 0.2 % (w/v) tryptone, 0.2 % (w/v) yeast extract, 0.15 % (w/v) $KH_2PO_4$, 0.05 % (w/v) KCl, 0.05 % (w/v) $MgSO_4$, 0.37 % (w/v) $NH_4Cl$, 0.01 % (w/v) $ZnSO_4$, 0.01 % (w/v) $Fe(II)Cl_2$, 0.01 % (w/v) $MnCl_2$, pH 6.5 (KOH)

PPP: 13 mM $Na_2HPO_4$, 45 mM $KH_2PO_4$, 600 mM KCL, pH 6.0 (KOH)

Lysis buffer: 50 mM Tris/HCl pH 7.5, 250 mM NaCl, 0.05 % NP-40, 0.05 % β-mercaptoethanol

MNase Buffer: 5 mM $CaCl_2$, 50 ml Tris-HCl, pH 8

Low salt washing buffer: 50 mM Tris-HCL pH 7.5, 10 mM EDTA, 10 mM NaCl

High Salt washing buffer: 50 mM Tris-HCL pH 7.5, 10 mM EDTA, 500 mM NaCl

TE buffer: 10 mM Tris-HCL, 1 mM EDTA
